# The serine/threonine kinase MINK1 directly regulates the function of promigratory proteins

**DOI:** 10.1101/2021.09.02.458712

**Authors:** Avais M. Daulat, Mônica S. Wagner, Stéphane Audebert, Malgorzata Kowalczewska, Jeremy Ariey-Bonnet, Pascal Finetti, François Bertucci, Luc Camoin, Jean-Paul Borg

## Abstract

Upregulation of the developmental Wnt/planar cell polarity pathway is observed in many cancers and is associated with cancer development at early and late stages. We recently showed that PRICKLE1 and VANGL2, two core Wnt/PCP components, are overexpressed in triple negative breast cancer and associated with poor prognosis. PRICKLE1 is a cytoplasmic protein phosphorylated by the poorly described serine/threonine kinase MINK1 which triggers its localization at the plasma membrane, a key step for its function. Knockdown experiments have demonstrated that MINK1 and PRICKLE1 contribute to TNBC cell motility and spreading *in vitro* and *in vivo*. However, the identity of MINK1 substrates and the role of MINK1 enzymatic activity in this process have not yet been addressed issues.

We carried out a phosphoproteomic strategy and identified novel MINK1 substrates including LL5β. LL5β is a membrane scaffold molecule that anchors microtubules at the cell cortex through its association with the plus-end MT proteins CLASPs to trigger focal adhesion disassembly. LL5β is a prominent member of the MINK1-PRICKLE1 protein complex and is directly phosphorylated by MINK1 that promotes its interaction with CLASP. Using a kinase inhibitor, we demonstrate that the enzymatic activity of MINK1 is involved in the protein complex assembly and localization, and cell migration. Analysis of gene expression data show that the concomitant up-regulation of *PRICKLE1* and *LL5*β mRNA levels encoding MINK1 substrates is associated with a poor metastasis-free survival for TNBC patients. Altogether, our results suggest that MINK1 may represent a potential target in TNBC.

## Introduction

Triple-negative breast cancer (TNBC) represents 15-20% of breast cancers and is characterized by poor prognosis. The main features of TNBC are molecular heterogeneity, high cell proliferation and metastatic dissemination, resistance to standard treatment, and a limited access to specific therapies (1). Transformation of mammary epithelial cells in cancer cells leads to the acquisition of specialized motile capacities that allow the most malignant cells to leave the primary tumor and invade the surrounding space and distant organs. For all cancers including TNBC such metastatic program accounts for 90% of deaths (2). Hence, a better understanding of the molecular mechanisms triggering the migration steps of TNBC cells is essential to define novel therapeutic targets and approaches. Recently, we and others discovered that an evolutionarily conserved group of genes controlling the establishment and maintenance of a morphogenetic process called Wnt/Planar Cell Polarity (Wnt/PCP) is deregulated in solid tumors, including TNBC, and that this alteration is tightly linked to cancer progression and dissemination (3,4).

The Wnt/PCP pathway utilizes a set of Wnt (co)-receptors and adaptors including PRICKLE1, a cytoplasmic scaffold protein that plays a pivotal role in embryonic development of *Drosophila* (5), Zebrafish(6) and Xenopus (7). This evolutionarily conserved adaptor contains a PET domain at the N-terminus followed by three LIM domains and a C-terminal farnesylation site (8). Previously, we showed that overexpressed *PRICKLE1* is a poor prognosis marker in TNBC (9,10). We and others have demonstrated that, at the molecular level, PRICKLE1 regulates the subcellular localization of associated proteins such as VANGL2 (8,11), RICTOR (10), and ARHGAP21/23 (12) to coordinate oriented cellular migration. PRICKLE1 has signaling activity through its association with RICTOR that regulates AKT phosphorylation in TNBC cell migration (10). During developmental processes, Prickle1 is asymmetrically localized in cells following phosphorylation by the serine-threonine kinase Mink1 (11). MINK1 is a conserved Ste-20-like serine-threonine kinase that belongs to the Germinal Center Kinase family involved in the migration of human fibrosarcoma and breast cancer cell lines (13). We previously demonstrated that MINK1 is tightly associated with PRICKLE1 and that the PRICKLE1-MINK1 complex relocates at the cell periphery of TNBC cells upon PRICKLE1 phosphorylation by MINK1 (11).

In the present study, we aimed to identify additional substrates of MINK1 in TNBC cells in order to better understand its signaling cascade and its implication in the prometastatic program. Using a combined strategy of protein purification and phosphoproteomics, we identified a set of potential MINK1 substrates including LL5β, an adaptor protein associated with PRICKLE1 (14). LL5β is a PH domain protein partially located at the cell membrane (15) that anchors microtubules (MTs) through its direct association with CLASPs, a family of MT plus-end proteins (16). This LL5β-CLASPs interaction leads to the disassembly of focal adhesions and promotes cell migration (17). We uncover a two-step phosphorylation cascade triggered by MINK1 that promotes the subcellular localization, association and function of PRICKLE1 and LL5β at the plasma membrane of TNBC cells. In addition, we show that combined overexpression of *PRICKLE1* and *LL5*β represents an independent poor prognosis marker in TNBC.

## Material and Methods

### Cell culture, reagents and antibodies

HEK293T, MDA-MB231, SUM149PT cells were obtained from the ATCC. Cells were grown in DMEM or DMEM/F12 containing 10% Fetal Calf Serum. Transfections were performed using Polyethyleimine (Santa Cruz), Lipofectamine 2000 and Lipofectamine LTX (Thermo). siRNA was used as reverse transfection using Lipofectamine RNAimax (Thermo). Cells were tested for Mycoplasma regularly. Antibodies targeting MINK1 were obtained from Bethyl. The following antibodies were obtained from Cell Signaling: α-AKT, α-pS473-AKT. CLASP2 antibody was obtained from Absea Biotechnology ltd. LL5β, Tubulin and actin antibodies were obtained from Merck. α-pT370-PRICKLE1 and α-PRICKLE were produced and purified from rabbits injected with the following peptides: FPGLSGNADDpTLSR and QETPEDPEEWADHEDY, respectively. KY05009 were purchased from Merck.

### Affinity purification, immunoprecipitation and western blot

48 hours post-transfection, cells were lysed with the TAP lysis buffer (Tris-HCl pH8.0 50mM, NaCl 150mM, EDTA 2mM, Glycerol 10%, NP40 0.1%) and incubated at 4°C for 30 minutes to solubilize proteins. Affinity purification and immunoprecipitations were performed using GFP-Trap beads (made in house) for 3 hours at 4°C. After extensive washes with lysis buffer, proteins were eluted with 2x Laemmli sample buffer and heated at 95°C for 5 min in the presence of β-mercaptoethanol (Sigma). Whole cell lysates or purified protein samples were resolved by SDS-polyacrylamide gel electrophoresis (SDS-PAGE) and transferred onto Biotrace NT Nitrocellulose Transfer Membranes (Cytiva). Western blotting was performed with antibodies as indicated in figures legend, followed by chemiluminescent detection using appropriate HRP-conjugated antibodies and ECL (Cytiva) reagent.

### *PRICKLE1* and *LL5β/PHLDB2* mRNA expression analysis in breast cancer samples

PRICKLE1 and LL5β/PHLDB2 mRNA expression in breast cancer was analyzed in our pooled data set including gene expression data of a total of 5,883 non-redundant pre-therapeutic samples of non-metastatic, non-inflammatory, primary, invasive breast cancers with and clinicopathological annotations. Additional details are available in Supplementary Methods.

### Migration assay

25,000 cells were seed on collagen precoated 6-wells plate. Cells were monitored using live cell imaging using Metamorph 7.8.1 and followed during 19 hours. Cells were then followed manually using ImageJ tracking plugin. The values for the assessment of migration speed, velocity and directionality were obtained using ImageJ software. Migration speed was determined by the ratio between the total distance and duration of cell migration. Cell velocity corresponds to the distance between the positions of the cells at the beginning and the end of the experiment divided by time. Cell directionality is the ratio between cell velocity and cell speed **(18)**. Wound healing assay was performed using insert from IBIDI. 70,000 cells were seeded and the next day the insert was removed. Cell migration was followed using live cell imaging and analysis was performed using in house generated script under Metamorph software.

### *In vitro* kinase assay

The MINK1 assay was purchased from Promega (Charbonnieres-les-bains, France). Enzyme, substrate, ATP, and inhibitors were diluted in Kinase Buffer as per the manufacturer’s instructions. Kinase reaction was performed in 384-wellplate in a final volume of 5 μL. Reaction was initiated using 1 μL of inhibitor for each concentration (1% DMSO), 1 μL of enzyme, and 3 μL of substrate/ATPmix (60 min, RT). Five microliter of ADP-GloTM reagent were used to stop kinase reaction by ATP depletion (40 min, RT). Then, ADP formed by kinase reaction was detected by adding 10 μL of Kinase Detection Reagent (30 min, RT). Luminescence was recorded using a PHERAstar plate reader. MINK1 kinase was used at optimized concentrations of 2 ng/well, ATP was used at 5 μM, DTT at 50 μM and the substrate of MINK1 was used at 0.25 ng/μL.

### Phosphoproteomic approach

MDA-MB231 cells were grown in media supplemented with heavy or light arginine and dyalysed FBS (Thermo) during 15 days until we reached near 100% isotope labeling. Cells were treated with shRNA targeting MINK1 (light sample) or Non-Targeting shRNA (heavy sample). shRNAs were cloned into PLKO backbone and lentiviral particles were produced using PSPAX and VSV-G systems in HEK293T cells. The following sequences of shRNA were used to target MINK1 (#03) AGCGGCTCAAGGTCATCTATG. Cells were lysed in RIPA buffer (20 mM HEPES pH 8.0, 9 M urea, 1 mM sodium orthovanadate, 2.5 mM sodium pyrophosphate, 1 mM β-glycerophosphate). 1-2×10^8^ cells per condition were used (20mg of protein). Cells were collected and sonicated at 15W output with 2 bursts of 30 sec each. The lysate was cleared by centrifugation at 20,000xg at 4°C for 15 min and supernatant were stored for mass spectrometry analysis. After protein quantification, lysates from each condition were mixed to 1:1 ratio and digested using Trypsin. Phospho-peptides were enriched using TiO2 column and analyzed by mass spectrometry.

### Mass spectrometry identification of phosphorylation site

Identification of LL5β phosphorylation site were performed using *in vitro* kinase assay followed by mass spectrometry analysis. Additional details are available in Supplementary Methods.

### Quantitative RT-PCR assays

MINK1, TNIK and MAP4K4 mRNA level were measured using RT-qPCR. Additional details are available in Supplementary Methods.

### Confocal imaging

Cells were seeded on coverslips pretreated with rat tail collagen (Roche). Cells were fixed using paraformaldehyde or ice-cold methanol followed by permeabilization using PBS/Triton X-100 at 0.2%. Cells were treated with the indicated antibody and imaged on confocal LSM 880 (Zeiss) with a UV laser and 63X objective. Confocal images were analyzed using ImageJ software.

### Image analysis

Lamellipodia structure were counted manually in each field and reported over the number of cells counted through the observation of DAPI staining of the nucelus. The colocalization of CLASP2 and LL5β were measured on area selected wthin the plasma membrane and enriched in microtubules ends as shown in Fig. 5B and Fig.5G. Colocalization were measured using ImageJ software and Pearson correlation coefficient were extracted.

### DNA constructs, siRNAs and chemical compounds

The following sequences of siRNAs were used to target MINK1 (#09) GGAACAAACUGCGGGUGUA, MINK1 (#10) GAAGUGGUCUAAGAAGUUC, MAP4K4 (#06) CAAAAGGGCUCAAAGACUA, MAP4K4 (#08) GAAAUACUCUCAUCACAGA, TNIK (#03) UAAGCGAGCUCAAAGGUUA, TNIK (#04) GAACAUACGGGCAAGUUUA.

The LL5α construct was obtained from Yuko Mimori-Kiyosue. LL5β cDNA was obtained from Joshua Sanes. PRICKLE1 constructs were described previously. Site-directed mutagenesis was performed using the Q5® site directed mutagenesis kit protocol (NEB). GST-LL5β was cloned into pGEX plasmid vector. KY05009 and MK2206 were obtained from Merck.

## Results

### A screening for MINK1 substrates identifies LL5 β as a putative candidate

Using the MDA-MB-231 TNBC cell as a model, we previously showed that downregulation of the MINK1-PRICKLE1 complex expression inhibits TNBC progression(10). MINK1 is supposed to directly phosphorylate PRICKLE1 to promote its promigratory function(11). To further explore the MINK1 mechanism of action and identify additional MINK1 substrates, we carried out a phosphoproteomic approach by using the Stable Isotope Labeling by Amino acids in Cell culture (SILAC) methodology followed by phosphopeptides enrichment and identification (**Fig. 1A**). We generated MDA-MB-231 cells stably transfected with shRNA control or shRNA MINK1 and cultured them in heavy or light media following the SILAC procedure. We confirmed the downregulation of MINK1 expression by western blot analysis (**Fig. 1B**). After lysis and affinity purification, a total of 1323 phosphopeptides differentially expressed were identified by mass spectrometry analysis. Among them, we selected phosphopeptides showing a log2 ratio difference above 1.5 accounting for a total of 44 peptides representing 37 proteins (**Fig. 1C, Supplementary Table 1**). Among the phosphopeptides downregulated in the shRNA MINK1 condition, we identified a MINK1 phosphopeptide, validating our procedure. Previously, we have shown that PRICKLE1 is not only a direct partner of MINK1 but also a substrate of the protein kinase(11). We cross-analyzed our phosphoproteomic dataset with the list of the MINK1-PRICKLE1 complex associated proteins identified in MDA-MB-231 cells that we published previously to uncover the putative direct substrates of the MINK1 kinase(9). This analysis selected five proteins being both binders and potential substrates of MINK1 (**Fig. 1D**). Among them, the most prominent protein associated with the MINK1-PRICKLE1 complex was PHLDB2/LL5β (hereafter named LL5β. We verified by western blot analysis that total LL5β expression was not altered by MINK1 downregulation (**Fig. 1B**). We then expressed GFP or GFP-PRICKLE1 in MDA-MB-231 cells and confirmed by coimmunoprecipitation followed by western blot analysis that endogenous LL5β was associated to PRICKLE1 (**Fig. 1E**). This interaction was mediated by the second LIM domain of PRICKLE1 and was increased with a PRICKLE1 mutant which contains a T370D mutation mimicking PRICKLE1 phosphorylation by MINK1 on threonine 370(11) (**Fig. S1**). To confirm the specificity of the association of PRICKLE1 with LL5β, we challenged the association with LL5α which shares 70% identity with LL5β. We overexpressed either GFP-LL5α or GFP-LL5β with FLAG-PRICKLE1 in HEK 293T cells. After FLAG immunopurification, we observed that LL5β, but not LL5α, was associated to PRICKLE1 (**Fig. 1F**). Together, our results show that, among the MINK1-PRICKLE1 complex-associated proteins, LL5β represents a putative phosphorylated substrate of the serine-threonine kinase MINK1.

**Figure 1:**
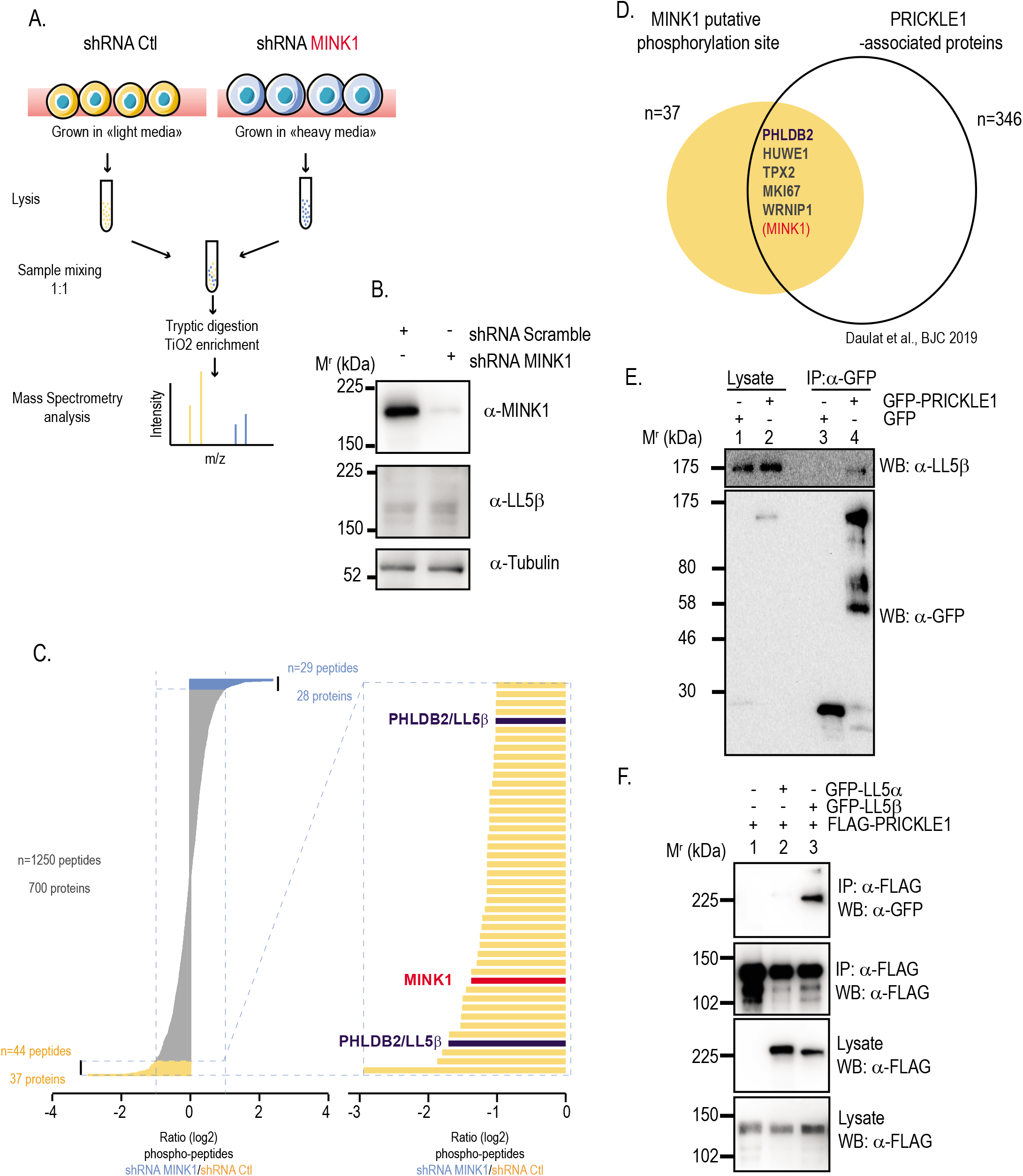
LL5β is associated to PRICKLE1 and is a putative substrate of the serine-threonine kinase MINK1. **A)** Schematic of the phosphoproteomic approach used in our study. **B)** Western blot analysis and detection of MINK1 expression in cell lysates. **C)** Histogram reporting the proteins and the peptides regulated upon MINK1 downregulation. **D)** Venn diagram representation of the cross analysis of the MINK1 putative phosphorylation sites and PRICKLE1-associated proteins. **E)** Co-immunoprecipitation assay followed by a western blot analysis to detect the interaction between GFP-PRICKLE1 and endogenous LL5β in MDA-MB-231 cells. **F)** Co-immunoprecipitation assay followed by a western blot analysis to determine the selectivity of association between LL5β and PRICKLE1. LL5β, but not LL5α, binds to PRICKLE1.

### LL5 β is a direct substrate of MINK1

To assess if LL5β is a direct substrate of MINK1, we performed an *in vitro* kinase assay mixing recombinant LL5β and MINK1 kinase followed by mass spectrometry analysis. To do so, we generated recombinant GST-LL5β and GST-PRICKLE1 in *E. coli* and 2μg of affinity-purified proteins were submitted to an *in vitro* kinase assay using the commercially available recombinant kinase domain of MINK1, in the presence or absence of ATP (**Fig. 2A, upper panel**). As we previously characterized threonine 370 (T370) of PRICKLE1 as a phosphorylation site for MINK1 (11), we used this post-translational modification as a positive control in our experiment. Hence, we could verify MINK1 phosphorylation of PRICKLE1 at T370 by western blot using a homemade antibody directed against this phosphosite (**Fig. 2A, lower panel**). After *in vitro* phosphorylation by MINK1, recombinant PRICKLE1 and LL5β were analyzed by mass spectrometry to identify the phosphorylation sites. Our analysis confirmed that PRICKLE1 is phosphorylated by MINK1 at threonine 370 as well as at other sites (**Fig. 2B**). Two phosphorylation sites, threonine 894 and threonine 217, were identified in LL5β (**Fig. 2B**), threonine 894 being one of the phosphosites detected in the SILAC screening (**Fig. 1B**). Alignment of the amino-acid sequence encompassing threonine 894 showed that this sequence was conserved among species (i.e. human, zebrafish, mouse) (**Fig. 1C**). In addition, this sequence fits very well with the typical consensus sequence motif of the MINK1 family kinase activity (19). Of note, this phosphorylation site is located within the domain of LL5β required for CLASP binding (**Fig. 1C**) (16). Together, our data show that MINK1 can directly phosphorylate LL5β at threonine 894.

**Figure 2:**
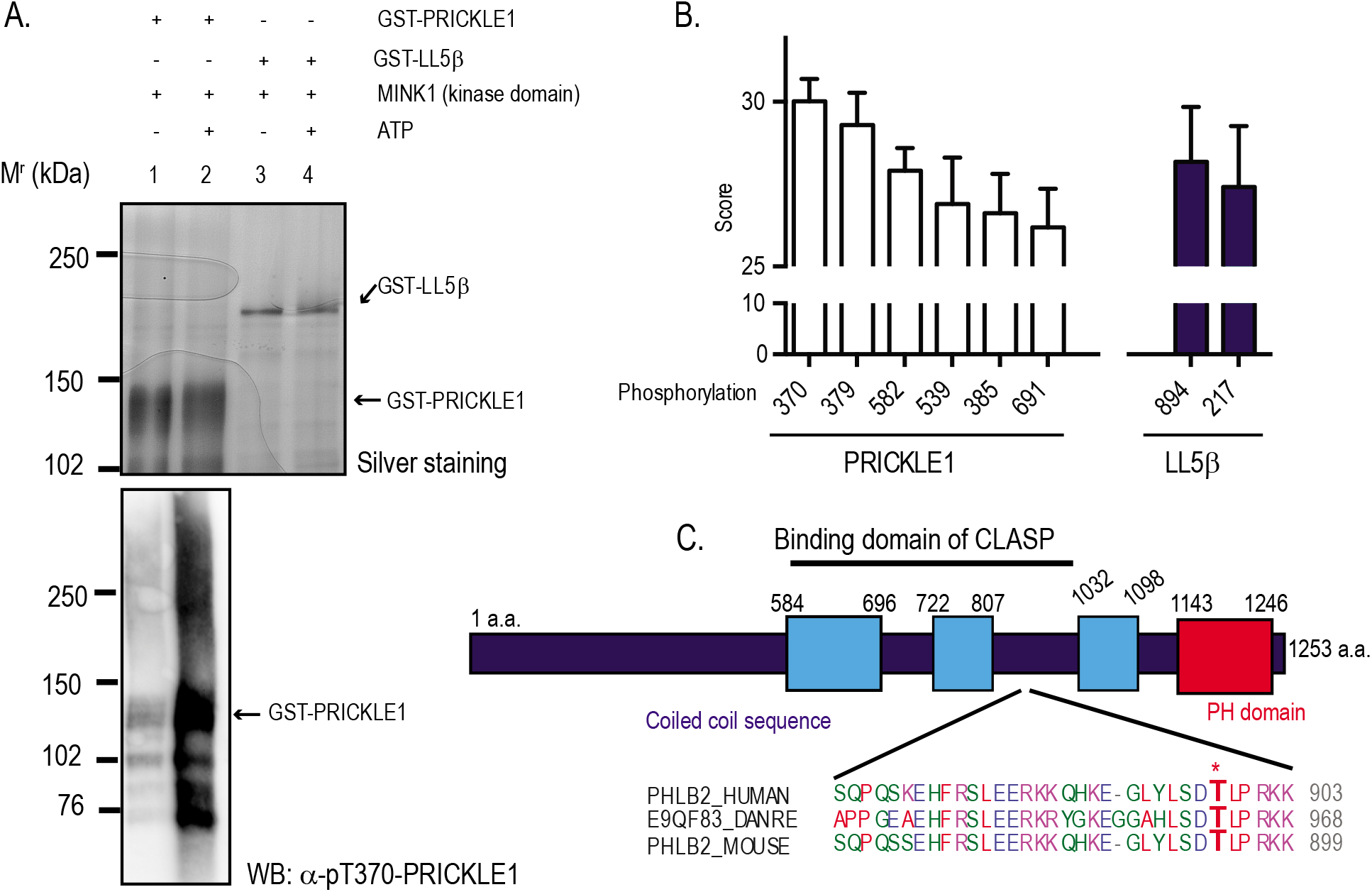
LL5β is a direct substrate of MINK1. **A)** Silver staining of acrylamide gel loaded with purified GST-PRICKLE1 and GST-LL5β subjected to an *in vitro* kinase assay (upper panel). Western blot analysis of the PRICKLE1 kinase assay using α-pT370-PRICKLE1 antibody (lower panel). **B)** Representation of phosphopeptides identified in GST-PRICKLE1 and GST-LL5β by mass spectrometry analysis. **C)** Schematic of LL5β and amino-acid sequence alignment of the peptide containing threonine 894 (asterisk) phosphorylated by MINK1.

### Inhibition of MINK1 catalytic activity phenocopies loss of MINK1

The role of MINK1 in the promigratory process of TNBC cells was previously demonstrated using siRNA or shRNA knockdown experiments (10). We decided to assess the role of the catalytic activity of MINK1 in cell migration using a commercially available protein kinase inhibitor. As MINK1 shares a high degree of homology with its paralogues TNIK and MAP4K4 regarding its ATP pocket (20), we tested KY05009, a chemical compound previously described as a TNIK inhibitor (21). We set up an *in vitro* kinase assay using the recombinant active kinase domain of MINK1 and found that KY05009 inhibited MINK1 with an IC50 of 1.2 nM (**Fig. 3A**). We previously showed that downregulation of MINK1 in TNBC cells leads to an increase of focal adhesion size and cell spreading, and to a decrease of cell motility (10). We next treated MDA-MB-231 cells with KY05009 and stained the cells with anti-vinculin antibody to assess the size of focal adhesions by immunofluorescence. We observed that KY05009 treatment increases the size of focal adhesions (**Fig. 3B**, **3C**) and cell spreading (**Fig. 3D, 3E**). MDA-MB-231 cells were also individually treated with MINK1, MAP4K4 and TNIK siRNAs, seeded on collagen coated coverslips and stained for actin (**Fig. S2**). SiRNAs efficiency were validated by RT-qPCR (**Fig. S2C**). Only MINK1 downregulation led to a similar cellular phenotype than KY05009 treatment with a strong cytoskeleton reorganization and higher cell spreading (**Fig. S2A, quantification in Fig. S2B**). We previously showed that downregulation of MINK1 led to a decrease of AKT phosphorylation and activity(10). Serum starved or FCS-treated MDA-MB-231 cells were treated with 1μM of KY05009 and analyzed for AKT phosphorylation. Stimulation with 5% FCS leads to phosphorylation of AKT on serine S473. This phosphorylation was strongly decreased by KY05009 treatment (**Fig. 3F**). Together, our data suggest that, in MDA-MB-231 cells, KY05009 behaves as a MINK1 inhibitor that phenocopies loss of MINK1 (10).

**Figure 3:**
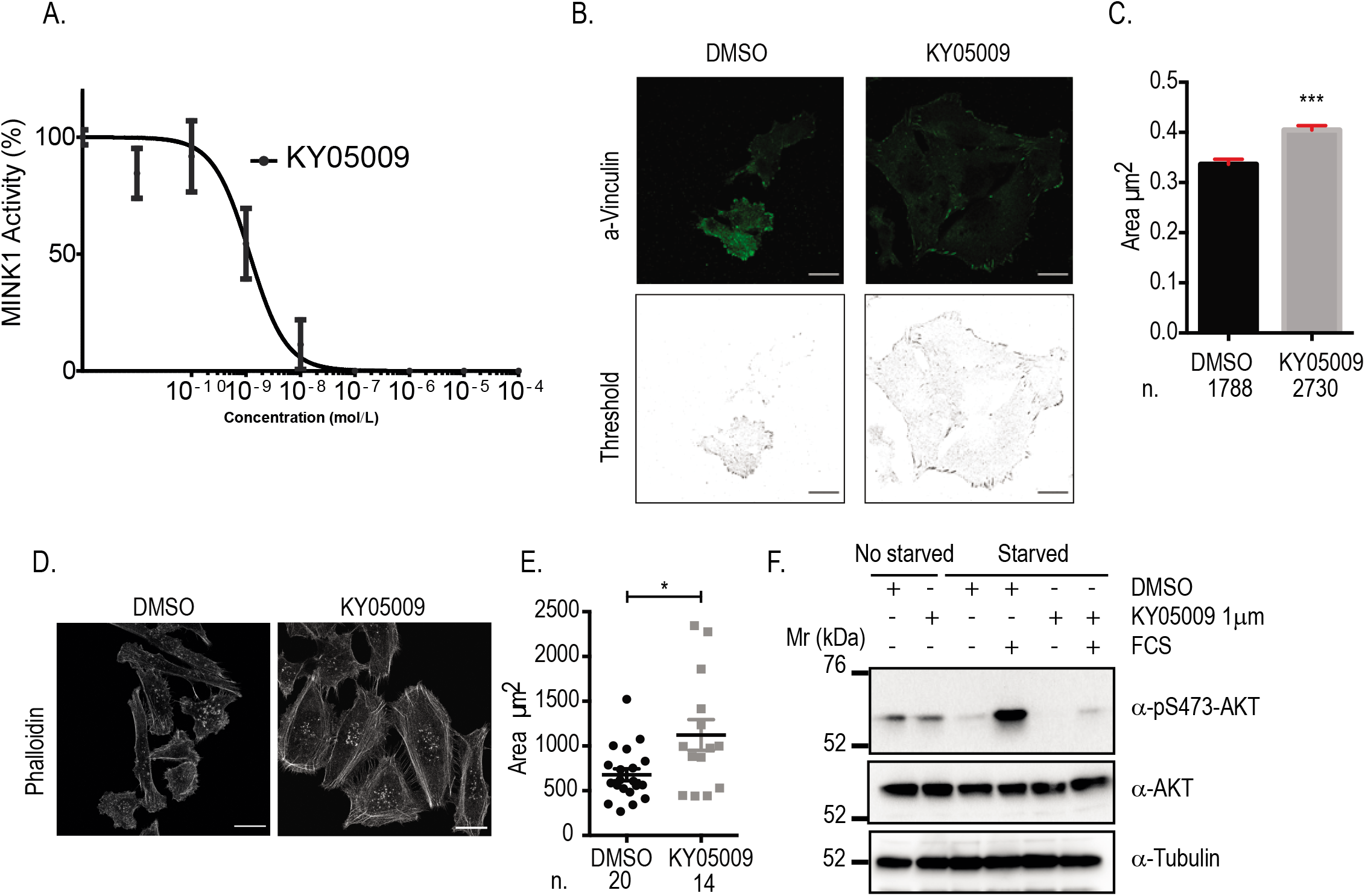
Characterization of the MINK1 inhibitor. **A)** *In vitro* kinase assay using a PRICKLE1 peptide encompassing the MINK1 phosphorylation site. Determination of the IC50 of KY05009. **B)** α-Vinculin staining of MDA-MB-231 cells seeded on collagen-coated coverslips and image treatment using ImageJ. **C)** Quantification of the area of the structures stained by Vinculin. **D)** Phalloidin staining and measurement of cell size (**E**). **F)** AKT activation in MDA-MB231 cells treated or not with KY05009. Scale bar 20μM. Image are representative of three independent experiments. Statistics were performed using the results of three independent experiments using Student’s t-test. *<0.05; ***P<0.001. Mean and SEM are represented.

### MINK1 activity promotes the association of PRICKLE1 with LL5 β and its subcellular localization

We then decided to explore the role of MINK1 catalytic activity in the formation and localization of the MINK1-PRICKLE1-LL5β complex. We expressed epitope-tagged versions of the three partners in HEK 293T cells and assessed their interaction. We found that GFP-LL5β associated to FLAG-MINK1 (**Fig. 4A**, lane 4) and that co-expression of HA-PRICKLE1, a direct MINK1 interactor, strongly potentiated the association between the two proteins (**Fig. 4A**, lane 3). We next used MDA-MB-231 cells which endogenously express LL5β and stably expressed GFP-PRICKLE1. We then performed an immunoprecipitation of GFP-PRICKLE1 using GFP antibodies and detected the presence of LL5β associated to PRICKLE1. This interaction was decreased by two independent MINK1 siRNAs (**Fig. 4B**). To determine if the catalytic activity of MINK1 is required for this association, we treated MDA-MB-231 cells with 1μM of KY05009. This treatment clearly decreased the association between PRICKLE1 and LL5β (**Fig. 4C**). As seen in **Fig. S1**, PRICKLE1 is bound to LL5β through its LIM2 domain and this interaction is positively regulated by the phosphorylation of PRICKLE1 by MINK1 on threonine 370. We next examined by immunofluorescence and confocal analysis the subcellular localization of endogenous PRICKLE1 in MDA-MB-231 cells using a homemade anti-PRICKLE1 antibody whose specificity was validated using two independent siRNAs targeting PRICKLE1 (**Fig. S3**). We observed that, in MDA-MB-231 cells, PRICKLE1 was localized at the plasma membrane in an actin-rich sub-region within the lamellipodia. Downregulation of MINK1 (**Fig. 4D**) or inhibition of its catalytic activity (**Fig. 4E**) led to a profound change of cellular morphology as previously described(10). In MINK1-deficient cells, we observed a drastic relocalization of PRICKLE1 from the plasma membrane to actin bundles. Similar observations were obtained usinng SUM149PT, another TNBC cell model (**Fig. S4**). Furthermore, we monitored LL5β subcellular localization upon MINK1 and PRICKLE1 downregulation by siRNAs or treatment with KY05009. We did not observe any influence of MINK1 or PRICKLE1 expression on LL5β membrane localization (**Fig. 4F**). Altogether, our data suggest that MINK1 activity promotes PRICKLE1, but not LL5β, localization at the plasma membrane.

**Figure 4:**
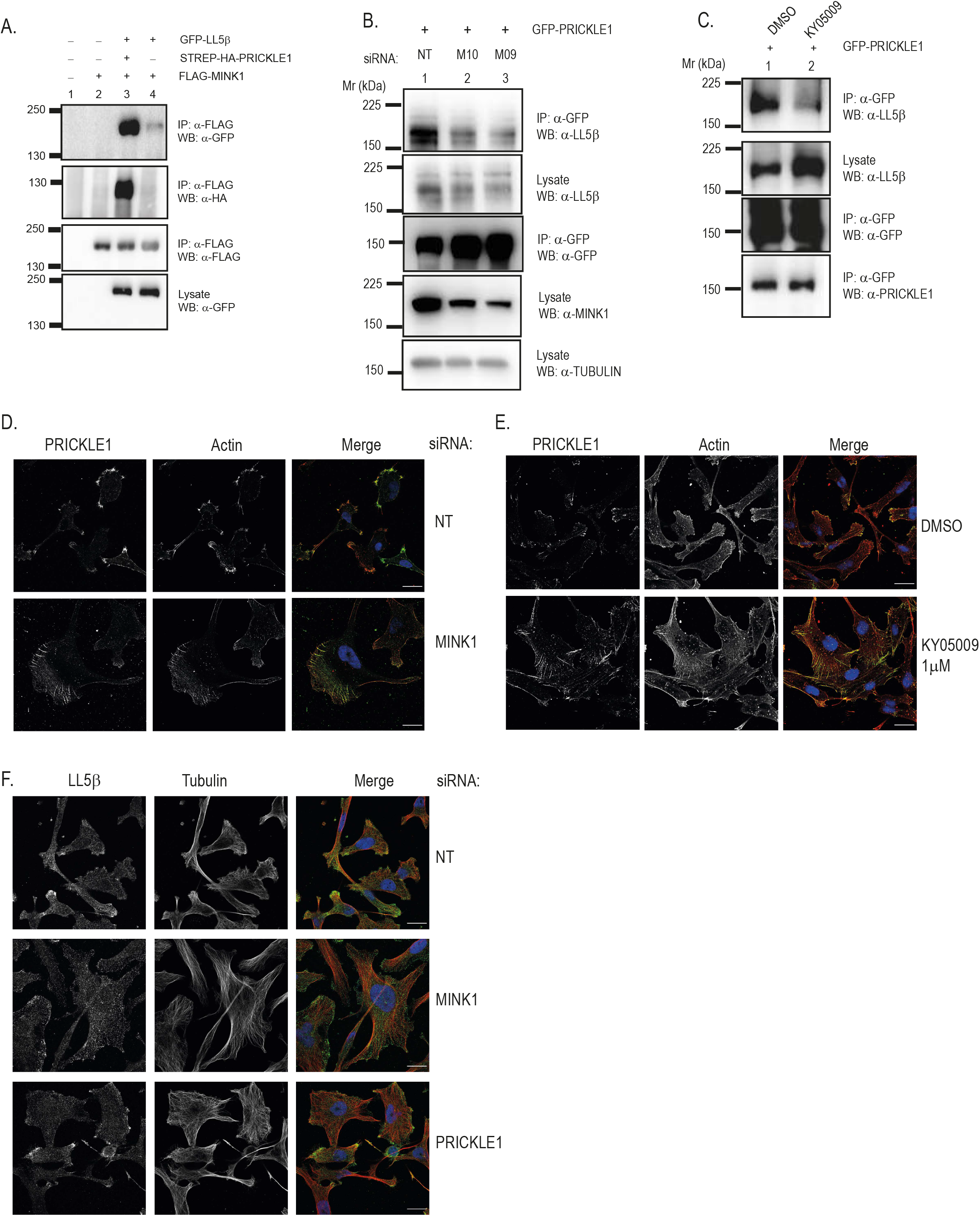
The MINK1 catalytic activity promotes PRICKLE1 association with LL5β and localization at the plasma membrane. **A)** Co-immunoprecipitation experiment showing the association between epitope-tagged MINK1 and LL5β in HEK 293 T cells. **B)** Co-immunoprecipitation in MDA-MB-231 cells of endogenous LL5β with GFP-PRICKLE1 in the presence of control (NT) or MINK1 siRNAs (M10, M09). **C)** Co-immunoprecipitation of endogenous LL5β with GFP-PRICKLE1 under KY05009 treatment. **D)** PRICKLE1 and actin staining on MDA-MB231 cells treated with a siRNA targeting MINK1. **E**) MDA-MB-231 cells treated with KY05009 and stained with PRICKLE1 and actin antibody**. F)** MDA-MB-231 cells treated with the indicated siRNAs and stained for tubulin and LL5β. Scale bar 20μM. Image are representative of three independent experiments.

### The LL5 β-CLASP2 interaction is promoted by MINK1-dependent phosphorylation

Previous results demonstrated that LL5β anchors microtubules (MTs) at the plasma membrane through its direct association with CLASPs (Cytoplasmic Linker Associated Proteins), a family of MT plus-end binding proteins (16,17). CLASP2 was present in the list of PRICKLE1-associated complex purified in MDA-MB-231 cells (9). Interestingly, the threonine 894 of LL5β phosphorylated by MINK1 lied within the binding domain of CLASPs (**Fig. 2C**). As expected, GFP-LL5β stably expressed in MDA-MB-231 cells could be readily coimmunoprecipitated with endogenous CLASP2 (**Fig. 5A**). However, we found that downregulation of MINK1 strongly diminished this interaction suggesting that the association between LL5β and CLASP2 is partially dependent on the MINK1-PRICKLE1 complex. We assessed the colocalization between LL5β and CLASP2 by immunofluorescence in MDA-MB-231 cells and confirmed that LL5β was colocalized with CLASP2 within the lamellipodia. Downregulation of MINK1 using siRNA treatment affect their colocalization. (**Fig. 5B**). We observed less lamellipodia at the periphery of cells when MINK1 expression is downregulated (**Fig. 5C**). Furthermore, we quantified the colocalization of LL5β and CLASP2 in regions enriched in tubulin at the plasma membrane and observed a decrease of the Pearson Correlation Coefficient (PCC) in the absence of MINK1 (**Fig. 5D**). Thus, MINK1 promotes CLASP2-LL5β interaction and colocalization at the plasma membrane.

**Figure 5:**
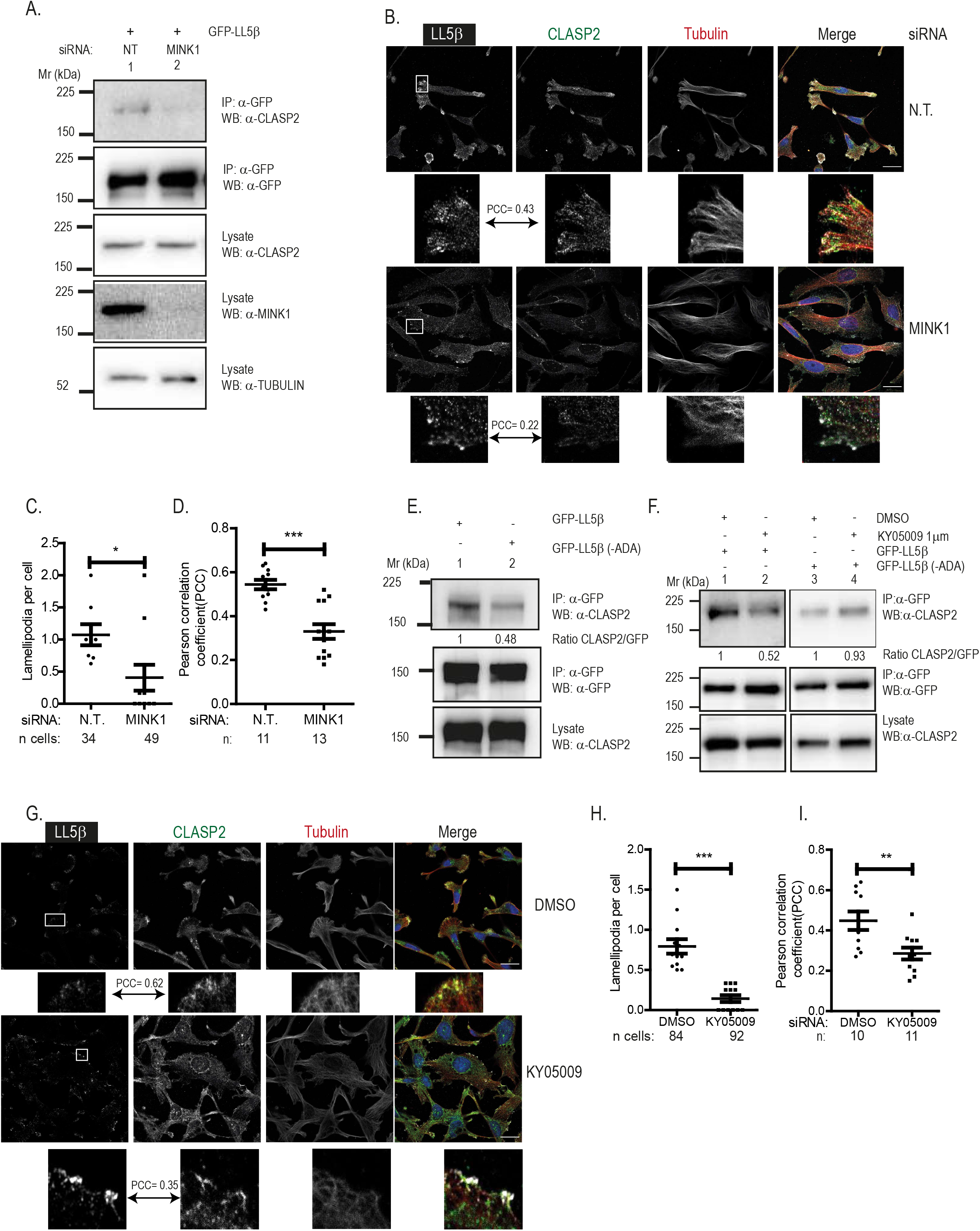
Phosphorylation of LL5β is required for its interaction with CLASP in MDA-MB-231 cells. **A)** Co-immunoprecipitation between LL5β and CLASP2 is decreased by loss of MINK1. **E)** Co-immunoprecipitation experiment between LL5β and CLASP2 is decreased by mutation of the threonine 894 phosphosite in LL5β (mutant ADA). and influence of 1μM of KY05009 treatment during 16 hours. **F)** Influence of KY05009 treatment on the interaction between CLASP2 and the wild type and mutant forms of LL5β. **B)** and **E)** Colocalization of LL5β and CLASP2 under treatment with the indicated siRNAs **(B)** or KY05009 **(E)**. **C)** and **H)** Quantification of lamellipodia structure per cell. **D)** and **I)** Pearson correlation coefficient (PCC) measured for colocalization between CLASP2 and LL5β. Example of colocalization area measured is given in **(B)** and **(G)**. Scale bar 20μM. Image are representative of three independent experiments. Statistics were performed using the results of three independent experiments using Student’s t-test. *<0.05; **P<0.01; ***P<0.001. Mean and SEM are represented.

We next examined if the association between LL5β and CLASP was regulated by MINK1 phosphorylation of threonine 894. We first generated a mutant of LL5β (-ADA) where serine 892 and threonine 894 were mutated in alanine. We mutated both residues to exclude a possible alternative phosphorylation of serine 892 by MINK1. We performed immunoprecipitation with anti-GFP antibody in lysates from MDA-MB-231 cells stably expressing GFP-LL5β or GFP-LL5β (-ADA). Western blot analysis showed that the association between LL5β and CLASP2 was decreased by 2-fold with GFP-LL5β (-ADA) compared to GFP-LL5β (**Fig. 5E**). To further evaluate if the remaining interaction could be due to other MINK1-dependent phosphorylation events, we treated the cells with KY05009 prior lysis and immunopurification. We observed that treatment of MDA-MB-231 cells expressing GFP-LL5β with KY05009 led to a decreased of GFP-LL5β-CLASP2 binding in the same range than between GFP-LL5β (-ADA) and CLASP2 (**Fig. 5F,** left panel). However, KY05009 treatment did not further decrease the binding between GFP-LL5β (-ADA) and CLASP2 suggesting that no other phosphorylation event due to MINK1 is implicated in the LL5β-CLASP2 association (**Fig. 5F**, right panel). As with MINK1 siRNA treatment (**Fig. 5B**), KY05009 treatment of MDA-MB-231 cells led to a loss of colocalization of LL5β and CLASP2, both proteins being still localized at the plasma membrane (**Fig. 5G**). As with MINK1 siRNA, treatment with KY05009 leads to a decrease of the number of lamellipodia per cell (**Fig. 5H**). Furthermore, we also observed a decrease in the co-localization of LL5β and CLASP2 at the plasma membrane (**Fig. 5I**). Altogether, our data showed that MINK1 phosphorylates LL5β within the CLASP2 binding site and that this event promotes the complete LL5β-CLASP2 association at the cell cortex.

### Inhibition of MINK1 catalytic activity inhibits cell motility

Our study highlights the role of the kinase activity of MINK1 in the assembly of the PRICKLE1-LL5β-CLASP1/2 complex. Considering its implication in cell migration, we used two complementary approaches to assess the influence of MINK1 catalytic activity in MDA-MB-231 cell motility. We monitored single cell motility and observed a decrease of the cumulative distance and Euclidean distance travelled by MDA-MB-231 cells when treated with KY05009 (**Fig. 6A**, **6B**). KY05009 treatment had also an impact on the speed of cells and no impact on their directionality (**Fig. 6B**). Furthermore, we observed that cell migration was also decreased under KY05009 treatment in a wound healing assay (**Fig. 6C**, **6D**). These results were well correlated with our previous results using MINK1 and PRICKLE1 siRNAs in MDA-MB-231 cells^12^. We obtained similar results using SUM149PT, another TNBC cell line, with MINK1 and PRICKLE1 siRNAs (**Fig. 6E**) and KY05009 (**Fig. 6F**). In addition, similarly to the experiments done in MDA-MB-231 cells^12^ (**Fig. 3 and 4, Fig. S2**), we obtained a similar increased of cell size and actin cytoskeleton reorganization in SUM149PT cells (**Fig. S4**). Together our data demonstrate that the catalytic activity of MINK1 is required for TNBC cell migration.

**Figure 6:**
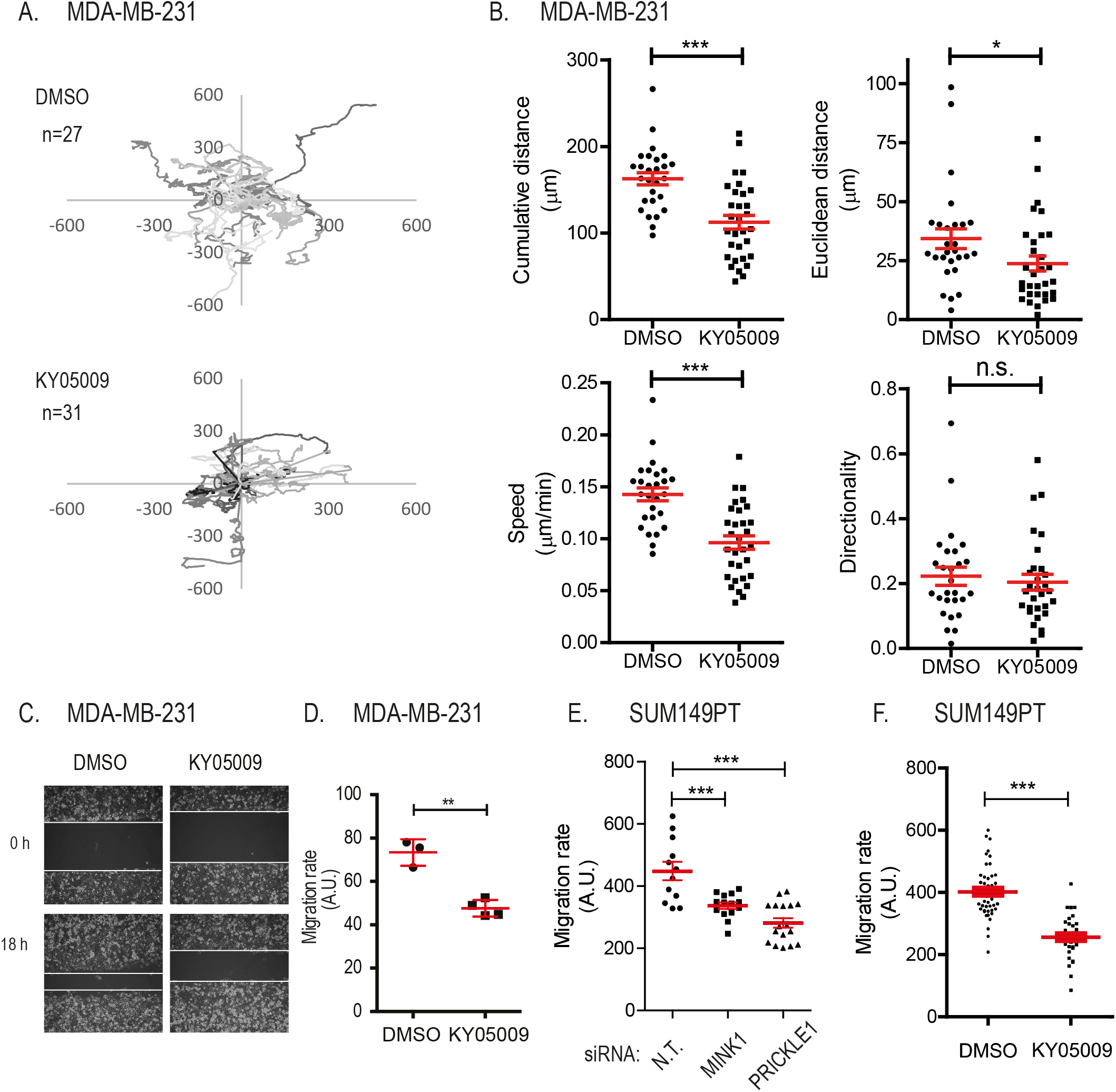
Inhibition of the MINK1 catalytic activity inhibit cell motility. **A-B)** Single MDA-MB-231 cell migration and treatment with KY05009 plot **(A)** and quantification **(B). C-D)** Wound healing assay using MDA-MB-231 cells treated or not with KY05009. Image taken at 0 and 16 hours **(C)** and quantification **(D). E-F)** Wound healing of SUM149PT cells treated with the indicated siRNAs **(E)** or KY05009 **(F).** Experiments were performed at least three times. Mean and SEM are represented. Statistical analysis was performed against control using Student’s t-test. *<0.05; **P<0.01; ***P<0.001.

### Unfavorable prognostic value of combined PRICKLE1 and LL5 β expression in TNBC

We have previously shown that *PRICKLE1* upregulation is associated with poor metastasis free survival (MFS) in basal breast cancer(9,10), a molecular subtype mainly composed of TNBC, as well as in TNBC^11^. In the present series of TNBC, including 471 informative patients, we confirmed that *PRICKLE1* upregulation was associated with shorter MFS, with 72% 5-year MFS (95Cl, 64-81) *versus* 56% (95Cl, 40-64) in the *PRICKLE1*-low group and the *PRICKLE1*-high group respectively (p=0.0027, log-rank test) (**Fig. 7A**). We next assessed the prognostic value of *LL5*β*/PHLDB2* expression in this series. We observed that an *LL5*β*/PHLDB2* upregulation was associated with shorter MFS, with 70% 5-year MFS (95Cl, 64-77) *versus* 47% (95Cl, 38-58) in the *LL5*β*/PHLDB2*-low group and *LL5*β*/PHLDB2*-high group, respectively (p=0.0001, log-rank test) (**Fig. 7B**). Since both genes did not show correlation of their expression level (**Fig. S5A**), we tested an eventual complementary prognostic value. Interestingly, the patients with upregulation of both genes (“*PRICKLE1*-high/*LL5*β*/PHLDB2*-high” group) displayed 35% 5-year MFS (95Cl, 25-49), whereas patients with downregulation of both genes displayed 74% 5-year MFS (95Cl, 65-84) and those with opposite deregulation of both genes (up/down, and down/up) displayed 67% 5-year MFS (p=1.16E06, log-rank test) (**Fig. 7C**). Since these last three patients’ groups showed similar MFS, they were merged and thereafter designated “no-*PRICKLE1*-high/*LL5*β*/PHLDB2*-high” group). The 5-year MFS of this latter was 69% (95Cl, 64-76) *versus* 35% (95Cl, 25-49) in the “*PRICKLE1*-high/*LL5*β*/PHLDB2*-high” group (p=7.15E-08, log-rank test) (**Fig. 7D**). Interestingly, such prognostic value of the combined expression of *PRICKLE1* and *LL5*β*/PHLDB2* was not observed in the HR+/HER2− and HER2+ subtypes (**Fig. S5B-C**). The prognostic complementarity of expression of both genes in TNBC was confirmed using the likelihood ratio (LR) test. The 2-gene combination provided more prognostic information than *PRICKLE1* alone (ΔLR-X2=14.45, p<2.2E-16), and *LL5*β*/PHLDB2* alone (ΔLR-X2=9.97, p<2.2E-16) (**Fig. S5D**). Finally, in multivariate analysis for RFS (Wald test; **Fig. 7E**), the 2-gene combination maintained its prognostic value (p=1.90E-02), whereas pT and pN tended to remain significant (p<0.10). Altogether, these data suggested a permissive function of *PRICKLE1* and *LL5*β in the metastatic process of TNBC.

**Figure 7:**
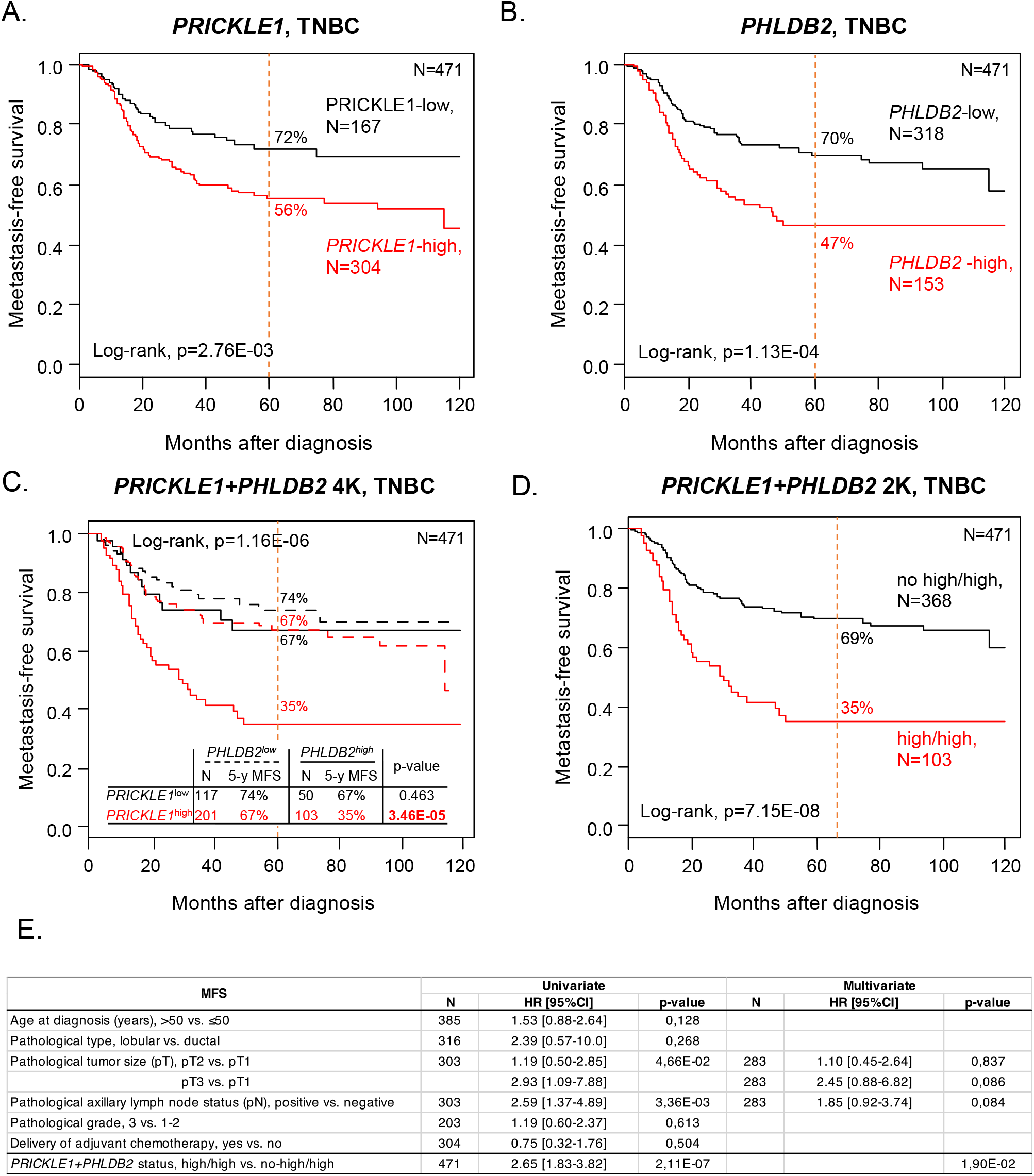
Prognostic value of PRICKLE1 and LL5β/PHLDB2 in TNBC. **A)** Kaplan-Meier MFS curves among TNBC patients according to *PRICKLE1* mRNA expression. **B)** Similar to A, but according to *LL5*β*/PHLDB2* mRNA expression. **C)** Similar to A, but according to both *PRICKLE1* and *LL5*β*/PHLDB2* mRNA expression: the patients are separated into four classes. **D)** Similar to **C**, but the patients are separated into two classes (“high/high” *vs.* “no-high/high”). **E)** Uni- and multivariate analysis for MFS in TNBC patients

## Discussion

Wnt/PCP signaling is now recognized as a pathway not only important in embryonic development and adult life but also in diseases, especially in cancer in which it is often deregulated (3). In the present study, we focused on TNBC, an aggressive subtype of breast cancer, and further studied MINK1, a Wnt/PCP prometastatic serine-threonine kinase with poorly described functions, especially regarding the role of its kinase activity and the identity of its substrates. We characterized a two-step phosphorylation event triggered by MINK1 in TNBC. In our working model (**Fig. 8**) based on our previous findings (10,11) and on the present study, MINK1 phosphorylates PRICKLE1, another Wnt/PCP molecule, at threonine 370 and, in doing so, promotes PRICKLE1 localization to the leading edge of migratory TNBC cells (10). At this location, MINK1, in complex with PRICKLE1, associates with LL5β that is then directly phosphorylated by MINK1 (**Fig. 1** and **Fig. 2**). This second phosphorylation event enhanced the association between CLASP2 and LL5β, which leads to an increase of cell motility (**Fig. 6**). It has been shown that CLASPs promote microtubule stabilization and tether microtubules to focal adhesions (FAs). In doing so, CLASPs facilitate the local delivery of exocytose proteins and extracellular matrix degradation releasing the integrin-matrix connection which in turn promotes integrin internalization and FA disassembly (17). We have shown that MINK1 and PRICKLE1 are involved in FA dynamics and promote the internalization of β1-integrin similarly to LL5β (10). We propose that the MINK1-PRICKLE1-LL5β-CLASP2 protein complex is involved in the disassembly of FAs and cell migration (**Fig. 8**).

**Figure 8:**
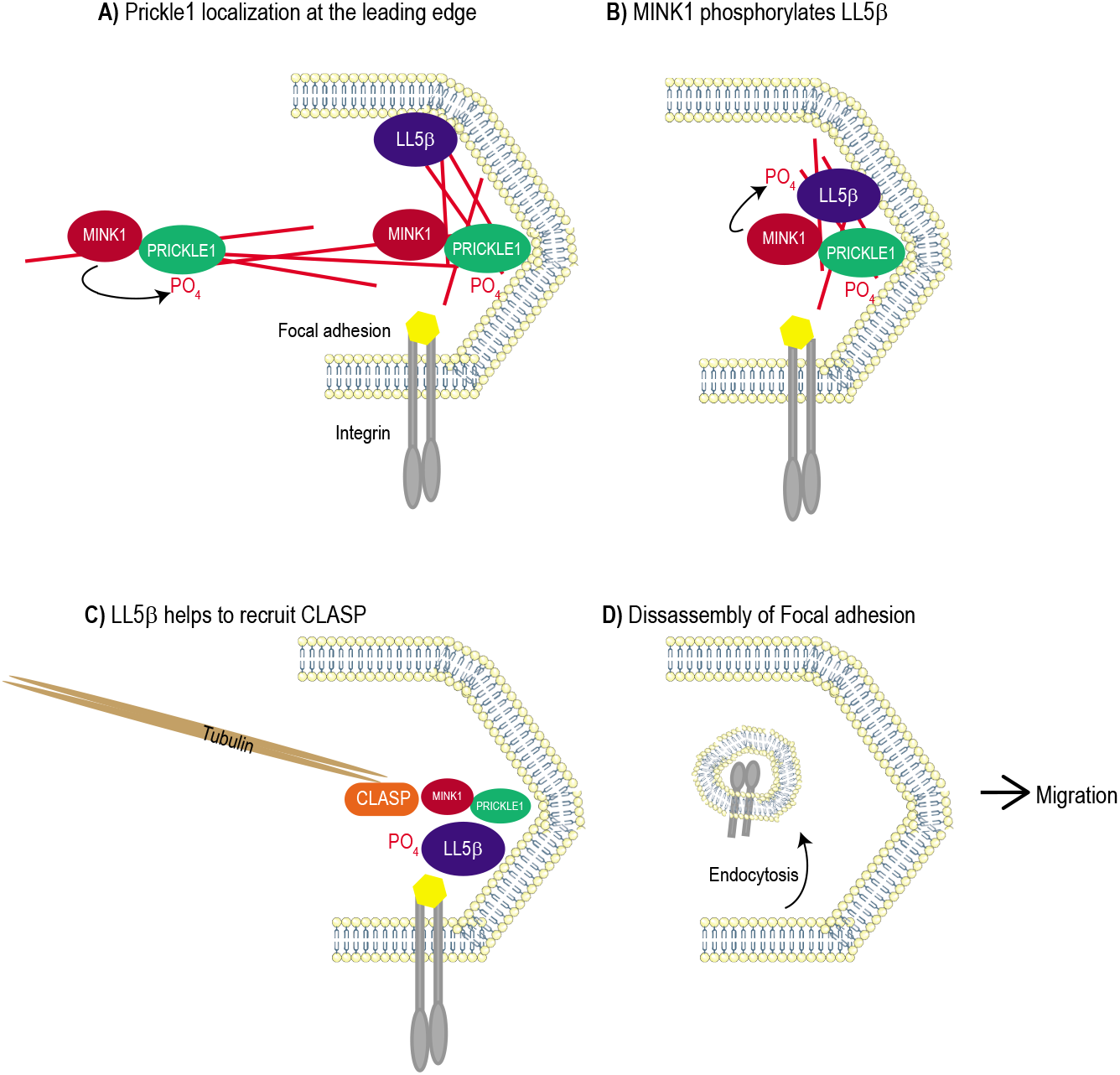
Working model. **A)** MINK1 phosphorylates PRICKLE1 and promotes it subcellular localization within the lamellipodia of the cell and PRICKLE1 binds to LL5β. **B)** MINK1 phosphorylates LL5β. **C)** This second phosphorylation event allows the direct binding of CLASPs to LL5β and tether the microtubule to the focal adhesion. **D)** Altogether this double step of phosphorylation promotes the dissambly of the focal adhesion and promotes cellular migration.

MINK1 has two paralogues (TNIK and MAP4K4) which share high degree of homology within their kinase domain, especially the ATP pocket, rendering difficult the generation of specific inhibitors for each family member (20). In this report, we show that treatment of TNBC cells with KY05009, an inhibitor of TNIK, phenocopies the loss of MINK1 or PRICKLE1 in terms of actin cytoskeleton reorganization, cell spreading and AKT phosphorylation (**Fig. 3**). We used KY05009 to demonstrate the role of MINK1 activity in the regulation of the assembly of the PRICKL1-CLASP2-LL5β complex and the implication of MINK1 catalytic activity in the promigratory process (**Fig. 4–6**), a finding that should stimulate the future development of specific MINK1 inhibitors. Of note, a recent study showed that *mink1* knockout mice are viable (22) suggesting that MINK1 inhibition may have some therapeutic avenue in cancer treatment. Using a SILAC method and *in vitro* kinase assays, we defined threonine 894 as a MINK1 phosphorylation site in LL5β (**Fig. 1** and **Fig. 2**). A whole proteome database search for MINK1 phosphorylation consensus motives based on the phosphorylation sites of PRICKLE1 at threonine 370 and LL5β at threonine 894 (D[ST]Lx[RK][RK]) identified ARHGAP21 and ARHGAP23 as putative MINK1 substrates. Interestingly, these two Rho-GTPase proteins have been shown to be associated to PRICKLE1 and to promote cellular migration through the regulation of Rho activity (12). This result opens up another avenue for further investigations about these potential MINK1 substrates. SMAD2 was previously shown to be phosphorylated and subsequently inhibited by MINK1 in T helper type (23). We did not find SMAD2 phosphopeptides in the list of MINK1 substrates in MDA-MB-231 cells (**Fig. 1**) suggesting cell context specificity of MINK1 substrates or dependency to the PRICKLE1 interaction that may not occur in lymphocytes. In conclusion, in MDA-MB-231 cells, PRICKLE1 acts as a scaffold protein able to bring substrates like LL5β close to MINK1 and hence provide specificity, a common theme in cell signaling(24).

Previously, it has been shown that LL5β and CLASPs form a complex with ELKS at the cell cortex(16). Our analysis of the PRICKLE1-associated proteins identified by mass spectrometry revealed the presence of LL5β and CLASP2, but not ELKS, peptides(9). We hypothesize that our purification procedure was not appropriate to recover this interaction or that two independent CLASPs-LL5β protein complexes coexist, one containing ELKS and one containing MINK1-PRICKLE1. This issue will have to be addressed in future studies.

Using a novel anti-PRICKLE1 antibody, we observed that PRICKLE1 is localized in cortical actin-containing structures of TNBC cells (**Fig. 4**). Loss or inhibition of MINK1 led to a strong reorganization of the actin cytoskeleton and delocalization of PRICKLE1 in actin bundles without modifying LL5β and CLASP2 membrane localization (**Fig. 4D–F**). This observation is in line with the fact that the plasma membrane recruitment of LL5β, a phosphatidylinositol-3,4,5-triphosphate (PIP3) binding protein, is under the control of PI3 kinase activity(16). However, CLASP2 and LL5β are less colocalized in the remaining lamellipodia when MINK1 are downregulated or inhibited by KY05009 (**Fig. 5B–D, 5G–I**). Added to the fact that MINK1 inhibition impairs the association between CLASP2 and LL5β (**Fig. 5A, 5E-F**), this suggests that the activity of MINK1 plays a role in the regulation and localization of CLASP2-LL5β, an important membrane complex implicated in MT tethering and cell migration (**Fig. 8**).

We found that MINK1 controls, through direct phosphorylation, the activity of PRICKLE1 and LL5β, two pro-migratory proteins that, when co-upregulated in TNBC, represent very poor prognosis markers (**Fig. 7, Fig S5**). Interestingly, expression of each gene provided additional prognostic information to that of each protein alone, suggesting a likely synergic effect on the metastatic program. Moreover, the prognostic value is molecular subtype-dependent as it is only observed in TNBC.

In conclusion, we describe here a MINK1-dependent pathway involving members of the Wnt/PCP pathway. The dependency of MINK1 activity to Wnt/PCP-related receptors and extracellular signals remains uncertain for the moment; this issue should be addressed in the future. However, previous works found that TNIK phosphorylates the transcription factor TCF, a key component of the Wnt pathway, and represents a pharmaceutical target in colon cancers having a constitutively active TNIK/TCF-dependent Wnt signaling(25). MINK1, TNIK and MAP4K4 were also implicated in the Hippo signaling which is frequently deregulated in cancer (26). This suggests that the targeting of MINK1 but also of other members of its family may have potential therapeutic value in TNBC and other cancers.

## Fundings

This work was funded by La Ligue Nationale Contre le Cancer (Label Ligue JPB and FB, and fellowship to AMD), Fondation de France (fellowship to AMD), Fondation ARC pour la Recherche sur le Cancer (grant to JPB), and AVIESAN through the NANOTUMOR project. M.S.W. was a recipient of the Science without Borders PhD program from Brazil Coordenação de Aperfeiçoamento de Pessoal de Nível Superior (CAPES). The Marseille Proteomics (IBiSA) is supported by Institut Paoli-Calmettes (IPC) and Canceropôle PACA. Samples of human origin and associated data were obtained from the IPC/CRCM Tumor Bank that operates under authorization # AC-2013-1905 granted by the French Ministry of Research. Prior to scientific use of samples and data, patients were appropriately informed and asked to express their consent in writing, in compliance with French and European regulations. The project was approved by the IPC Institutional Review Board. JPB is a scholar of Institut Universitaire de France

## Acknowledgements

The authors wish to thank Dr Mukhtar Ahmad for the critical review of the manuscript, Emilie Beaudelet and Yves Toiron for technical assistance in mass spectrometry analysis, Sébastien Germain for assistance with the isolated cell trajectory analysis and Dr Rudra Kashyap for his help to generate recombinant PRICKLE1 and LL5β proteins.

## Author’s contributions

AMD designed the project, supervised experiments, analyzed data and wrote the manuscript. MSW prepared the samples for MS analysis. SA and LC analyzed the protein samples by MS. FB and PF analyzed the transcriptomic data, MK performed the small siRNA screen, JAB performed *in vitro* kinase assays. JPB supervised, analyzed, funded the project and wrote the manuscript.

## Notes

**Conflict of interest:** The authors declare no potential conflicts of interest

### Competing Interest Statement

The authors have declared no competing interest.

