## Supplementary figures and images for "The serine/threonine kinase MINK1 directly regulates the function of promigratory proteins"

### Figure S1

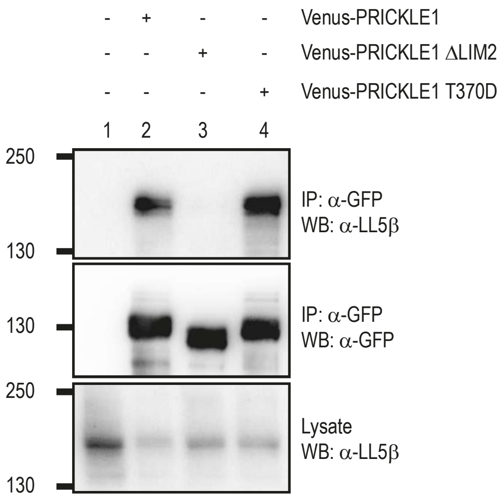

### Figure S2

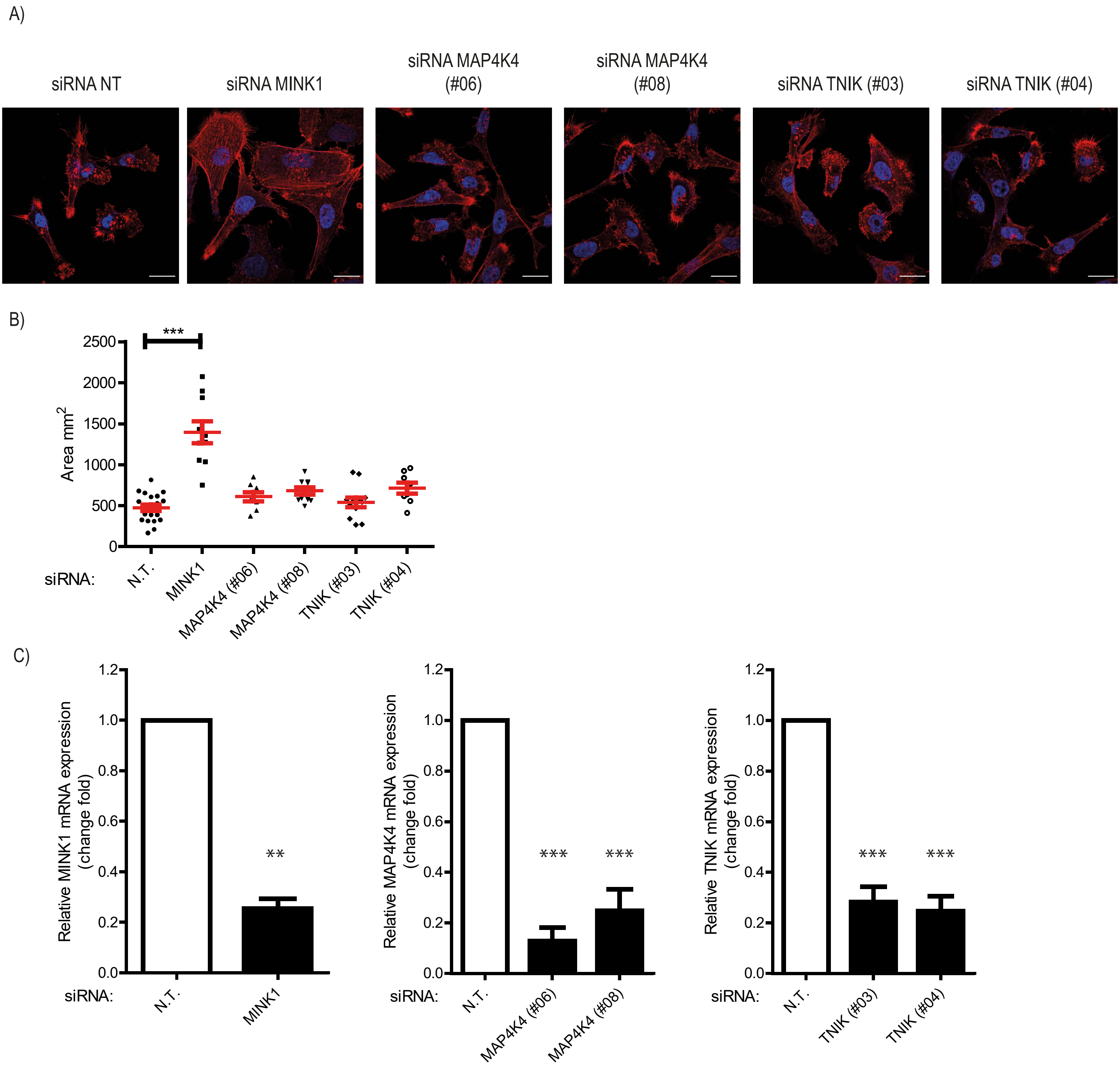

### Figure S3

FIGURE S3

siRNAs: NT PRICKLE1 (#02) PRICKLE1 (#04)

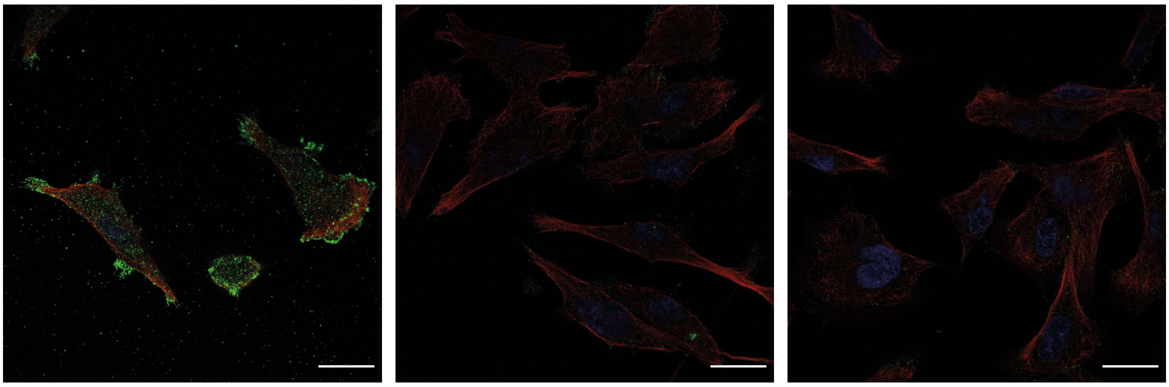

### Figure S4

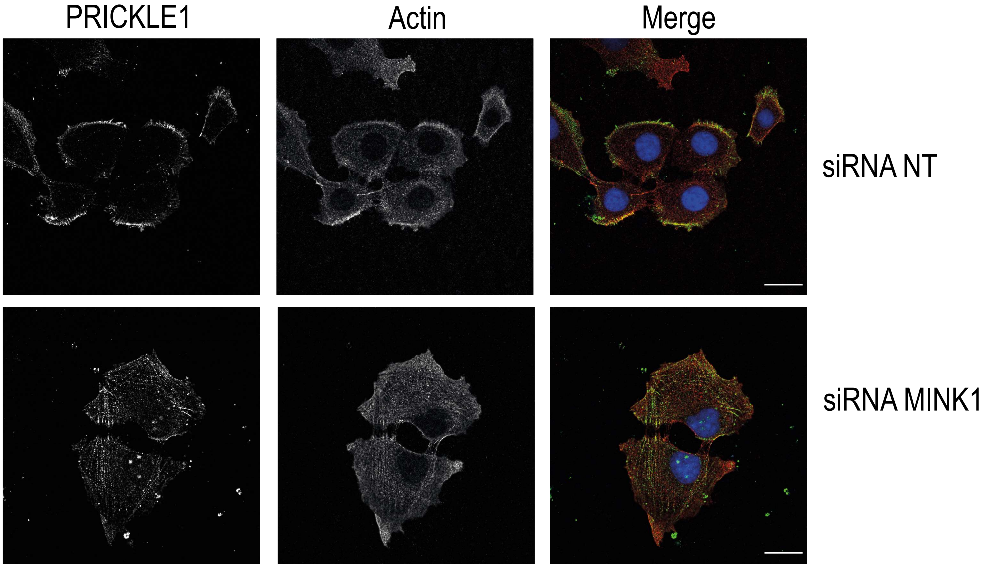

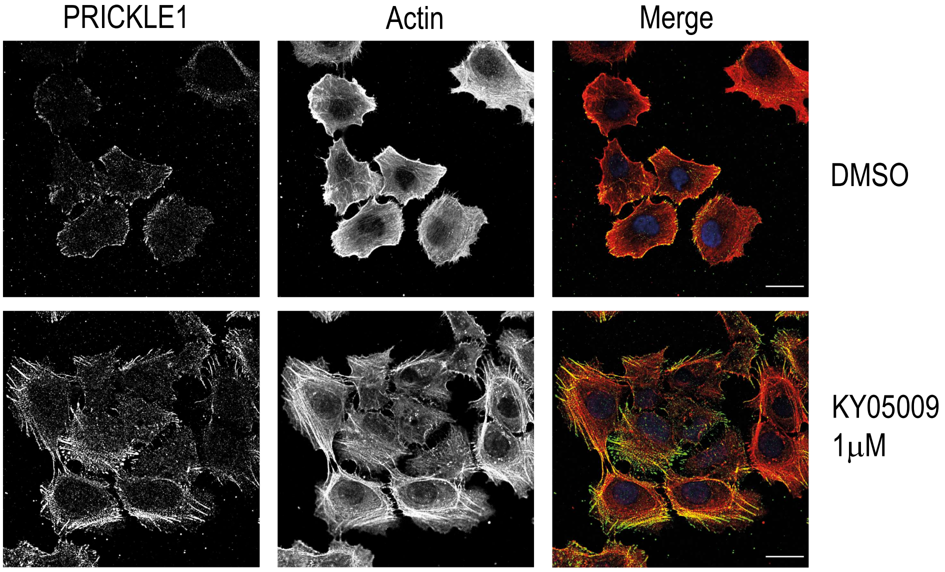

### Figure S5

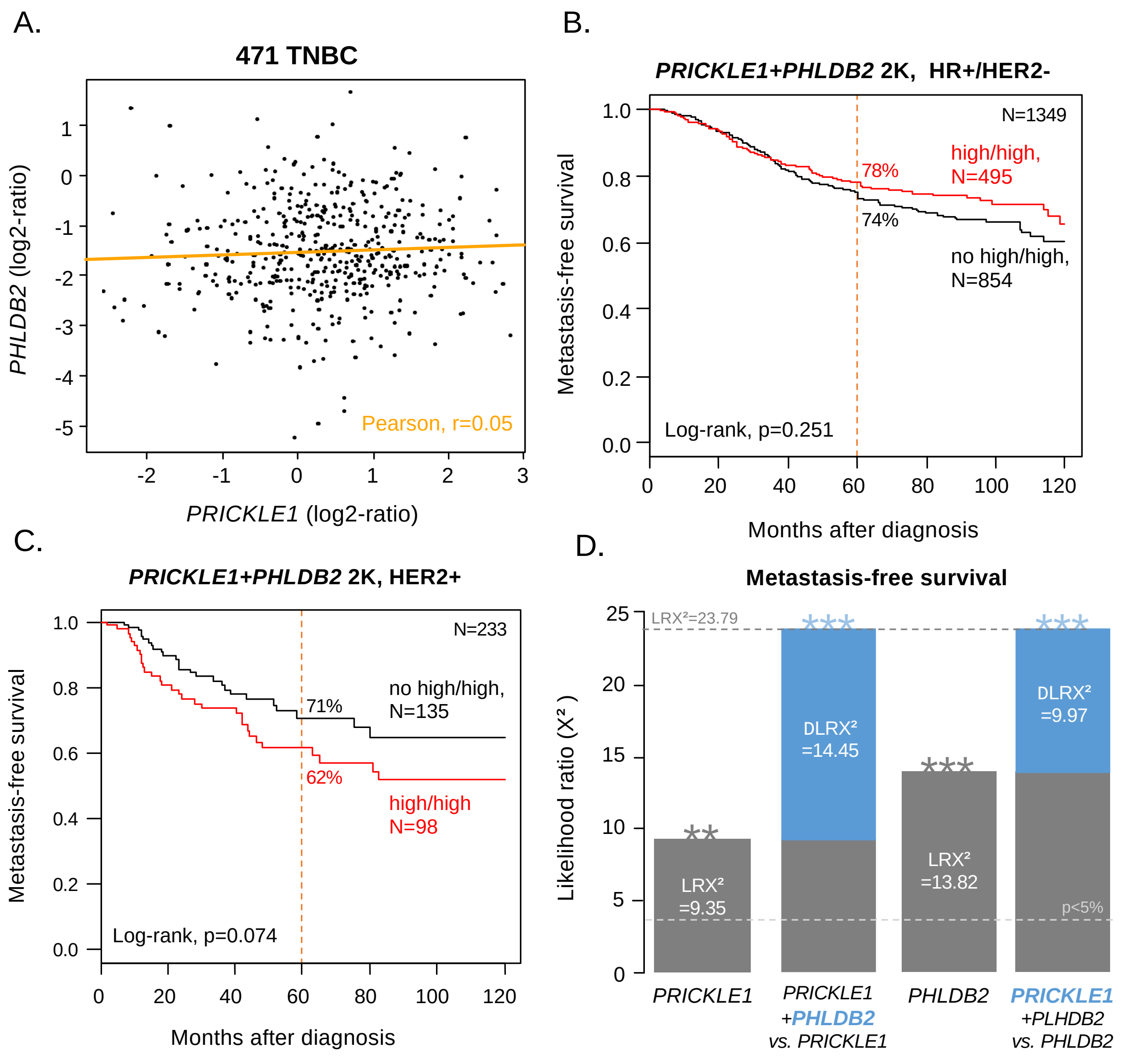
