## Supplementary Material and Methods for "The serine/threonine kinase MINK1 directly regulates the function of promigratory proteins"

***PRICKLE1*** and ***LL5β/PHLDB2*** **mRNA expression analysis in breast cancer samples**

*PRICKLE1* and *LL5β/PHLDB2* mRNA expression in breast cancer was analyzed in our own gene expression dataset (353 patients with invasive adenocarcinoma) coupled with publicly available data sets. The 17 public data sets comprising at least one probe set representing *PRICKLE1* and *LL5β/PHLDB2* were collected from the National Center for Biotechnology Information (NCBI)/Genbank GEO database, the European Bioinformatics Institute (EBI) ArrayExpress database or TCGA portal (Supplementary Table 2). This resulted in a total of 5,883 non-redundant pre-therapeutic samples of non-metastatic, non-inflammatory, primary, invasive breast cancers with and clinicopathological annotations (Supplementary Table 3). Data analysis required pre-analytic processing as previously described (2). To be comparable across data sets and to exclude bias from population heterogeneity, PRICKLE1 and *LL5β/PHLDB2* expression levels were standardized within each data set using the luminal A population as reference. All steps were done in R using Bioconductor and associated packages.

*PRICKLE1* and *LL5β/PHLDB2* upregulation and downregulation were defined using the median level as cut-off. The ER, PR, and ERBB2/HER2 statutes of samples were based on mRNA expression of *ESR1*, *PGR*, and *ERBB2* genes and defined as discrete values (positive/negative) using a 2-component Gaussian mixture distribution model (3). The molecular subtypes of tumors were defined as HR+/HER2- (ER- and/or PR-positive and HER2-negative), HER2+ (HER2-positive, regardless ER and PR), and TN (ER-, PR-, and HER2-negative). Metastasis-free survival (MFS) was calculated from the date of diagnosis until the date of distant relapse. Follow-up was measured from the date of diagnosis to the date of last news for event-free patients. Survivals were calculated using the Kaplan-Meier method and curves were compared with the log-rank test. Univariate and multivariate prognostic analyses for MFS were done using Cox regression analysis (Wald test). The variables tested in univariate analysis included patients’ age (> vs. ≤50 years), pathological tumor type (lobular *vs.* ductal), size (pT2, pT3 *vs.* pT1), grade (3 *vs.* 1-2), and axillary lymph node status (positive *vs.* negative), delivery of adjuvant chemotherapy (yes *vs.* no), and the *PRICKLE1/LL5β/PHLDB2* expression status (high/high *vs.* no-high/high). Multivariate analysis included the variables with a p-value inferior to 5% in univariate analysis. The likelihood ratio (LR) tests were used to assess the prognostic information provided beyond that of each gene, assuming a X2 distribution. Changes in the LR values (LR-ΔX2) quantified the relative amount of information of one Cox model compared with another. All statistical tests were two-sided at the 5% level of significance. Statistical analysis was done using the survival package (version 2.30) in the R software (version 2.15.2; http://www.cran.r-project.org/).

**Mass spectrometry identification of phosphorylation site**

Recombinant MINK kinase domain (kinase Logistics) was incubated with recombinant GST–PRICKLE1 proteins purified from Escherichia coli and immobilized on Glutathione beads. Phosphorylation reactions were performed in kinase buffer (25 mM HEPES, pH 7.4, 25 mM β-glycerophosphate, 25 mM MgCL2, 0.1 mM Na3VO4, 0.5 mM DTT) supplemented with 20µM ATP at 37°C for 1 hour. Reactions were stopped by five successive washes with kinase buffer and the addition of 4x Laemmli sample buffer. Proteins were resolved by SDS–PAGE, extracted and digested with Trypsin. The resulting peptides mixture analyzed by LC-MS/MS. Peptides and proteins identification were performed as described above but search parameters were modified to allow for the presence of one or more phosphate groups on serine, threonine or tyrosine residues. MS/MS spectra attributed to a phosphorylated peptide in PRICKLE1 and LL5β were manually inspected and validated to locate the position of the phosphoamino acid.

**Quantitative RT-PCR assays**

Total RNA was isolated from MDA-MB231 cells using the RNA isolation kit (RNeasy Mini Kit, Qiagen) according to manufacturer’s instructions. After DNase (Turbo DNA-free^TM^ Kit, Promega) treatment, a standard PCR (GoTaq G2 Green Master Mix, Promega) was perform on RNA samples to check genomic DNA contamination. cDNA was synthetized from 1 µg of total RNA using Reverse Transcriptase kit (SuperScript^TM^ II Reverse Transcriptase, Invitrogen). Quantitative real-time RT-PCR was performed using Syber green kit (Applied) and CFX96^TM^ Real-Time PCR System (Bio-Rad) as follows: 95°C for 3min, 40 cycles at 95°C for 15s, 60°C for 30s and 72°C for 30s. Total RNA was isolated from MDA-MB231 cells using the RNA isolation kit (RNeasy Mini Kit, Qiagen) according to manufacturer’s instructions. After DNase (Turbo DNA-freeTM Kit, Promega) treatment, a standard PCR (GoTaq G2 Green Master Mix, Promega) was perform on RNA samples to check genomic DNA contamination. cDNA was synthetized from 1 µg of total RNA using Reverse Transcriptase kit (SuperScriptTM II Reverse Transcriptase, Invitrogen). Quantitative real-time RT-PCR was performed using Syber green kit (Applied) and CFX96TM Real-Time PCR System (Bio-Rad) as follows: 95°C for 3min, 40 cycles at 95°C for 15s, 60°C for 30s and 72°C for 30s. M

INK1 forward primer: CTGATGTTGCTGGACCGAAG ,

MINK1 reverse primer: AGAATCTTGTTCCGGAGCCA ,

TNIK forward primer: AAGGTAACACGTTGAAAGAGGAG ,

TNIK reverse primer: AGTCAGCAAGACATTTTGCCC ,

MAP4K4 forward primer: GACTCCCCTGCAAAAAGTCTG ,

MAP4K4 reverse primer: GTCCATAGGTGCCATTTCCAA ,

ACTINB forward primer: CACCATTGGCAATGAGCGGTTC

ACTINB reverse primer: AGGTCTTTGCGGATGTCCACGT

**Supplementary files**

**Supplementary Table 1:** List of phosphopeptides identified by SILAC approach

**Supplementary Table 2:** List of breast cancer data sets included in the study

**Supplementary Table 3:** Clinicopathological characteristics of samples in the whole data set

**Figure S1:** Co-immunoprecipitation experiment in HEK293T cells showing that a PRICKLE1 mutant deleted of its LIM2 domain unable to bind MINK1 is not associated to LL5β (lane 3). A PRICKLE1 mutant with a phosphomimicked site (T370D) presents higher binding to LL5β (lane 4).

**Figure S2: Cellular phenotypes obtained following treatment with siRNAs targeting members of the MINK1 family. A)** MDA-MB-231 cells treated with the mentioned siRNAs were seeded on collagen-coated coverslips and fixed using PFA. Staining were performed with phalloidin. **B)** Quantification of cell area measured by ImageJ. **C)** Quantification by RT-qPCR of the expression of the genes targeted by siRNAs. Data represent at least three independent experiments and are reported as mean ±SD. Human β-actin was used as an internal standard. The relative quantification (fold change) of mRNA expression in the sample was estimated using the 2^-ΔΔC^_T_ formula. The statistical significance of the difference between control (transfected with no targeting siRNA) and specific siRNA- transfected samples was determined using Student’s t-test. **P<0.01; ***P<0.001.

**Figure S3: Validation of a homemade anti-PRICKLE1 antibody.** MDA-MB-231 cells treated with the indicated siRNAs were stained with anti-actin (red) and anti-PRICKLE1 antibody (green).

**Figure S4: Immunofluorescence of SUM149PT cells. A** and **B.** SUM149PT cells were treated as indicated and stained using anti-PRICKLE1 and -actin antibodies.

**Figure S5: PRICKLE1 and LL5β/PHLDB2 expression in TNBC. A)** Absence of **c**orrelation between *PRICKLE1* and *LL5β/PHLDB2* mRNA expression in TNBC patients. **B)** Kaplan-Meier MFS curves among HR+/HER2- breast cancer patients according to both *PRICKLE1* and *LL5β/PHLDB2* mRNA expression: the patients are separated into two classes (“high/high” *vs.* “no-high/high”). **C)** Similar to B, but among HER2+ breast cancer patients. **D)** Prognostic complementarity of expression of both genes in TNBC using the likelihood ratio (LR) test.
