## Supplementray Table1 for "The serine/threonine kinase MINK1 directly regulates the function of promigratory proteins"

**Supplementary Table 1:** List of phosphopeptides identified by SILAC approach

| **Protein** | **RatioHsurL** | **modified residue** | **Protein** | **Protein names** | **Position** | **Rank** | **Present in Prickle1-associated protein complex** |
| --- | --- | --- | --- | --- | --- | --- | --- |
| Q92614 | -2,9421 | Phosphoserine | Q92614 | Unconventional myosin-XVIIIa | 1970 | 1 |  |
| F8WBF5 | -1,868664 | Phosphoserine | F8WBF5 | Dual specificity protein kinase CLK1 | 140 | 2 |  |
| Q9Y385 | -1,798388 | Phosphoserine | Q9Y385 | Ubiquitin-conjugating enzyme E2 J1 | 266 | 3 |  |
| E9PFQ4 | -1,706317 | Phosphoserine | E9PFQ4 | Pleckstrin homology-like domain family B member 2 | 415 | 4 |  |
| Q7Z5L9 | -1,69797 | Phosphoserine | Q7Z5L9 | Interferon regulatory factor 2-binding protein 2 | 175 | 5 |  |
| E7EMC7 | -1,531637 | Phosphothreonine | E7EMC7 | Sequestosome-1 | 269 | 6 |  |
| Q96S55 | -1,521704 | Phosphoserine | Q96S55 | ATPase WRNIP1 | 75 | 7 | yes |
| Q9NR30 | -1,506817 | Phosphoserine | Q9NR30 | Nucleolar RNA helicase 2 | 89 | 8 |  |
| Q96B36 | -1,497238 | Phosphoserine | Q96B36 | Proline-rich AKT1 substrate 1 | 183 | 9 |  |
| Q7Z5L9 | -1,450542 | Phosphoserine | Q7Z5L9 | Interferon regulatory factor 2-binding protein 2 | 360 | 10 |  |
| Q8N4C8 | -1,374951 | Phosphoserine | Q8N4C8 | Misshapen-like kinase 1 | 763 | 11 | yes |
| Q92922 | -1,374527 | Phosphoserine | Q92922 | SWI/SNF complex subunit SMARCC1 | 310 | 12 |  |
| Q96B36 | -1,29368 | Phosphoserine | Q96B36 | Proline-rich AKT1 substrate 1 | 211 | 13 |  |
| Q14126 | -1,279174 | Phosphoserine | Q14126 | Desmoglein-2 | 680 | 14 |  |
| E7EMC7 | -1,251098 | Phosphoserine | E7EMC7 | Sequestosome-1 | 272 | 15 |  |
| F8WE35 | -1,246284 | Phosphoserine | F8WE35 | Activating transcription factor 7-interacting protein 1 | 113 | 16 |  |
| A0A0C4DGZ0 | -1,225123 | Phosphoserine | A0A0C4DGZ0 | DNA-directed RNA polymerase II subunit RPB1 | 1913 | 17 |  |
| Q7Z6Z7 | -1,213407 | Phosphoserine | Q7Z6Z7 | E3 ubiquitin-protein ligase HUWE1 | 3919 | 18 | yes |
| Q9ULW0 | -1,175162 | Phosphoserine | Q9ULW0 | Targeting protein for Xklp2 | 738 | 19 | yes |
| Q9H6F5 | -1,151814 | Phosphoserine | Q9H6F5 | Coiled-coil domain-containing protein 86 | 47 | 20 |  |
| P10412 | -1,1458 | Phosphothreonine | P10412 | Histone H1.4 | 18 | 21 |  |
| A0A087WZH7 | -1,145584 | Phosphoserine | A0A087WZH7 | Myristoylated alanine-rich C-kinase substrate | 27 | 22 |  |
| I3L2J0 | -1,144825 | Phosphoserine | I3L2J0 | Protein capicua homolog | 2303 | 23 |  |
| I3L2J0 | -1,144825 | Phosphoserine | I3L2J0 | Protein capicua homolog | 2315 | 24 |  |
| Q53EL6 | -1,143922 | Phosphoserine | Q53EL6 | Programmed cell death protein 4 | 94 | 25 |  |
| A0A087WXZ1 | -1,139884 | Phosphoserine\| by AURKA\| CDK1 and CDK2 | A0A087WXZ1 | Cellular tumor antigen p53 | 156 | 26 |  |
| A0A087WV66 | -1,135427 | Phosphoserine | A0A087WV66 | Antigen KI-67 | 1130 | 27 | yes |
| C9JZR2 | -1,113839 | Phosphoserine | C9JZR2 | Catenin delta-1 | 268 | 28 |  |
| A0A087WXM0 | -1,113238 | Phosphoserine | A0A087WXM0 | Forkhead box protein K2 | 394 | 29 |  |
| P51397 | -1,112213 | Phosphoserine\| by MTOR | P51397 | Death-associated protein 1 | 3 | 30 |  |
| Q92614 | -1,10858 | Phosphoserine | Q92614 | Unconventional myosin-XVIIIa | 2020 | 31 |  |
| P17096 | -1,107875 | Phosphothreonine\| by HIPK2 and CDC2 | P17096 | High mobility group protein HMG-I/HMG-Y | 53 | 32 |  |
| Q9Y2D5 | -1,070682 | Phosphoserine | Q9Y2D5 | A-kinase anchor protein 2 | 748 | 33 |  |
| O94762 | -1,054812 | Phosphoserine | O94762 | ATP-dependent DNA helicase Q5 | 815 | 34 |  |
| H0YEV2 | -1,048749 | Phosphothreonine | H0YEV2 | Src substrate cortactin | 35 | 35 |  |
| O95817 | -1,046925 | Phosphoserine | O95817 | BAG family molecular chaperone regulator 3 | 377 | 36 |  |
| C9JW01 | -1,044261 | Phosphoserine | C9JW01 | Pumilio homolog 2 | 136 | 37 |  |
| P60174 | -1,022956 | Phosphoserine | P60174 | Triosephosphate isomerase | 58 | 38 |  |
| E9PFQ4 | -1,01974 | Phosphothreonine | E9PFQ4 | Pleckstrin homology-like domain family B member 2 | 855 | 39 |  |
| Q9UJM3 | -1,019011 | Phosphoserine | Q9UJM3 | ERBB receptor feedback inhibitor 1 | 302 | 40 |  |
| B4E135 | -1,017191 | Phosphothreonine | B4E135 | DNA ligase;DNA ligase 1 | 163 | 41 |  |
| I3L0Y9 | -1,015506 | Phosphothreonine | I3L0Y9 | Brain-specific angiogenesis inhibitor 1-associated protein 2 | 97 | 42 |  |
| Q96TA1 | -1,006812 | Phosphoserine | Q96TA1 | Niban-like protein 1 | 665 | 43 |  |
| Q96TA1 | -1,006812 | Phosphoserine | Q96TA1 | Niban-like protein 1 | 681 | 44 |  |
| Q86V48 | -0,9925364 | Phosphoserine | Q86V48 | Leucine zipper protein 1 | 659 | 45 |  |
| K7ENR9 | -0,9854024 | Phosphoserine | K7ENR9 | DNA ligase 3 | 123 | 46 |  |
| P16949 | -0,9824296 | Phosphoserine\| by PKA | P16949 | Stathmin | 63 | 47 |  |
| B4E135 | -0,9669538 | Phosphoserine | B4E135 | DNA ligase;DNA ligase 1 | 76 | 48 |  |
| E9PPX2 | -0,9659641 | Phosphoserine | E9PPX2 | Fos-related antigen 1 | 229 | 49 |  |
| C9JQE8 | -0,9636045 | Phosphoserine | C9JQE8 | Nuclear receptor corepressor 2 | 956 | 50 |  |
| R4GN84 | -0,9531284 | Phosphoserine | R4GN84 | SH2B adapter protein 3 | 86 | 51 |  |
| Q96EB6 | -0,9522745 | Phosphoserine | Q96EB6 | NAD-dependent protein deacetylase sirtuin-1;SirtT1 75 kDa fragment | 14 | 52 |  |
| E7EPK0 | -0,9452736 | Phosphoserine | E7EPK0 | LIM and calponin homology domains-containing protein 1 | 70 | 53 |  |
| Q96B36 | -0,9431674 | Phosphoserine | Q96B36 | Proline-rich AKT1 substrate 1 | 212 | 54 |  |
| Q9BUA3 | -0,9426962 | Phosphoserine | Q9BUA3 | Uncharacterized protein C11orf84 | 308 | 55 |  |
| P17096 | -0,9410642 | Phosphoserine | P17096 | High mobility group protein HMG-I/HMG-Y | 49 | 56 |  |
| P02545 | -0,939434 | Phosphoserine | P02545 | Prelamin-A/C;Lamin-A/C | 392 | 57 |  |
| E9PFQ4 | -0,9320584 | Phosphoserine | E9PFQ4 | Pleckstrin homology-like domain family B member 2 | 513 | 58 |  |
| Q5ST79 | -0,9151687 | Phosphoserine | Q5ST79 | Radiation-inducible immediate-early gene IEX-1 | 31 | 59 |  |
| G5E972 | -0,9002723 | Phosphoserine | G5E972 | Lamina-associated polypeptide 2, isoforms beta/gamma;Thymopoietin;Thymopentin;Lamina-associated polypeptide 2, isoform alpha;Thymopoietin;Thymopentin | 66 | 60 |  |
| G5E972 | -0,9002723 | Phosphoserine | G5E972 | Lamina-associated polypeptide 2, isoforms beta/gamma;Thymopoietin;Thymopentin;Lamina-associated polypeptide 2, isoform alpha;Thymopoietin;Thymopentin | 67 | 61 |  |
| G5E972 | -0,9002723 | Phosphothreonine | G5E972 | Lamina-associated polypeptide 2, isoforms beta/gamma;Thymopoietin;Thymopentin;Lamina-associated polypeptide 2, isoform alpha;Thymopoietin;Thymopentin | 74 | 62 |  |
| Q5QPJ9 | -0,8811281 | Phosphoserine | Q5QPJ9 | Dolichol-phosphate mannosyltransferase subunit 1 | 9 | 63 |  |
| K7ENW5 | -0,8758418 | Phosphoserine | K7ENW5 | Thymidine kinase;Thymidine kinase, cytosolic | 13 | 64 |  |
| P22314 | -0,8730564 | Phosphoserine | P22314 | Ubiquitin-like modifier-activating enzyme 1 | 46 | 65 |  |
| Q9NQW6 | -0,8614331 | Phosphoserine | Q9NQW6 | Actin-binding protein anillin | 323 | 66 |  |
| P16949 | -0,8604242 | Phosphoserine\| by CDK1\| MAPK1 and MAPK3 | P16949 | Stathmin | 38 | 67 |  |
| A0A087WV66 | -0,8580236 | Phosphoserine | A0A087WV66 | Antigen KI-67 | 308 | 68 |  |
| F5GYF8 | -0,854947 | Phosphoserine | F5GYF8 | Signal-induced proliferation-associated 1-like protein 1 | 1003 | 69 |  |
| O14745 | -0,8519656 | Phosphoserine | O14745 | Na(+)/H(+) exchange regulatory cofactor NHE-RF1 | 290 | 70 |  |
| A0A087WZ13 | -0,8506393 | Phosphoserine | A0A087WZ13 | Ribonucleoprotein PTB-binding 1 | 14 | 71 |  |
| P42684 | -0,8493729 | Phosphoserine | P42684 | Abelson tyrosine-protein kinase 2 | 620 | 72 |  |
| P02545 | -0,8417397 | Phosphoserine | P02545 | Prelamin-A/C;Lamin-A/C | 390 | 73 |  |
| P02545 | -0,8417397 | Phosphoserine | P02545 | Prelamin-A/C;Lamin-A/C | 395 | 74 |  |
| P19634 | -0,8379967 | Phosphoserine | P19634 | Sodium/hydrogen exchanger 1 | 703 | 75 |  |
| P51397 | -0,8375294 | Phosphoserine\| by MTOR | P51397 | Death-associated protein 1 | 51 | 76 |  |
| O14745 | -0,8366831 | Phosphoserine | O14745 | Na(+)/H(+) exchange regulatory cofactor NHE-RF1 | 280 | 77 |  |
| Q13177 | -0,8326911 | Phosphoserine | Q13177 | Serine/threonine-protein kinase PAK 2;PAK-2p27;PAK-2p34 | 141 | 78 |  |
| A0A087WV66 | -0,8288553 | Phosphoserine | A0A087WV66 | Antigen KI-67 | 584 | 79 |  |
| C9JZR2 | -0,8178694 | Phosphoserine | C9JZR2 | Catenin delta-1 | 349 | 80 |  |
| C9JZR2 | -0,8178694 | Phosphoserine | C9JZR2 | Catenin delta-1 | 352 | 81 |  |
| P48634 | -0,8112319 | Phosphothreonine | P48634 | Protein PRRC2A | 610 | 82 |  |
| Q9H7N4 | -0,807567 | Phosphoserine | Q9H7N4 | Splicing factor, arginine/serine-rich 19 | 874 | 83 |  |
| P31350 | -0,8056238 | Phosphoserine | P31350 | Ribonucleoside-diphosphate reductase subunit M2 | 20 | 84 |  |
| C9JKH7 | -0,8014891 | Phosphoserine | C9JKH7 | Fibronectin type III domain-containing protein 3B | 181 | 85 |  |
| H0YKH0 | -0,8011188 | Phosphoserine | H0YKH0 | Transducin-like enhancer protein 3 | 286 | 86 |  |
| A0A0A0MSK6 | -0,7973661 | Phosphoserine | A0A0A0MSK6 | Rho GTPase-activating protein 5 | 1217 | 87 |  |
| J3QLL0 | -0,7971957 | Phosphoserine | J3QLL0 | Importin subunit alpha-1 | 62 | 88 |  |
| D6REC6 | -0,793906 | Phosphothreonine\| by CDK2 | D6REC6 | Cyclin-dependent kinase 7 | 77 | 89 |  |
| O75815 | -0,7938778 | Phosphoserine | O75815 | Breast cancer anti-estrogen resistance protein 3 | 83 | 90 |  |
| Q6WCQ1 | -0,7937078 | Phosphoserine | Q6WCQ1 | Myosin phosphatase Rho-interacting protein | 365 | 91 |  |
| H3BM18 | -0,7926886 | Phosphoserine | H3BM18 | Gamma-interferon-inducible protein 16 | 106 | 92 |  |
| E9PRB3 | -0,792547 | Phosphoserine | E9PRB3 | Deoxynucleotidyltransferase terminal-interacting protein 2 | 148 | 93 |  |
| Q53EZ4 | -0,7838852 | Phosphoserine\| by CDK1 and MAPK1 | Q53EZ4 | Centrosomal protein of 55 kDa | 428 | 94 |  |
| Q9P206 | -0,7782984 | Phosphoserine | Q9P206 | Uncharacterized protein KIAA1522 | 545 | 95 |  |
| Q9NZ63 | -0,7743248 | Phosphoserine | Q9NZ63 | Uncharacterized protein C9orf78 | 261 | 96 |  |
| Q8WUX9 | -0,7694985 | Phosphoserine | Q8WUX9 | Charged multivesicular body protein 7 | 417 | 97 |  |
| Q1RLN5 | -0,7692757 | Phosphoserine | Q1RLN5 | Rho GTPase-activating protein 12 | 240 | 98 |  |
| O75533 | -0,7657714 | Phosphothreonine | O75533 | Splicing factor 3B subunit 1 | 142 | 99 |  |
| P11388 | -0,7623586 | Phosphoserine | P11388 | DNA topoisomerase 2-alpha | 1393 | 100 |  |
| Q15059 | -0,760946 | Phosphoserine | Q15059 | Bromodomain-containing protein 3 | 263 | 101 |  |
| J3KSZ6 | -0,7563573 | Phosphoserine | J3KSZ6 | Telomeric repeat-binding factor 2 | 109 | 102 |  |
| Q02880 | -0,7552263 | Phosphoserine | Q02880 | DNA topoisomerase 2-beta | 1581 | 103 |  |
| C9J6P4 | -0,7544543 | Phosphoserine | C9J6P4 | Zinc finger CCCH-type antiviral protein 1 | 284 | 104 |  |
| O60343 | -0,7523336 | Phosphoserine\| by PKB/AKT1 | O60343 | TBC1 domain family member 4 | 318 | 105 |  |
| A0A0B4J210 | -0,7497214 | Phosphoserine | A0A0B4J210 | La-related protein 1 | 774 | 106 |  |
| H3BUZ5 | -0,7484308 | Phosphoserine | H3BUZ5 | Pseudopodium-enriched atypical kinase 1 | 281 | 107 |  |
| Q15773 | -0,746785 | Phosphoserine | Q15773 | Myeloid leukemia factor 2 | 238 | 108 |  |
| Q9H501 | -0,7439639 | Phosphoserine | Q9H501 | ESF1 homolog | 153 | 109 |  |
| A0A087WUZ3 | -0,7405204 | Phosphothreonine | A0A087WUZ3 | Spectrin beta chain, non-erythrocytic 1 | 2330 | 110 |  |
| Q8NFJ5 | -0,7398927 | Phosphoserine | Q8NFJ5 | Retinoic acid-induced protein 3 | 345 | 111 |  |
| H0Y589 | -0,7337395 | Phosphoserine | H0Y589 | Histone lysine demethylase PHF8 | 585 | 112 |  |
| Q6WCQ1 | -0,7324631 | Phosphoserine | Q6WCQ1 | Myosin phosphatase Rho-interacting protein | 993 | 113 |  |
| H3BS86 | -0,7291823 | Phosphoserine | H3BS86 | E3 ubiquitin-protein ligase CHIP | 19 | 114 |  |
| Q13541 | -0,726044 | Phosphothreonine\| by MTOR | Q13541 | Eukaryotic translation initiation factor 4E-binding protein 1 | 37 | 115 |  |
| O95365 | -0,7176369 | Phosphoserine | O95365 | Zinc finger and BTB domain-containing protein 7A | 525 | 116 |  |
| Q9H6F5 | -0,7167776 | Phosphoserine | Q9H6F5 | Coiled-coil domain-containing protein 86 | 91 | 117 |  |
| F5GX58 | -0,7152478 | Phosphoserine | F5GX58 | Cell division cycle-associated protein 3 | 68 | 118 |  |
| Q53EL6 | -0,7140144 | Phosphoserine\| by PKB | Q53EL6 | Programmed cell death protein 4 | 457 | 119 |  |
| C9JAT7 | -0,7034405 | Phosphoserine | C9JAT7 | UBX domain-containing protein 7 | 126 | 120 |  |
| A0A0B4J210 | -0,700756 | Phosphoserine | A0A0B4J210 | La-related protein 1 | 766 | 121 |  |
| G3V4T7 | -0,6997207 | Phosphoserine | G3V4T7 | FERM domain-containing protein 6 | 467 | 122 |  |
| Q9H1E3 | -0,6988188 | Phosphoserine | Q9H1E3 | Nuclear ubiquitous casein and cyclin-dependent kinase substrate 1 | 181 | 123 |  |
| Q9H1E3 | -0,6988188 | Phosphothreonine | Q9H1E3 | Nuclear ubiquitous casein and cyclin-dependent kinase substrate 1 | 179 | 124 |  |
| C9J0A7 | -0,6805047 | Phosphoserine | C9J0A7 | Charged multivesicular body protein 2b | 169 | 125 |  |
| Q02880 | -0,6797194 | Phosphothreonine | Q02880 | DNA topoisomerase 2-beta | 1575 | 126 |  |
| A0A087WZH7 | -0,674912 | Phosphoserine | A0A087WZH7 | Myristoylated alanine-rich C-kinase substrate | 101 | 127 |  |
| O75815 | -0,6740776 | Phosphoserine | O75815 | Breast cancer anti-estrogen resistance protein 3 | 375 | 128 |  |
| Q15136 | -0,6716554 | Phosphothreonine;Phosphothreonine\| by autocatalysis;Phosphothreonine\| by PDPK1 | Q15136 | cAMP-dependent protein kinase catalytic subunit beta;cAMP-dependent protein kinase catalytic subunit gamma;cAMP-dependent protein kinase catalytic subunit alpha | 181 | 129 |  |
| F8VPY7 | -0,6669526 | Phosphoserine | F8VPY7 | Periphilin-1 | 140 | 130 |  |
| Q96PU5 | -0,656897 | Phosphoserine\| by PKA and SGK1 | Q96PU5 | E3 ubiquitin-protein ligase NEDD4-like | 448 | 131 |  |
| A0A0A0MRN5 | -0,6563304 | Phosphoserine | A0A0A0MRN5 | Opioid growth factor receptor | 326 | 132 |  |
| Q8NFC6 | -0,6511897 | Phosphoserine | Q8NFC6 | Biorientation of chromosomes in cell division protein 1-like 1 | 1531 | 133 |  |
| Q96TA1 | -0,6505997 | Phosphoserine | Q96TA1 | Niban-like protein 1 | 665 | 134 |  |
| P49756 | -0,6387481 | Phosphoserine | P49756 | RNA-binding protein 25 | 677 | 135 |  |
| Q71RC2 | -0,6313646 | Phosphoserine | Q71RC2 | La-related protein 4 | 583 | 136 |  |
| P27824 | -0,6305045 | Phosphoserine | P27824 | Calnexin | 583 | 137 |  |
| E7EPK0 | -0,6278006 | Phosphoserine | E7EPK0 | LIM and calponin homology domains-containing protein 1 | 559 | 138 |  |
| Q96TA1 | -0,6247745 | Phosphoserine | Q96TA1 | Niban-like protein 1 | 692 | 139 |  |
| Q9H7N4 | -0,6173116 | Phosphoserine | Q9H7N4 | Splicing factor, arginine/serine-rich 19 | 498 | 140 |  |
| Q9H7N4 | -0,6173116 | Phosphoserine | Q9H7N4 | Splicing factor, arginine/serine-rich 19 | 500 | 141 |  |
| Q96S55 | -0,6168856 | Phosphothreonine | Q96S55 | ATPase WRNIP1 | 116 | 142 |  |
| Q9H6F5 | -0,6151078 | Phosphoserine | Q9H6F5 | Coiled-coil domain-containing protein 86 | 113 | 143 |  |
| Q13425 | -0,6138572 | Phosphoserine | Q13425 | Beta-2-syntrophin | 95 | 144 |  |
| F8WBF9 | -0,6117335 | Phosphoserine | F8WBF9 | Protein NDRG3 | 236 | 145 |  |
| Q9H1B7 | -0,607396 | Phosphoserine | Q9H1B7 | Interferon regulatory factor 2-binding protein-like | 547 | 146 |  |
| Q86WR7 | -0,6051824 | Phosphoserine | Q86WR7 | Proline and serine-rich protein 2 | 179 | 147 |  |
| I3L4N6 | -0,6049091 | Phosphoserine | I3L4N6 | Solute carrier family 12 member 4 | 919 | 148 |  |
| Q92922 | -0,6023768 | Phosphoserine | Q92922 | SWI/SNF complex subunit SMARCC1 | 330 | 149 |  |
| Q92922 | -0,6023768 | Phosphoserine | Q92922 | SWI/SNF complex subunit SMARCC1 | 328 | 150 |  |
| A6NFI3 | -0,6022528 | Phosphoserine | A6NFI3 | Zinc finger protein 316 | 10 | 151 |  |
| Q96T23 | -0,5976964 | Phosphoserine | Q96T23 | Remodeling and spacing factor 1 | 622 | 152 |  |
| P11717 | -0,5964113 | Phosphoserine | P11717 | Cation-independent mannose-6-phosphate receptor | 2409 | 153 |  |
| Q8IWS0 | -0,5937459 | Phosphoserine | Q8IWS0 | PHD finger protein 6 | 155 | 154 |  |
| C9J9W2 | -0,5925629 | Phosphothreonine | C9J9W2 | LIM and SH3 domain protein 1 | 68 | 155 |  |
| H0YM23 | -0,5907903 | Phosphoserine | H0YM23 | Ankyrin repeat domain-containing protein 17 | 2285 | 156 |  |
| Q86UP2 | -0,5882334 | Phosphoserine | Q86UP2 | Kinectin | 75 | 157 |  |
| O96013 | -0,5877177 | Phosphoserine | O96013 | Serine/threonine-protein kinase PAK 4 | 181 | 158 |  |
| Q9NYV4 | -0,5858529 | Phosphothreonine | Q9NYV4 | Cyclin-dependent kinase 12 | 893 | 159 |  |
| H0Y917 | -0,5857303 | Phosphoserine | H0Y917 | WD repeat-containing protein 26 | 113 | 160 |  |
| Q96E09 | -0,5702679 | Phosphoserine | Q96E09 | Protein FAM122A | 270 | 161 |  |
| F5GZP6 | -0,5696616 | Phosphoserine | F5GZP6 | Liprin-beta-1 | 410 | 162 |  |
| A0A0B4J210 | -0,5680378 | Phosphothreonine | A0A0B4J210 | La-related protein 1 | 526 | 163 |  |
| Q9H7N4 | -0,5665369 | Phosphoserine | Q9H7N4 | Splicing factor, arginine/serine-rich 19 | 239 | 164 |  |
| Q16204 | -0,56482 | Phosphoserine | Q16204 | Coiled-coil domain-containing protein 6 | 240 | 165 |  |
| Q16204 | -0,56482 | Phosphoserine | Q16204 | Coiled-coil domain-containing protein 6 | 244 | 166 |  |
| Q9C0B5 | -0,5639986 | Phosphoserine | Q9C0B5 | Palmitoyltransferase ZDHHC5 | 554 | 167 |  |
| H7C1L3 | -0,5618023 | Phosphoserine | H7C1L3 | ELM2 and SANT domain-containing protein 1 | 283 | 168 |  |
| F8WDB4 | -0,5611755 | Phosphoserine | F8WDB4 | Leucine-rich repeat and WD repeat-containing protein 1 | 60 | 169 |  |
| Q96N64 | -0,560838 | Phosphoserine | Q96N64 | PWWP domain-containing protein 2A | 81 | 170 |  |
| J3KSY7 | -0,5521639 | Phosphoserine | J3KSY7 | Protein CASC3 | 35 | 171 |  |
| P06400 | -0,549077 | Phosphothreonine | P06400 | Retinoblastoma-associated protein | 356 | 172 |  |
| Q8N3V7 | -0,5409483 | Phosphoserine | Q8N3V7 | Synaptopodin | 685 | 173 |  |
| Q13541 | -0,540877 | Phosphothreonine\| by MTOR | Q13541 | Eukaryotic translation initiation factor 4E-binding protein 1 | 46 | 174 |  |
| Q96TA1 | -0,540473 | Phosphoserine | Q96TA1 | Niban-like protein 1 | 696 | 175 |  |
| A0A0C4DGZ0 | -0,5395466 | Phosphoserine | A0A0C4DGZ0 | DNA-directed RNA polymerase II subunit RPB1 | 1878 | 176 |  |
| Q13263 | -0,5345209 | Phosphoserine | Q13263 | Transcription intermediary factor 1-beta | 19 | 177 |  |
| A1L390 | -0,5342607 | Phosphoserine | A1L390 | Pleckstrin homology domain-containing family G member 3 | 1037 | 178 |  |
| A1L390 | -0,5342607 | Phosphoserine | A1L390 | Pleckstrin homology domain-containing family G member 3 | 1040 | 179 |  |
| V9GYM8 | -0,5297958 | Phosphoserine\| by PAK1 and AURKA | V9GYM8 | Rho guanine nucleotide exchange factor 2 | 931 | 180 |  |
| Q96TA1 | -0,525768 | Phosphoserine | Q96TA1 | Niban-like protein 1 | 692 | 181 |  |
| Q15477 | -0,5241458 | Phosphoserine | Q15477 | Helicase SKI2W | 256 | 182 |  |
| Q02241 | -0,5239814 | Phosphoserine | Q02241 | Kinesin-like protein KIF23 | 902 | 183 |  |
| E3W994 | -0,5210713 | Phosphoserine | E3W994 | CLIP-associating protein 2 | 575 | 184 |  |
| Q7Z5R6 | -0,5174185 | Phosphoserine | Q7Z5R6 | Amyloid beta A4 precursor protein-binding family B member 1-interacting protein | 526 | 185 |  |
| F8WBF9 | -0,517208 | Phosphoserine | F8WBF9 | Protein NDRG3 | 239 | 186 |  |
| P46937 | -0,5169742 | Phosphoserine\| by LATS1 and LATS2 | P46937 | Yorkie homolog | 109 | 187 |  |
| C9JW01 | -0,5103499 | Phosphoserine | C9JW01 | Pumilio homolog 2 | 182 | 188 |  |
| Q6P1N0 | -0,5085824 | Phosphoserine | Q6P1N0 | Coiled-coil and C2 domain-containing protein 1A | 455 | 189 |  |
| I3L1L3 | -0,5073742 | Phosphoserine | I3L1L3 | Myb-binding protein 1A | 1187 | 190 |  |
| Q9P0K7 | -0,5054247 | Phosphothreonine | Q9P0K7 | Ankycorbin | 249 | 191 |  |
| Q8NE71 | -0,5051001 | Phosphoserine | Q8NE71 | ATP-binding cassette sub-family F member 1 | 22 | 192 |  |
| E9PCX8 | -0,4971458 | Phosphoserine | E9PCX8 | Tensin-3 | 435 | 193 |  |
| H7BYJ3 | -0,4933228 | Phosphoserine | H7BYJ3 | Citron Rho-interacting kinase | 68 | 194 |  |
| M0R0X3 | -0,492955 | Phosphoserine | M0R0X3 | Protein Smaug homolog 2 | 271 | 195 |  |
| J3KP36 | -0,4911857 | Phosphoserine | J3KP36 | WASH complex subunit FAM21C | 483 | 196 |  |
| A0A087WUV6 | -0,4905428 | Phosphoserine | A0A087WUV6 | AT-rich interactive domain-containing protein 1A | 313 | 197 |  |
| A0A087WZH7 | -0,4896249 | Phosphoserine | A0A087WZH7 | Myristoylated alanine-rich C-kinase substrate | 118 | 198 |  |
| H3BTX0 | -0,4871953 | Phosphoserine | H3BTX0 | PAXIP1-associated glutamate-rich protein 1 | 67 | 199 |  |
| E9PCX8 | -0,4839243 | Phosphoserine | E9PCX8 | Tensin-3 | 763 | 200 |  |
| Q9C0C2 | -0,4827366 | Phosphoserine | Q9C0C2 | 182 kDa tankyrase-1-binding protein | 893 | 201 |  |
| Q00587 | -0,4806833 | Phosphoserine | Q00587 | Cdc42 effector protein 1 | 121 | 202 |  |
| P17252 | -0,4776316 | Phosphothreonine\| by PDPK1 | P17252 | Protein kinase C alpha type;Protein kinase C beta type;Protein kinase C gamma type | 497 | 203 |  |
| Q14160 | -0,4766764 | Phosphoserine | Q14160 | Protein scribble homolog | 1348 | 204 |  |
| Q8NHV4 | -0,4745636 | Phosphoserine | Q8NHV4 | Protein NEDD1 | 516 | 205 |  |
| C9JWF0 | -0,4728847 | Phosphoserine | C9JWF0 | Structural maintenance of chromosomes protein 4;Structural maintenance of chromosomes protein | 22 | 206 |  |
| F8W726 | -0,471321 | Phosphoserine | F8W726 | Ubiquitin-associated protein 2-like | 465 | 207 |  |
| C9JZR2 | -0,4708228 | Phosphoserine | C9JZR2 | Catenin delta-1 | 230 | 208 |  |
| P54727 | -0,467566 | Phosphoserine | P54727 | UV excision repair protein RAD23 homolog B | 160 | 209 |  |
| Q8NE71 | -0,46734 | Phosphothreonine | Q8NE71 | ATP-binding cassette sub-family F member 1 | 108 | 210 |  |
| Q9NYV4 | -0,466008 | Phosphoserine | Q9NYV4 | Cyclin-dependent kinase 12 | 681 | 211 |  |
| Q9NYV4 | -0,466008 | Phosphoserine | Q9NYV4 | Cyclin-dependent kinase 12 | 685 | 212 |  |
| Q96E09 | -0,4641812 | Phosphoserine | Q96E09 | Protein FAM122A | 143 | 213 |  |
| Q96E09 | -0,4641812 | Phosphoserine | Q96E09 | Protein FAM122A | 147 | 214 |  |
| P02545 | -0,463798 | Phosphoserine | P02545 | Prelamin-A/C;Lamin-A/C | 390 | 215 |  |
| Q9NRY4 | -0,4627619 | Phosphoserine | Q9NRY4 | Rho GTPase-activating protein 35 | 1179 | 216 |  |
| Q16555 | -0,4613891 | Phosphothreonine | Q16555 | Dihydropyrimidinase-related protein 2 | 509 | 217 |  |
| O95391 | -0,4599726 | Phosphoserine | O95391 | Pre-mRNA-splicing factor SLU7 | 215 | 218 |  |
| O43290 | -0,4562693 | Phosphoserine | O43290 | U4/U6.U5 tri-snRNP-associated protein 1 | 448 | 219 |  |
| Q9H1E3 | -0,4552386 | Phosphoserine | Q9H1E3 | Nuclear ubiquitous casein and cyclin-dependent kinase substrate 1 | 214 | 220 |  |
| Q96T23 | -0,4543653 | Phosphoserine | Q96T23 | Remodeling and spacing factor 1 | 604 | 221 |  |
| F5GYF8 | -0,4520168 | Phosphoserine | F5GYF8 | Signal-induced proliferation-associated 1-like protein 1 | 1039 | 222 |  |
| A6NNN6 | -0,4510337 | Phosphoserine | A6NNN6 | Pericentriolar material 1 protein | 1676 | 223 |  |
| P17096 | -0,4493372 | Phosphoserine\| by CK | P17096 | High mobility group protein HMG-I/HMG-Y | 102 | 224 |  |
| P17096 | -0,4493372 | Phosphoserine\| by CK | P17096 | High mobility group protein HMG-I/HMG-Y | 103 | 225 |  |
| Q7Z3K3 | -0,445461 | Phosphoserine | Q7Z3K3 | Pogo transposable element with ZNF domain | 425 | 226 |  |
| Q14966 | -0,4451717 | Phosphoserine | Q14966 | Zinc finger protein 638 | 605 | 227 |  |
| Q96TA1 | -0,445105 | Phosphoserine | Q96TA1 | Niban-like protein 1 | 641 | 228 |  |
| Q96TA1 | -0,445105 | Phosphoserine | Q96TA1 | Niban-like protein 1 | 646 | 229 |  |
| O95425 | -0,4433711 | Phosphoserine | O95425 | Supervillin | 245 | 230 |  |
| G5E972 | -0,4414619 | Phosphoserine | G5E972 | Lamina-associated polypeptide 2, isoforms beta/gamma;Thymopoietin;Thymopentin;Lamina-associated polypeptide 2, isoform alpha;Thymopoietin;Thymopentin | 67 | 231 |  |
| Q14160 | -0,4409959 | Phosphoserine | Q14160 | Protein scribble homolog | 1475 | 232 |  |
| O60832 | -0,436766 | Phosphoserine | O60832 | H/ACA ribonucleoprotein complex subunit 4 | 451 | 233 |  |
| B3KR49 | -0,4362133 | Phosphothreonine\| by MAP2K1 and MAP2K2 | B3KR49 | Mitogen-activated protein kinase;Mitogen-activated protein kinase 3 | 88 | 234 |  |
| B3KR49 | -0,4362133 | Phosphotyrosine\| by MAP2K1 and MAP2K2 | B3KR49 | Mitogen-activated protein kinase;Mitogen-activated protein kinase 3 | 90 | 235 |  |
| Q9UJM3 | -0,4351745 | Phosphoserine | Q9UJM3 | ERBB receptor feedback inhibitor 1 | 273 | 236 |  |
| O14545 | -0,4333422 | Phosphoserine | O14545 | TRAF-type zinc finger domain-containing protein 1 | 327 | 237 |  |
| Q9UJM3 | -0,4312479 | Phosphothreonine | Q9UJM3 | ERBB receptor feedback inhibitor 1 | 131 | 238 |  |
| Q86YV5 | -0,4295526 | Phosphoserine | Q86YV5 | Tyrosine-protein kinase SgK223 | 745 | 239 |  |
| B9A053 | -0,42731 | Phosphoserine | B9A053 | pre-mRNA 3' end processing protein WDR33 | 7 | 240 |  |
| H0YJA2 | -0,4264314 | Phosphoserine | H0YJA2 | Zinc finger CCCH domain-containing protein 14 | 431 | 241 |  |
| E7ETY4 | -0,425312 | Phosphoserine | E7ETY4 | Serine/threonine-protein kinase MARK2 | 422 | 242 |  |
| Q9P2B4 | -0,4243032 | Phosphoserine | Q9P2B4 | CTTNBP2 N-terminal-like protein | 488 | 243 |  |
| Q96JM3 | -0,4210839 | Phosphoserine | Q96JM3 | Chromosome alignment-maintaining phosphoprotein 1 | 282 | 244 |  |
| P40222 | -0,4206902 | Phosphoserine | P40222 | Alpha-taxilin | 515 | 245 |  |
| G3V2D5 | -0,4196408 | Phosphoserine | G3V2D5 | Zinc finger protein 36, C3H1 type-like 1 | 32 | 246 |  |
| M0R2S2 | -0,4196191 | Phosphoserine | M0R2S2 | Epidermal growth factor receptor substrate 15-like 1 | 101 | 247 |  |
| H3BMG9 | -0,4192475 | Phosphoserine | H3BMG9 | Protein FAM65A | 310 | 248 |  |
| Q8NE71 | -0,4183521 | Phosphoserine\| by CK2 | Q8NE71 | ATP-binding cassette sub-family F member 1 | 109 | 249 |  |
| P02545 | -0,4172173 | Phosphoserine | P02545 | Prelamin-A/C;Lamin-A/C | 22 | 250 |  |
| P49327 | -0,4160831 | Phosphothreonine | P49327 | Fatty acid synthase;[Acyl-carrier-protein] S-acetyltransferase;[Acyl-carrier-protein] S-malonyltransferase;3-oxoacyl-[acyl-carrier-protein] synthase;3-oxoacyl-[acyl-carrier-protein] reductase;3-hydroxyacyl-[acyl-carrier-protein] dehydratase;Enoyl-[acyl-carrier-protein] reductase;Oleoyl-[acyl-carrier-protein] hydrolase | 2204 | 251 |  |
| Q14004 | -0,4142097 | Phosphoserine | Q14004 | Cyclin-dependent kinase 13 | 383 | 252 |  |
| A0A087WVV2 | -0,414079 | Phosphoserine | A0A087WVV2 | Ribosome-binding protein 1 | 801 | 253 |  |
| G5E972 | -0,4127734 | Phosphothreonine | G5E972 | Lamina-associated polypeptide 2, isoforms beta/gamma;Thymopoietin;Thymopentin;Lamina-associated polypeptide 2, isoform alpha;Thymopoietin;Thymopentin | 160 | 254 |  |
| A0A087WY61 | -0,4118385 | Phosphothreonine | A0A087WY61 | Nuclear mitotic apparatus protein 1 | 1984 | 255 |  |
| H3BRT8 | -0,4106001 | Phosphoserine | H3BRT8 | Dual specificity protein kinase CLK3 | 9 | 256 |  |
| K7EIH3 | -0,4083867 | Phosphoserine | K7EIH3 | Protein unc-13 homolog D | 150 | 257 |  |
| Q8N5F7 | -0,4083867 | Phosphoserine | Q8N5F7 | NF-kappa-B-activating protein | 149 | 258 |  |
| Q8TAD8 | -0,4071947 | Phosphoserine | Q8TAD8 | Smad nuclear-interacting protein 1 | 52 | 259 |  |
| Q8TAD8 | -0,4071947 | Phosphoserine | Q8TAD8 | Smad nuclear-interacting protein 1 | 54 | 260 |  |
| A0A0B4J210 | -0,4021337 | Phosphoserine | A0A0B4J210 | La-related protein 1 | 774 | 261 |  |
| I3L1L3 | -0,4019827 | Phosphoserine | I3L1L3 | Myb-binding protein 1A | 1083 | 262 |  |
| C9JME2 | -0,3981882 | Phosphoserine | C9JME2 | FERM, RhoGEF and pleckstrin domain-containing protein 1 | 427 | 263 |  |
| Q6NZI2 | -0,3968754 | Phosphoserine | Q6NZI2 | Polymerase I and transcript release factor | 167 | 264 |  |
| Q9H6F5 | -0,3967463 | Phosphoserine | Q9H6F5 | Coiled-coil domain-containing protein 86 | 18 | 265 |  |
| P49792 | -0,3965312 | Phosphoserine | P49792 | E3 SUMO-protein ligase RanBP2 | 1160 | 266 |  |
| Q504U8 | -0,3945112 | Phosphothreonine\| by PKD/PRKD1 | Q504U8 | Epidermal growth factor receptor | 648 | 267 |  |
| P49006 | -0,3932017 | Phosphoserine | P49006 | MARCKS-related protein | 22 | 268 |  |
| E9PI52 | -0,3918934 | Phosphoserine | E9PI52 | Arginine/serine-rich coiled-coil protein 2 | 32 | 269 |  |
| E7EVA0 | -0,3917219 | Phosphoserine | E7EVA0 | Microtubule-associated protein;Microtubule-associated protein 4 | 653 | 270 |  |
| P11388 | -0,3875055 | Phosphoserine | P11388 | DNA topoisomerase 2-alpha | 1377 | 271 |  |
| B4E135 | -0,3841966 | Phosphoserine | B4E135 | DNA ligase;DNA ligase 1 | 66 | 272 |  |
| C9K098 | -0,3834292 | Phosphoserine | C9K098 | Cytoplasmic protein NCK1 | 85 | 273 |  |
| Q6WCQ1 | -0,3833013 | Phosphoserine | Q6WCQ1 | Myosin phosphatase Rho-interacting protein | 289 | 274 |  |
| Q14966 | -0,382364 | Phosphoserine | Q14966 | Zinc finger protein 638 | 552 | 275 |  |
| E9PMS6 | -0,3799809 | Phosphoserine | E9PMS6 | LIM domain only protein 7 | 432 | 276 |  |
| P85037 | -0,3744424 | Phosphoserine | P85037 | Forkhead box protein K1 | 445 | 277 |  |
| J3KN59 | -0,373532 | Phosphoserine | J3KN59 | BCL2/adenovirus E1B 19 kDa protein-interacting protein 2 | 235 | 278 |  |
| Q96T23 | -0,3720723 | Phosphoserine | Q96T23 | Remodeling and spacing factor 1 | 1345 | 279 |  |
| Q9Y2D5 | -0,3716706 | Phosphoserine | Q9Y2D5 | A-kinase anchor protein 2 | 393 | 280 |  |
| Q15678 | -0,36844 | Phosphoserine | Q15678 | Tyrosine-protein phosphatase non-receptor type 14 | 594 | 281 |  |
| P46821 | -0,3670064 | Phosphoserine | P46821 | Microtubule-associated protein 1B;MAP1B heavy chain;MAP1 light chain LC1 | 1265 | 282 |  |
| Q8N556 | -0,3668379 | Phosphoserine | Q8N556 | Actin filament-associated protein 1 | 668 | 283 |  |
| A0A087X0K9 | -0,3649851 | Phosphoserine | A0A087X0K9 | Tight junction protein ZO-1 | 605 | 284 |  |
| J3KQ96 | -0,3642489 | Phosphoserine | J3KQ96 | Treacle protein | 1340 | 285 |  |
| K7EMU2 | -0,363576 | Phosphoserine | K7EMU2 | cAMP-dependent protein kinase type I-alpha regulatory subunit;cAMP-dependent protein kinase type I-alpha regulatory subunit, N-terminally processed | 83 | 286 |  |
| Q14573 | -0,3619164 | Phosphoserine | Q14573 | Inositol 1,4,5-trisphosphate receptor type 3 | 934 | 287 |  |
| A0A0A0MT33 | -0,3604684 | Phosphoserine | A0A0A0MT33 | Protein SCAF8 | 695 | 288 |  |
| Q13242 | -0,3601537 | Phosphoserine | Q13242 | Serine/arginine-rich splicing factor 9 | 211 | 289 |  |
| B4DP61 | -0,3595458 | Phosphoserine | B4DP61 | Survival motor neuron protein | 28 | 290 |  |
| B4DP61 | -0,3595458 | Phosphoserine | B4DP61 | Survival motor neuron protein | 31 | 291 |  |
| Q9UKJ3 | -0,3558403 | Phosphoserine | Q9UKJ3 | G patch domain-containing protein 8 | 1107 | 292 |  |
| A0A087WZH7 | -0,355464 | Phosphoserine | A0A087WZH7 | Myristoylated alanine-rich C-kinase substrate | 145 | 293 |  |
| Q3MII6 | -0,3553386 | Phosphoserine | Q3MII6 | TBC1 domain family member 25 | 506 | 294 |  |
| J3KRF7 | -0,3460883 | Phosphoserine | J3KRF7 | Tether containing UBX domain for GLUT4 | 107 | 295 |  |
| B8QGS9 | -0,3445939 | Phosphoserine | B8QGS9 | Plakophilin-2 | 151 | 296 |  |
| B8QGS9 | -0,3445939 | Phosphoserine | B8QGS9 | Plakophilin-2 | 155 | 297 |  |
| Q4KMP7 | -0,3440546 | Phosphoserine | Q4KMP7 | TBC1 domain family member 10B | 661 | 298 |  |
| K7ENL0 | -0,3432047 | Phosphoserine | K7ENL0 | Septin-9 | 15 | 299 |  |
| F8VQE1 | -0,3431424 | Phosphoserine | F8VQE1 | LIM domain and actin-binding protein 1 | 202 | 300 |  |
| A0A0A0MRM9 | -0,3423551 | Phosphoserine | A0A0A0MRM9 | Nucleolar and coiled-body phosphoprotein 1 | 707 | 301 |  |
| G3XAM7 | -0,3390448 | Phosphoserine | G3XAM7 | Catenin alpha-1 | 641 | 302 |  |
| Q52LW3 | -0,3355977 | Phosphoserine | Q52LW3 | Rho GTPase-activating protein 29 | 190 | 303 |  |
| Q9UQ35 | -0,3354327 | Phosphoserine | Q9UQ35 | Serine/arginine repetitive matrix protein 2 | 1387 | 304 |  |
| A0A087WY61 | -0,3354121 | Phosphoserine | A0A087WY61 | Nuclear mitotic apparatus protein 1 | 169 | 305 |  |
| A0A0B4J210 | -0,3330435 | Phosphoserine | A0A0B4J210 | La-related protein 1 | 90 | 306 |  |
| Q15007 | -0,3330435 | Phosphoserine | Q15007 | Pre-mRNA-splicing regulator WTAP | 306 | 307 |  |
| A0A0A0MR52 | -0,3324879 | Phosphoserine | A0A0A0MR52 | Eukaryotic translation initiation factor 4 gamma 1 | 1032 | 308 |  |
| Q9BRD0 | -0,33035 | Phosphoserine | Q9BRD0 | BUD13 homolog | 271 | 309 |  |
| Q9Y3T9 | -0,3299391 | Phosphoserine | Q9Y3T9 | Nucleolar complex protein 2 homolog | 49 | 310 |  |
| Q15648 | -0,3285844 | Phosphothreonine | Q15648 | Mediator of RNA polymerase II transcription subunit 1 | 1051 | 311 |  |
| E9PI52 | -0,322177 | Phosphoserine | E9PI52 | Arginine/serine-rich coiled-coil protein 2 | 17 | 312 |  |
| Q9UKA4 | -0,3207479 | Phosphoserine | Q9UKA4 | A-kinase anchor protein 11 | 1242 | 313 |  |
| E9PCX8 | -0,3196464 | Phosphoserine | E9PCX8 | Tensin-3 | 879 | 314 |  |
| A0A0C4DGH2 | -0,3186068 | Phosphoserine | A0A0C4DGH2 | Ras-related protein R-Ras2 | 185 | 315 |  |
| F8VZJ2 | -0,3184642 | Phosphoserine | F8VZJ2 | Nascent polypeptide-associated complex subunit alpha;Nascent polypeptide-associated complex subunit alpha, muscle-specific form | 87 | 316 |  |
| E9PER6 | -0,3177513 | Phosphoserine\| by autocatalysis | E9PER6 | Putative 3-phosphoinositide-dependent protein kinase 2;3-phosphoinositide-dependent protein kinase 1 | 214 | 317 |  |
| C9JZR2 | -0,3172424 | Phosphoserine | C9JZR2 | Catenin delta-1 | 349 | 318 |  |
| Q5VUA4 | -0,3152692 | Phosphoserine | Q5VUA4 | Zinc finger protein 318 | 136 | 319 |  |
| F8W038 | -0,3146797 | Phosphoserine | F8W038 | Chromatin complexes subunit BAP18 | 96 | 320 |  |
| E9PNJ0 | -0,313989 | Phosphoserine | E9PNJ0 | Mitochondrial fission regulator 1-like | 6 | 321 |  |
| A0A0B4J210 | -0,312568 | Phosphoserine | A0A0B4J210 | La-related protein 1 | 548 | 322 |  |
| Q9UDY2 | -0,309305 | Phosphoserine | Q9UDY2 | Tight junction protein ZO-2 | 986 | 323 |  |
| E9PR38 | -0,3063321 | Phosphoserine | E9PR38 | Pumilio homolog 1 | 467 | 324 |  |
| F6XM96 | -0,306211 | Phosphoserine | F6XM96 | Tuftelin-interacting protein 11 | 98 | 325 |  |
| O94804 | -0,3061302 | Phosphothreonine | O94804 | Serine/threonine-protein kinase 10 | 952 | 326 |  |
| C9JME2 | -0,3052618 | Phosphothreonine | C9JME2 | FERM, RhoGEF and pleckstrin domain-containing protein 1 | 24 | 327 |  |
| F5H836 | -0,3038695 | Phosphoserine | F5H836 | Paxillin | 86 | 328 |  |
| A0A0A0MSK5 | -0,3031034 | Phosphothreonine | A0A0A0MSK5 | Torsin-1A-interacting protein 1 | 99 | 329 |  |
| Q14738 | -0,3030026 | Phosphoserine | Q14738 | Serine/threonine-protein phosphatase 2A 56 kDa regulatory subunit delta isoform | 573 | 330 |  |
| O75379 | -0,3026398 | Phosphoserine | O75379 | Vesicle-associated membrane protein 4 | 30 | 331 |  |
| Q5JTD0 | -0,3015118 | Phosphoserine | Q5JTD0 | Tight junction-associated protein 1 | 300 | 332 |  |
| A0A0A0MTP7 | -0,3007267 | Phosphoserine | A0A0A0MTP7 | Protein SCAF11 | 548 | 333 |  |
| A0A0A0MRX3 | -0,2997611 | Phosphoserine | A0A0A0MRX3 | DENN domain-containing protein 4C | 8 | 334 |  |
| C9J542 | -0,2993992 | Phosphoserine | C9J542 | PDZ and LIM domain protein 4 | 53 | 335 |  |
| E5RIS7 | -0,2992986 | Phosphoserine | E5RIS7 | Transcription elongation factor A protein 1 | 100 | 336 |  |
| M0QXD6 | -0,2990774 | Phosphoserine | M0QXD6 | General transcription factor IIF subunit 1 | 307 | 337 |  |
| Q86U06 | -0,2985148 | Phosphoserine | Q86U06 | Probable RNA-binding protein 23 | 149 | 338 |  |
| Q32MZ4 | -0,2974703 | Phosphoserine | Q32MZ4 | Leucine-rich repeat flightless-interacting protein 1 | 714 | 339 |  |
| F8WAE5 | -0,2972092 | Phosphoserine | F8WAE5 | Eukaryotic translation initiation factor 2A;Eukaryotic translation initiation factor 2A, N-terminally processed | 501 | 340 |  |
| Q9BRD0 | -0,2891016 | Phosphoserine | Q9BRD0 | BUD13 homolog | 163 | 341 |  |
| Q9BRD0 | -0,2891016 | Phosphothreonine | Q9BRD0 | BUD13 homolog | 159 | 342 |  |
| P31323 | -0,2871864 | Phosphoserine | P31323 | cAMP-dependent protein kinase type II-beta regulatory subunit | 114 | 343 |  |
| E7EPN9 | -0,2851543 | Phosphoserine | E7EPN9 | Protein PRRC2C | 880 | 344 |  |
| Q7Z5K2 | -0,2849155 | Phosphoserine | Q7Z5K2 | Wings apart-like protein homolog | 223 | 345 |  |
| Q8N556 | -0,2845572 | Phosphoserine | Q8N556 | Actin filament-associated protein 1 | 665 | 346 |  |
| Q5JTD0 | -0,2844378 | Phosphoserine | Q5JTD0 | Tight junction-associated protein 1 | 545 | 347 |  |
| O95425 | -0,2840597 | Phosphoserine | O95425 | Supervillin | 221 | 348 |  |
| H0YAZ3 | -0,2839006 | Phosphoserine\| by MAPK1 | H0YAZ3 | Vinexin | 42 | 349 |  |
| F8VPD4 | -0,2821315 | Phosphoserine\| by RPS6KB1 and PKA | F8VPD4 | CAD protein;Glutamine-dependent carbamoyl-phosphate synthase;Aspartate carbamoyltransferase;Dihydroorotase | 1796 | 350 |  |
| Q68EM7 | -0,2819129 | Phosphoserine | Q68EM7 | Rho GTPase-activating protein 17 | 575 | 351 |  |
| O14545 | -0,281337 | Phosphoserine | O14545 | TRAF-type zinc finger domain-containing protein 1 | 415 | 352 |  |
| B7Z8D3 | -0,2798486 | Phosphoserine | B7Z8D3 | Proteasome activator complex subunit 3 | 24 | 353 |  |
| Q9Y217 | -0,2797693 | Phosphoserine | Q9Y217 | Myotubularin-related protein 6 | 561 | 354 |  |
| Q13112 | -0,2768962 | Phosphoserine | Q13112 | Chromatin assembly factor 1 subunit B | 409 | 355 |  |
| Q16659 | -0,2749971 | Phosphoserine\| by PAK1\| PAK2 and PAK3 | Q16659 | Mitogen-activated protein kinase 6 | 189 | 356 |  |
| A0A0A0MSR2 | -0,2729624 | Phosphoserine | A0A0A0MSR2 | Alpha-adducin | 358 | 357 |  |
| G5E972 | -0,2729228 | Phosphoserine | G5E972 | Lamina-associated polypeptide 2, isoforms beta/gamma;Thymopoietin;Thymopentin;Lamina-associated polypeptide 2, isoform alpha;Thymopoietin;Thymopentin | 66 | 358 |  |
| Q15637 | -0,271522 | Phosphoserine | Q15637 | Splicing factor 1 | 80 | 359 |  |
| A2AB15 | -0,2702999 | Phosphoserine | A2AB15 | Putative pre-mRNA-splicing factor ATP-dependent RNA helicase DHX16 | 100 | 360 |  |
| O95365 | -0,2692952 | Phosphoserine | O95365 | Zinc finger and BTB domain-containing protein 7A | 549 | 361 |  |
| Q15149 | -0,266325 | Phosphoserine | Q15149 | Plectin | 1435 | 362 |  |
| P46821 | -0,2653821 | Phosphoserine | P46821 | Microtubule-associated protein 1B;MAP1B heavy chain;MAP1 light chain LC1 | 1312 | 363 |  |
| Q09666 | -0,2632236 | Phosphoserine | Q09666 | Neuroblast differentiation-associated protein AHNAK | 5749 | 364 |  |
| Q9UQ35 | -0,2621455 | Phosphoserine | Q9UQ35 | Serine/arginine repetitive matrix protein 2 | 377 | 365 |  |
| Q8WX92 | -0,2605985 | Phosphoserine | Q8WX92 | Negative elongation factor B | 557 | 366 |  |
| Q9UIG0 | -0,2604029 | Phosphoserine | Q9UIG0 | Tyrosine-protein kinase BAZ1B | 1468 | 367 |  |
| O60292 | -0,2595421 | Phosphoserine | O60292 | Signal-induced proliferation-associated 1-like protein 3 | 1544 | 368 |  |
| E9PJ24 | -0,2587208 | Phosphoserine | E9PJ24 | PHD and RING finger domain-containing protein 1 | 911 | 369 |  |
| Q5T5U3 | -0,2586231 | Phosphoserine | Q5T5U3 | Rho GTPase-activating protein 21 | 923 | 370 |  |
| Q9UQ35 | -0,2573532 | Phosphoserine | Q9UQ35 | Serine/arginine repetitive matrix protein 2 | 1188 | 371 |  |
| G3V4T7 | -0,2569041 | Phosphoserine | G3V4T7 | FERM domain-containing protein 6 | 448 | 372 |  |
| P34932 | -0,2556357 | Phosphoserine | P34932 | Heat shock 70 kDa protein 4 | 76 | 373 |  |
| D6RCK3 | -0,2554797 | Phosphothreonine;Phosphothreonine\| by STK4/MST1 | D6RCK3 | MOB kinase activator 1A;MOB kinase activator 1B | 40 | 374 |  |
| H0Y9B7 | -0,2544854 | Phosphoserine | H0Y9B7 | LEM domain-containing protein 2 | 81 | 375 |  |
| Q9NYV4 | -0,2537839 | Phosphoserine | Q9NYV4 | Cyclin-dependent kinase 12 | 1083 | 376 |  |
| E9PFQ4 | -0,2517983 | Phosphoserine | E9PFQ4 | Pleckstrin homology-like domain family B member 2 | 468 | 377 |  |
| Q8NE71 | -0,2506122 | Phosphoserine | Q8NE71 | ATP-binding cassette sub-family F member 1 | 105 | 378 |  |
| H0Y2Y8 | -0,2482428 | Phosphoserine | H0Y2Y8 | Zyxin | 312 | 379 |  |
| Q7Z6Z7 | -0,2462454 | Phosphoserine | Q7Z6Z7 | E3 ubiquitin-protein ligase HUWE1 | 1907 | 380 |  |
| H0Y2Y8 | -0,2453929 | Phosphoserine | H0Y2Y8 | Zyxin | 227 | 381 |  |
| Q9Y2U5 | -0,2438831 | Phosphoserine | Q9Y2U5 | Mitogen-activated protein kinase kinase kinase 2 | 164 | 382 |  |
| E9PCX8 | -0,2431095 | Phosphotyrosine | E9PCX8 | Tensin-3 | 883 | 383 |  |
| H0Y449 | -0,2391316 | Phosphoserine | H0Y449 | Nuclease-sensitive element-binding protein 1 | 215 | 384 |  |
| A0A0A0MRN5 | -0,2379366 | Phosphoserine | A0A0A0MRN5 | Opioid growth factor receptor | 263 | 385 |  |
| Q96P48 | -0,2363575 | Phosphoserine | Q96P48 | Arf-GAP with Rho-GAP domain, ANK repeat and PH domain-containing protein 1 | 229 | 386 |  |
| Q6UX15 | -0,2362806 | Phosphoserine | Q6UX15 | Layilin | 299 | 387 |  |
| Q8WUF5 | -0,2329166 | Phosphoserine | Q8WUF5 | RelA-associated inhibitor | 113 | 388 |  |
| Q5QP22 | -0,2320911 | Phosphoserine | Q5QP22 | RNA-binding protein 39 | 97 | 389 |  |
| O95544 | -0,2317075 | Phosphoserine | O95544 | NAD kinase | 48 | 390 |  |
| Q9C0C2 | -0,2316882 | Phosphoserine | Q9C0C2 | 182 kDa tankyrase-1-binding protein | 672 | 391 |  |
| Q8IVF2 | -0,2316882 | Phosphoserine | Q8IVF2 | Protein AHNAK2 | 294 | 392 |  |
| Q9BU76 | -0,2306718 | Phosphoserine | Q9BU76 | Multiple myeloma tumor-associated protein 2 | 220 | 393 |  |
| K7EQH1 | -0,2290241 | Phosphoserine | K7EQH1 | Uncharacterized protein C18orf25 | 66 | 394 |  |
| Q6NYC8 | -0,2279905 | Phosphoserine | Q6NYC8 | Phostensin | 224 | 395 |  |
| K7EQI9 | -0,2270341 | Phosphoserine | K7EQI9 | Protein FAM134C | 223 | 396 |  |
| Q7Z6E9 | -0,2270341 | Phosphoserine | Q7Z6E9 | E3 ubiquitin-protein ligase RBBP6 | 1179 | 397 |  |
| E9PMG4 | -0,2246076 | Phosphoserine | E9PMG4 | Telomerase Cajal body protein 1 | 54 | 398 |  |
| Q7Z417 | -0,223806 | Phosphoserine | Q7Z417 | Nuclear fragile X mental retardation-interacting protein 2 | 629 | 399 |  |
| Q9NR30 | -0,2222424 | Phosphoserine | Q9NR30 | Nucleolar RNA helicase 2 | 121 | 400 |  |
| E9PNJ4 | -0,2217088 | Phosphoserine | E9PNJ4 | Stromal interaction molecule 1 | 402 | 401 |  |
| E9PGM9 | -0,2213469 | Phosphoserine | E9PGM9 | RNA-binding protein 6 | 893 | 402 |  |
| F5H0B0 | -0,2200522 | Phosphoserine | F5H0B0 | Tumor protein D52 | 176 | 403 |  |
| E9PH18 | -0,2198049 | Phosphoserine | E9PH18 | DnaJ homolog subfamily B member 6 | 162 | 404 |  |
| P38432 | -0,219063 | Phosphothreonine | P38432 | Coilin | 122 | 405 |  |
| P29317 | -0,2161943 | Phosphoserine | P29317 | Ephrin type-A receptor 2 | 901 | 406 |  |
| B7ZL14 | -0,2124413 | Phosphoserine | B7ZL14 | Formin-binding protein 1 | 296 | 407 |  |
| G5E9C7 | -0,2123468 | Phosphoserine | G5E9C7 | Dual specificity mitogen-activated protein kinase kinase 2 | 198 | 408 |  |
| Q9C0C2 | -0,2114573 | Phosphoserine | Q9C0C2 | 182 kDa tankyrase-1-binding protein | 494 | 409 |  |
| Q8TAD8 | -0,2112871 | Phosphoserine | Q8TAD8 | Smad nuclear-interacting protein 1 | 35 | 410 |  |
| J3QSV6 | -0,2094724 | Phosphoserine | J3QSV6 | Ribosomal L1 domain-containing protein 1 | 360 | 411 |  |
| Q9BU76 | -0,2069621 | Phosphothreonine | Q9BU76 | Multiple myeloma tumor-associated protein 2 | 215 | 412 |  |
| C9JWF0 | -0,20634 | Phosphoserine | C9JWF0 | Structural maintenance of chromosomes protein 4;Structural maintenance of chromosomes protein | 22 | 413 |  |
| Q14160 | -0,2063211 | Phosphoserine | Q14160 | Protein scribble homolog | 1448 | 414 |  |
| Q9BU76 | -0,2038728 | Phosphothreonine | Q9BU76 | Multiple myeloma tumor-associated protein 2 | 219 | 415 |  |
| F8VZJ2 | -0,2031014 | Phosphothreonine | F8VZJ2 | Nascent polypeptide-associated complex subunit alpha;Nascent polypeptide-associated complex subunit alpha, muscle-specific form | 82 | 416 |  |
| Q9BTU6 | -0,2015037 | Phosphoserine | Q9BTU6 | Phosphatidylinositol 4-kinase type 2-alpha | 47 | 417 |  |
| Q9BTU6 | -0,2015037 | Phosphoserine | Q9BTU6 | Phosphatidylinositol 4-kinase type 2-alpha | 51 | 418 |  |
| Q7Z6Z7 | -0,200527 | Phosphoserine | Q7Z6Z7 | E3 ubiquitin-protein ligase HUWE1 | 3662 | 419 |  |
| Q8NDC0 | -0,2004707 | Phosphoserine | Q8NDC0 | MAPK-interacting and spindle-stabilizing protein-like | 15 | 420 |  |
| O15014 | -0,1972079 | Phosphoserine | O15014 | Zinc finger protein 609 | 576 | 421 |  |
| O15014 | -0,1972079 | Phosphoserine | O15014 | Zinc finger protein 609 | 578 | 422 |  |
| A0A087WWA3 | -0,1953547 | Phosphoserine | A0A087WWA3 | Kinesin-like protein KIF1B | 1031 | 423 |  |
| F5GX58 | -0,194214 | Phosphoserine | F5GX58 | Cell division cycle-associated protein 3 | 87 | 424 |  |
| A0A087WUI1 | -0,1941206 | Phosphoserine | A0A087WUI1 | RNA-binding motif protein, X-linked 2 | 188 | 425 |  |
| Q5VUA4 | -0,1919727 | Phosphoserine | Q5VUA4 | Zinc finger protein 318 | 2101 | 426 |  |
| A0A096LNL9 | -0,1915623 | Phosphoserine | A0A096LNL9 | Transcriptional regulator ATRX | 609 | 427 |  |
| E7EXA6 | -0,1900144 | Phosphoserine | E7EXA6 | Chromosome transmission fidelity protein 18 homolog | 871 | 428 |  |
| F2Z2E2 | -0,1884311 | Phosphoserine | F2Z2E2 | Ras GTPase-activating-like protein IQGAP3 | 1381 | 429 |  |
| Q96IF1 | -0,1883938 | Phosphoserine | Q96IF1 | LIM domain-containing protein ajuba | 119 | 430 |  |
| Q8IVF2 | -0,186738 | Phosphoserine | Q8IVF2 | Protein AHNAK2 | 280 | 431 |  |
| Q92733 | -0,1860687 | Phosphoserine | Q92733 | Proline-rich protein PRCC | 157 | 432 |  |
| Q92733 | -0,1860687 | Phosphoserine | Q92733 | Proline-rich protein PRCC | 159 | 433 |  |
| Q9UKA4 | -0,1854183 | Phosphoserine | Q9UKA4 | A-kinase anchor protein 11 | 448 | 434 |  |
| A6NNN6 | -0,1850281 | Phosphoserine | A6NNN6 | Pericentriolar material 1 protein | 65 | 435 |  |
| A6NNN6 | -0,1850281 | Phosphoserine | A6NNN6 | Pericentriolar material 1 protein | 69 | 436 |  |
| A8MT37 | -0,1829494 | Phosphotyrosine | A8MT37 | Glycogen synthase kinase-3 beta;Glycogen synthase kinase-3 alpha | 197 | 437 |  |
| Q4KMP7 | -0,1816146 | Phosphoserine | Q4KMP7 | TBC1 domain family member 10B | 678 | 438 |  |
| Q6KC79 | -0,1807811 | Phosphoserine | Q6KC79 | Nipped-B-like protein | 2658 | 439 |  |
| H0Y911 | -0,1804293 | Phosphoserine | H0Y911 | Transforming acidic coiled-coil-containing protein 2 | 56 | 440 |  |
| H0Y911 | -0,1804293 | Phosphoserine | H0Y911 | Transforming acidic coiled-coil-containing protein 2 | 60 | 441 |  |
| A0A0A0MSK6 | -0,1777101 | Phosphoserine | A0A0A0MSK6 | Rho GTPase-activating protein 5 | 968 | 442 |  |
| E5RIU6 | -0,1766019 | Phosphothreonine;Phosphothreonine\| by PKMYT1 | E5RIU6 | Cyclin-dependent kinase 1;Cyclin-dependent kinase 2;Cyclin-dependent kinase 3 | 14 | 443 |  |
| P29317 | -0,1764911 | Phosphoserine\| by PKB | P29317 | Ephrin type-A receptor 2 | 897 | 444 |  |
| C9JWF0 | -0,1763065 | Phosphoserine | C9JWF0 | Structural maintenance of chromosomes protein 4;Structural maintenance of chromosomes protein | 28 | 445 |  |
| Q09666 | -0,1753652 | Phosphoserine | Q09666 | Neuroblast differentiation-associated protein AHNAK | 5448 | 446 |  |
| H0Y7H7 | -0,1746275 | Phosphoserine | H0Y7H7 | Dedicator of cytokinesis protein 4 | 1081 | 447 |  |
| P46821 | -0,1735951 | Phosphoserine | P46821 | Microtubule-associated protein 1B;MAP1B heavy chain;MAP1 light chain LC1 | 614 | 448 |  |
| A0A0C4DH09 | -0,1726005 | Phosphoserine | A0A0C4DH09 | Protein transport protein Sec16A | 477 | 449 |  |
| Q9ULX3 | -0,1720298 | Phosphoserine | Q9ULX3 | RNA-binding protein NOB1 | 201 | 450 |  |
| Q8IX90 | -0,1708521 | Phosphoserine | Q8IX90 | Spindle and kinetochore-associated protein 3 | 155 | 451 |  |
| Q9H7N4 | -0,1695837 | Phosphoserine | Q9H7N4 | Splicing factor, arginine/serine-rich 19 | 734 | 452 |  |
| Q9H7N4 | -0,1695837 | Phosphoserine | Q9H7N4 | Splicing factor, arginine/serine-rich 19 | 738 | 453 |  |
| E9PFQ4 | -0,1678573 | Phosphoserine | E9PFQ4 | Pleckstrin homology-like domain family B member 2 | 334 | 454 |  |
| Q8IZ21 | -0,167527 | Phosphoserine | Q8IZ21 | Phosphatase and actin regulator 4 | 118 | 455 |  |
| J3KQ96 | -0,1631115 | Phosphoserine | J3KQ96 | Treacle protein | 171 | 456 |  |
| E7EN95 | -0,1627823 | Phosphoserine | E7EN95 | Filamin-B | 1914 | 457 |  |
| Q9UMZ2 | -0,1615392 | Phosphoserine | Q9UMZ2 | Synergin gamma | 752 | 458 |  |
| Q86WR7 | -0,16132 | Phosphoserine | Q86WR7 | Proline and serine-rich protein 2 | 43 | 459 |  |
| C9JQV3 | -0,1612103 | Phosphoserine | C9JQV3 | Serine/threonine-protein kinase 11-interacting protein | 387 | 460 |  |
| C9J0A5 | -0,160772 | Phosphoserine | C9J0A5 | E3 ubiquitin-protein ligase BRE1A | 138 | 461 |  |
| H0Y3P2 | -0,1597313 | Phosphothreonine | H0Y3P2 | Eukaryotic translation initiation factor 4 gamma 2 | 470 | 462 |  |
| Q14258 | -0,1592021 | Phosphoserine | Q14258 | E3 ubiquitin/ISG15 ligase TRIM25 | 100 | 463 |  |
| E7ETY4 | -0,1583084 | Phosphoserine | E7ETY4 | Serine/threonine-protein kinase MARK2 | 452 | 464 |  |
| J3QT54 | -0,1581991 | Phosphothreonine | J3QT54 | Cleavage and polyadenylation specificity factor subunit 7 | 194 | 465 |  |
| Q9C0B5 | -0,1578344 | Phosphoserine | Q9C0B5 | Palmitoyltransferase ZDHHC5 | 694 | 466 |  |
| H0YEN2 | -0,1572149 | Phosphoserine | H0YEN2 | Serine/threonine-protein phosphatase 6 regulatory subunit 3 | 324 | 467 |  |
| J3QSV6 | -0,1572149 | Phosphothreonine | J3QSV6 | Ribosomal L1 domain-containing protein 1 | 357 | 468 |  |
| C9JCN8 | -0,1535212 | Phosphoserine | C9JCN8 | Fos-related antigen 2 | 191 | 469 |  |
| P29317 | -0,1531941 | Phosphoserine | P29317 | Ephrin type-A receptor 2 | 901 | 470 |  |
| Q09666 | -0,1530125 | Phosphoserine | Q09666 | Neuroblast differentiation-associated protein AHNAK | 5841 | 471 |  |
| O94826 | -0,1524675 | Phosphoserine | O94826 | Mitochondrial import receptor subunit TOM70 | 91 | 472 |  |
| O00629 | -0,1512877 | Phosphoserine | O00629 | Importin subunit alpha-3 | 60 | 473 |  |
| Q96KR1 | -0,1506528 | Phosphoserine | Q96KR1 | Zinc finger RNA-binding protein | 1054 | 474 |  |
| Q6PD62 | -0,15 | Phosphothreonine | Q6PD62 | RNA polymerase-associated protein CTR9 homolog | 925 | 475 |  |
| O95466 | -0,149855 | Phosphoserine | O95466 | Formin-like protein 1 | 624 | 476 |  |
| A0A0A0MQR1 | -0,1496376 | Phosphoserine | A0A0A0MQR1 | Mitogen-activated protein kinase kinase kinase kinase 5 | 335 | 477 |  |
| Q9Y6D5 | -0,1493476 | Phosphoserine | Q9Y6D5 | Brefeldin A-inhibited guanine nucleotide-exchange protein 2 | 227 | 478 |  |
| Q7Z6Z7 | -0,1481341 | Phosphoserine | Q7Z6Z7 | E3 ubiquitin-protein ligase HUWE1 | 1395 | 479 |  |
| B4DWJ1 | -0,1473739 | Phosphoserine | B4DWJ1 | Focal adhesion kinase 1 | 220 | 480 |  |
| Q92508 | -0,1461982 | Phosphoserine | Q92508 | Piezo-type mechanosensitive ion channel component 1 | 1646 | 481 |  |
| O00264 | -0,1458909 | Phosphoserine | O00264 | Membrane-associated progesterone receptor component 1 | 181 | 482 |  |
| Q5QP22 | -0,1454932 | Phosphoserine | Q5QP22 | RNA-binding protein 39 | 136 | 483 |  |
| H3BRM1 | -0,1440663 | Phosphoserine | H3BRM1 | Abscission/NoCut checkpoint regulator | 262 | 484 |  |
| Q14676 | -0,1429655 | Phosphoserine | Q14676 | Mediator of DNA damage checkpoint protein 1 | 168 | 485 |  |
| Q8IWZ8 | -0,1428031 | Phosphoserine | Q8IWZ8 | SURP and G-patch domain-containing protein 1 | 485 | 486 |  |
| A0A087WUZ3 | -0,1426588 | Phosphoserine | A0A087WUZ3 | Spectrin beta chain, non-erythrocytic 1 | 2343 | 487 |  |
| P05455 | -0,1419015 | Phosphoserine\| by CK2 | P05455 | Lupus La protein | 366 | 488 |  |
| I3L3A7 | -0,1409104 | Phosphoserine | I3L3A7 | Protein FAM195B | 21 | 489 |  |
| A0A087WUV8 | -0,1392901 | Phosphoserine | A0A087WUV8 | Basigin | 166 | 490 |  |
| P20290 | -0,1392001 | Phosphoserine | P20290 | Transcription factor BTF3 | 30 | 491 |  |
| E9PMS6 | -0,137366 | Phosphoserine | E9PMS6 | LIM domain only protein 7 | 553 | 492 |  |
| E9PHI6 | -0,1362882 | Phosphoserine | E9PHI6 | Cytoplasmic dynein 1 light intermediate chain 1 | 400 | 493 |  |
| P52756 | -0,1352651 | Phosphoserine | P52756 | RNA-binding protein 5 | 624 | 494 |  |
| F5H836 | -0,1335973 | Phosphoserine | F5H836 | Paxillin | 107 | 495 |  |
| A0A087WUH0 | -0,1319493 | Phosphoserine | A0A087WUH0 | DNA repair protein complementing XP-G cells | 384 | 496 |  |
| Q92538 | -0,1292664 | Phosphoserine | Q92538 | Golgi-specific brefeldin A-resistance guanine nucleotide exchange factor 1 | 1298 | 497 |  |
| E9PMS6 | -0,1290698 | Phosphoserine | E9PMS6 | LIM domain only protein 7 | 1185 | 498 |  |
| Q9BRD0 | -0,1287661 | Phosphoserine | Q9BRD0 | BUD13 homolog | 325 | 499 |  |
| F5GWT4 | -0,127427 | Phosphoserine | F5GWT4 | Serine/threonine-protein kinase WNK1 | 1784 | 500 |  |
| O14974 | -0,1270344 | Phosphothreonine\| by ROCK1\| ROCK2\| CDC42BP\| ZIPK/DAPK3 and RAF1 | O14974 | Protein phosphatase 1 regulatory subunit 12A | 696 | 501 |  |
| A0A087WUZ3 | -0,1263923 | Phosphoserine | A0A087WUZ3 | Spectrin beta chain, non-erythrocytic 1 | 2104 | 502 |  |
| A0A087WV73 | -0,1263388 | Phosphoserine | A0A087WV73 | Probable 28S rRNA (cytosine(4447)-C(5))-methyltransferase | 732 | 503 |  |
| P35606 | -0,1260535 | Phosphoserine | P35606 | Coatomer subunit beta' | 859 | 504 |  |
| A0A087WZH7 | -0,12559 | Phosphothreonine | A0A087WZH7 | Myristoylated alanine-rich C-kinase substrate | 150 | 505 |  |
| F5H777 | -0,1237197 | Phosphoserine | F5H777 | Coiled-coil domain-containing protein 82 | 131 | 506 |  |
| Q09666 | -0,1235773 | Phosphoserine | Q09666 | Neuroblast differentiation-associated protein AHNAK | 5763 | 507 |  |
| Q09666 | -0,1219051 | Phosphoserine | Q09666 | Neuroblast differentiation-associated protein AHNAK | 5763 | 508 |  |
| A0A0A0MR66 | -0,1213897 | Phosphoserine | A0A0A0MR66 | RNA-binding protein 10 | 788 | 509 |  |
| A0A0A0MSW3 | -0,1209454 | Phosphoserine | A0A0A0MSW3 | Nuclear pore complex protein Nup214 | 107 | 510 |  |
| Q5JW53 | -0,1200041 | Phosphoserine | Q5JW53 | Kinetochore-associated protein DSN1 homolog | 58 | 511 |  |
| Q09666 | -0,1199864 | Phosphoserine | Q09666 | Neuroblast differentiation-associated protein AHNAK | 5731 | 512 |  |
| P19338 | -0,1190634 | Phosphoserine | P19338 | Nucleolin | 563 | 513 |  |
| P49736 | -0,1187795 | Phosphoserine | P49736 | DNA replication licensing factor MCM2 | 139 | 514 |  |
| O00499 | -0,1185135 | Phosphoserine | O00499 | Myc box-dependent-interacting protein 1 | 331 | 515 |  |
| P16949 | -0,1165815 | Phosphoserine\| by CDK1\| MAPK1 and MAPK3 | P16949 | Stathmin | 25 | 516 |  |
| Q13283 | -0,1163336 | Phosphoserine | Q13283 | Ras GTPase-activating protein-binding protein 1 | 149 | 517 |  |
| P29317 | -0,115944 | Phosphoserine\| by PKB | P29317 | Ephrin type-A receptor 2 | 897 | 518 |  |
| O00505 | -0,1149175 | Phosphoserine | O00505 | Importin subunit alpha-4 | 60 | 519 |  |
| P18669 | -0,1145106 | Phosphoserine | P18669 | Phosphoglycerate mutase 1 | 14 | 520 |  |
| P50548 | -0,1132023 | Phosphothreonine\| by MAPK1 | P50548 | ETS domain-containing transcription factor ERF | 526 | 521 |  |
| E7EW05 | -0,1129372 | Phosphoserine | E7EW05 | Protein SDA1 homolog | 548 | 522 |  |
| P48634 | -0,1127782 | Phosphoserine | P48634 | Protein PRRC2A | 1219 | 523 |  |
| Q09666 | -0,1124956 | Phosphoserine | Q09666 | Neuroblast differentiation-associated protein AHNAK | 5752 | 524 |  |
| E9PS34 | -0,1117187 | Phosphoserine | E9PS34 | Nucleosome assembly protein 1-like 4 | 5 | 525 |  |
| P46821 | -0,110801 | Phosphoserine | P46821 | Microtubule-associated protein 1B;MAP1B heavy chain;MAP1 light chain LC1 | 1797 | 526 |  |
| Q8N655 | -0,1103776 | Phosphoserine | Q8N655 | Uncharacterized protein C10orf12 | 273 | 527 |  |
| P42684 | -0,1073649 | Phosphoserine | P42684 | Abelson tyrosine-protein kinase 2 | 631 | 528 |  |
| F6XM96 | -0,1072768 | Phosphoserine | F6XM96 | Tuftelin-interacting protein 11 | 59 | 529 |  |
| P46821 | -0,1061158 | Phosphoserine | P46821 | Microtubule-associated protein 1B;MAP1B heavy chain;MAP1 light chain LC1 | 1917 | 530 |  |
| E5RIU6 | -0,1060807 | Phosphothreonine;Phosphothreonine\| by PKMYT1 | E5RIU6 | Cyclin-dependent kinase 1;Cyclin-dependent kinase 2;Cyclin-dependent kinase 3 | 14 | 531 |  |
| E5RIU6 | -0,1060807 | Phosphotyrosine\| by PKMYT1\| WEE1\| WEE2 and PKC/PRKCD;Phosphotyrosine\| by WEE1 | E5RIU6 | Cyclin-dependent kinase 1;Cyclin-dependent kinase 2;Cyclin-dependent kinase 3 | 15 | 532 |  |
| Q13283 | -0,1054829 | Phosphoserine | Q13283 | Ras GTPase-activating protein-binding protein 1 | 232 | 533 |  |
| Q5VWU8 | -0,1040598 | Phosphoserine | Q5VWU8 | Partitioning defective 3 homolog | 383 | 534 |  |
| K7EQH1 | -0,1040248 | Phosphoserine | K7EQH1 | Uncharacterized protein C18orf25 | 66 | 535 |  |
| Q16643 | -0,1035858 | Phosphoserine | Q16643 | Drebrin | 142 | 536 |  |
| O43491 | -0,1022524 | Phosphoserine | O43491 | Band 4.1-like protein 2 | 715 | 537 |  |
| Q9UKM9 | -0,1015159 | Phosphoserine | Q9UKM9 | RNA-binding protein Raly | 135 | 538 |  |
| E7EPN9 | -0,1014458 | Phosphoserine | E7EPN9 | Protein PRRC2C | 2107 | 539 |  |
| G5E9C7 | -0,09965903 | Phosphothreonine | G5E9C7 | Dual specificity mitogen-activated protein kinase kinase 2 | 299 | 540 |  |
| H7C1M2 | -0,09562078 | Phosphoserine | H7C1M2 | Protein SON | 692 | 541 |  |
| Q6P2E9 | -0,09487033 | Phosphoserine | Q6P2E9 | Enhancer of mRNA-decapping protein 4 | 879 | 542 |  |
| Q9Y487 | -0,0940156 | Phosphoserine | Q9Y487 | V-type proton ATPase 116 kDa subunit a isoform 2 | 695 | 543 |  |
| H0Y9Z8 | -0,08828989 | Phosphoserine | H0Y9Z8 | Condensin complex subunit 3 | 239 | 544 |  |
| P04049 | -0,0877862 | Phosphoserine | P04049 | RAF proto-oncogene serine/threonine-protein kinase;Serine/threonine-protein kinase A-Raf | 621 | 545 |  |
| O14828 | -0,08757791 | Phosphoserine | O14828 | Secretory carrier-associated membrane protein 3 | 76 | 546 |  |
| E7EVA0 | -0,08563489 | Phosphoserine | E7EVA0 | Microtubule-associated protein;Microtubule-associated protein 4 | 2218 | 547 |  |
| E7EPK0 | -0,08490688 | Phosphoserine | E7EPK0 | LIM and calponin homology domains-containing protein 1 | 813 | 548 |  |
| A9Z1X7 | -0,08393684 | Phosphoserine | A9Z1X7 | Serine/arginine repetitive matrix protein 1 | 635 | 549 |  |
| A9Z1X7 | -0,08393684 | Phosphoserine | A9Z1X7 | Serine/arginine repetitive matrix protein 1 | 637 | 550 |  |
| Q16512 | -0,08379835 | Phosphoserine | Q16512 | Serine/threonine-protein kinase N1 | 916 | 551 |  |
| E9PMG4 | -0,08376366 | Phosphothreonine | E9PMG4 | Telomerase Cajal body protein 1 | 456 | 552 |  |
| Q3KQU3 | -0,08270788 | Phosphoserine | Q3KQU3 | MAP7 domain-containing protein 1 | 460 | 553 |  |
| H0Y7L2 | -0,08208513 | Phosphoserine | H0Y7L2 | Dedicator of cytokinesis protein 7 | 72 | 554 |  |
| P49736 | -0,08090964 | Phosphoserine | P49736 | DNA replication licensing factor MCM2 | 41 | 555 |  |
| A0A087WV57 | -0,08058131 | Phosphoserine | A0A087WV57 | Protein kinase C-binding protein 1 | 643 | 556 |  |
| Q9Y4E8 | -0,08042583 | Phosphoserine | Q9Y4E8 | Ubiquitin carboxyl-terminal hydrolase 15 | 229 | 557 |  |
| P35568 | -0,07994229 | Phosphoserine\| by RPS6KB1 and PKC/PRKCQ | P35568 | Insulin receptor substrate 1 | 1101 | 558 |  |
| A0A087WTQ7 | -0,07961419 | Phosphoserine | A0A087WTQ7 | DNA damage-binding protein 2 | 26 | 559 |  |
| Q9Y4H2 | -0,07847515 | Phosphoserine | Q9Y4H2 | Insulin receptor substrate 2 | 915 | 560 |  |
| Q7Z5K2 | -0,07781979 | Phosphoserine | Q7Z5K2 | Wings apart-like protein homolog | 221 | 561 |  |
| Q9Y2D5 | -0,07432362 | Phosphoserine | Q9Y2D5 | A-kinase anchor protein 2 | 152 | 562 |  |
| Q9H4L5 | -0,07420325 | Phosphoserine | Q9H4L5 | Oxysterol-binding protein-related protein 3 | 304 | 563 |  |
| E9PIE9 | -0,0728966 | Phosphoserine | E9PIE9 | G-protein-signaling modulator 3 | 39 | 564 |  |
| P05412 | -0,07262161 | Phosphoserine\| by MAPK8 and PLK3 | P05412 | Transcription factor AP-1;Transcription factor jun-D | 73 | 565 |  |
| A0A087WUZ3 | -0,07081872 | Phosphoserine | A0A087WUZ3 | Spectrin beta chain, non-erythrocytic 1 | 2140 | 566 |  |
| H3BRN3 | -0,07049279 | Phosphoserine\| by CDK1 and CDK2 | H3BRN3 | Protein PML | 68 | 567 |  |
| H3BRN3 | -0,07049279 | Phosphoserine\| by MAPK1 | H3BRN3 | Protein PML | 77 | 568 |  |
| A0A0D9SF54 | -0,07047561 | Phosphoserine | A0A0D9SF54 | Spectrin alpha chain, non-erythrocytic 1 | 1197 | 569 |  |
| Q9UQ35 | -0,06975529 | Phosphothreonine | Q9UQ35 | Serine/arginine repetitive matrix protein 2 | 1531 | 570 |  |
| Q96C19 | -0,06864107 | Phosphoserine | Q96C19 | EF-hand domain-containing protein D2 | 74 | 571 |  |
| A6NNN6 | -0,06804152 | Phosphoserine | A6NNN6 | Pericentriolar material 1 protein | 65 | 572 |  |
| E9PJ24 | -0,06754497 | Phosphoserine | E9PJ24 | PHD and RING finger domain-containing protein 1 | 1198 | 573 |  |
| F8WBF9 | -0,06742516 | Phosphoserine | F8WBF9 | Protein NDRG3 | 240 | 574 |  |
| Q92733 | -0,06621017 | Phosphoserine | Q92733 | Proline-rich protein PRCC | 267 | 575 |  |
| Q5T200 | -0,06573127 | Phosphoserine | Q5T200 | Zinc finger CCCH domain-containing protein 13 | 198 | 576 |  |
| P11388 | -0,06566288 | Phosphoserine | P11388 | DNA topoisomerase 2-alpha | 1247 | 577 |  |
| A0A0B4J210 | -0,06515002 | Phosphoserine | A0A0B4J210 | La-related protein 1 | 766 | 578 |  |
| K7EMR0 | -0,06441526 | Phosphoserine | K7EMR0 | Serine/threonine-protein kinase STK11 | 31 | 579 |  |
| Q9C0B5 | -0,06429562 | Phosphoserine | Q9C0B5 | Palmitoyltransferase ZDHHC5 | 621 | 580 |  |
| E9PHI6 | -0,0634588 | Phosphoserine | E9PHI6 | Cytoplasmic dynein 1 light intermediate chain 1 | 394 | 581 |  |
| P08670 | -0,06245193 | Phosphoserine | P08670 | Vimentin | 214 | 582 |  |
| F8W860 | -0,05963962 | Phosphoserine | F8W860 | Phosphatidylinositol 4-kinase beta | 96 | 583 |  |
| G3V340 | -0,05963962 | Phosphoserine | G3V340 | Striatin-3 | 229 | 584 |  |
| Q16204 | -0,05938429 | Phosphoserine | Q16204 | Coiled-coil domain-containing protein 6 | 244 | 585 |  |
| Q9H307 | -0,05761488 | Phosphoserine | Q9H307 | Pinin | 100 | 586 |  |
| V9GYM8 | -0,05683288 | Phosphoserine | V9GYM8 | Rho guanine nucleotide exchange factor 2 | 977 | 587 |  |
| H3BUY5 | -0,05477807 | Phosphoserine | H3BUY5 | RNA binding motif protein, X-linked-like-1;RNA-binding motif protein, X chromosome;RNA-binding motif protein, X chromosome, N-terminally processed | 80 | 588 |  |
| E5RJ37 | -0,05428597 | Phosphoserine | E5RJ37 | Tyrosine-protein kinase Lyn | 13 | 589 |  |
| G5E9C7 | -0,05425209 | Phosphothreonine | G5E9C7 | Dual specificity mitogen-activated protein kinase kinase 2 | 297 | 590 |  |
| Q5JWB9 | -0,0539128 | Phosphoserine | Q5JWB9 | Transmembrane protein 230 | 24 | 591 |  |
| H0YC41 | -0,05270913 | Phosphoserine | H0YC41 | Protein virilizer homolog | 12 | 592 |  |
| Q9BTE3 | -0,05260752 | Phosphoserine | Q9BTE3 | Mini-chromosome maintenance complex-binding protein | 154 | 593 |  |
| A0A0A0MR52 | -0,04961142 | Phosphoserine\| by PKC/PRKCA | A0A0A0MR52 | Eukaryotic translation initiation factor 4 gamma 1 | 986 | 594 |  |
| P26368 | -0,04930706 | Phosphoserine | P26368 | Splicing factor U2AF 65 kDa subunit | 79 | 595 |  |
| Q9UQ35 | -0,04792148 | Phosphoserine | Q9UQ35 | Serine/arginine repetitive matrix protein 2 | 323 | 596 |  |
| B5MDQ0 | -0,04746552 | Phosphoserine | B5MDQ0 | DNA excision repair protein ERCC-6-like | 946 | 597 |  |
| H0YEV2 | -0,04699279 | Phosphoserine | H0YEV2 | Src substrate cortactin | 39 | 598 |  |
| Q9Y2X3 | -0,04687469 | Phosphoserine | Q9Y2X3 | Nucleolar protein 58 | 514 | 599 |  |
| Q7Z6E9 | -0,04679029 | Phosphoserine | Q7Z6E9 | E3 ubiquitin-protein ligase RBBP6 | 861 | 600 |  |
| A2AB15 | -0,04574433 | Phosphoserine | A2AB15 | Putative pre-mRNA-splicing factor ATP-dependent RNA helicase DHX16 | 43 | 601 |  |
| E9PL53 | -0,04481713 | Phosphoserine | E9PL53 | Protein SAAL1 | 6 | 602 |  |
| J3KQ96 | -0,04267836 | Phosphoserine | J3KQ96 | Treacle protein | 1190 | 603 |  |
| Q13242 | -0,04227449 | Phosphoserine | Q13242 | Serine/arginine-rich splicing factor 9 | 204 | 604 |  |
| Q9Y3Z3 | -0,04116455 | Phosphothreonine | Q9Y3Z3 | Deoxynucleoside triphosphate triphosphohydrolase SAMHD1 | 592 | 605 |  |
| Q9Y2X3 | -0,0409797 | Phosphoserine | Q9Y2X3 | Nucleolar protein 58 | 502 | 606 |  |
| E9PS34 | -0,03965244 | Phosphoserine | E9PS34 | Nucleosome assembly protein 1-like 4 | 7 | 607 |  |
| K7ESQ0 | -0,03916545 | Phosphoserine | K7ESQ0 | Ran-binding protein 3 | 52 | 608 |  |
| H0YE74 | -0,03899767 | Phosphoserine | H0YE74 | Cleavage stimulation factor subunit 3 | 65 | 609 |  |
| J3KNN7 | -0,0388633 | Phosphoserine | J3KNN7 | BRCA1-associated protein | 89 | 610 |  |
| P17096 | -0,03861162 | Phosphoserine | P17096 | High mobility group protein HMG-I/HMG-Y | 44 | 611 |  |
| Q13439 | -0,03830959 | Phosphoserine | Q13439 | Golgin subfamily A member 4 | 71 | 612 |  |
| E7EVA0 | -0,03715244 | Phosphothreonine | E7EVA0 | Microtubule-associated protein;Microtubule-associated protein 4 | 371 | 613 |  |
| H0YEV2 | -0,03696799 | Phosphothreonine | H0YEV2 | Src substrate cortactin | 35 | 614 |  |
| E7ERS3 | -0,03646524 | Phosphoserine | E7ERS3 | Zinc finger CCCH domain-containing protein 18 | 892 | 615 |  |
| Q96A49 | -0,0361302 | Phosphothreonine | Q96A49 | Synapse-associated protein 1 | 248 | 616 |  |
| A2AB15 | -0,03417146 | Phosphoserine | A2AB15 | Putative pre-mRNA-splicing factor ATP-dependent RNA helicase DHX16 | 46 | 617 |  |
| P46821 | -0,03373657 | Phosphoserine | P46821 | Microtubule-associated protein 1B;MAP1B heavy chain;MAP1 light chain LC1 | 1396 | 618 |  |
| P46821 | -0,03373657 | Phosphoserine | P46821 | Microtubule-associated protein 1B;MAP1B heavy chain;MAP1 light chain LC1 | 1400 | 619 |  |
| D6REM6 | -0,03273349 | Phosphoserine | D6REM6 | Matrin-3 | 598 | 620 |  |
| D6REM6 | -0,03273349 | Phosphoserine | D6REM6 | Matrin-3 | 604 | 621 |  |
| A0A087WVV2 | -0,0301453 | Phosphoserine | A0A087WVV2 | Ribosome-binding protein 1 | 139 | 622 |  |
| O60885 | -0,02824472 | Phosphoserine | O60885 | Bromodomain-containing protein 4 | 1117 | 623 |  |
| Q08AD1 | -0,02776168 | Phosphoserine | Q08AD1 | Calmodulin-regulated spectrin-associated protein 2 | 599 | 624 |  |
| I3L436 | -0,02727869 | Phosphoserine\| by MAPK8 and MTOR | I3L436 | Regulatory-associated protein of mTOR | 23 | 625 |  |
| H7BXG7 | -0,02726212 | Phosphoserine | H7BXG7 | Protein transport protein Sec31A | 532 | 626 |  |
| E9PI52 | -0,02679597 | Phosphoserine | E9PI52 | Arginine/serine-rich coiled-coil protein 2 | 4 | 627 |  |
| Q9UG54 | -0,02594741 | Phosphoserine | Q9UG54 | Mitogen-activated protein kinase kinase kinase 7 | 93 | 628 |  |
| P49736 | -0,0248998 | Phosphoserine | P49736 | DNA replication licensing factor MCM2 | 27 | 629 |  |
| P61978 | -0,02179481 | Phosphoserine | P61978 | Heterogeneous nuclear ribonucleoprotein K | 116 | 630 |  |
| H7C559 | -0,02119781 | Phosphoserine | H7C559 | Serine/threonine-protein phosphatase 4 regulatory subunit 2 | 58 | 631 |  |
| Q09666 | -0,01920958 | Phosphoserine | Q09666 | Neuroblast differentiation-associated protein AHNAK | 216 | 632 |  |
| F8VQE1 | -0,01821655 | Phosphoserine | F8VQE1 | LIM domain and actin-binding protein 1 | 329 | 633 |  |
| F5H1N9 | -0,01752175 | Phosphoserine | F5H1N9 | Probable ATP-dependent RNA helicase DDX47 | 9 | 634 |  |
| F5GWT4 | -0,01704228 | Phosphoserine | F5GWT4 | Serine/threonine-protein kinase WNK1 | 1784 | 635 |  |
| A0A0D9SEV0 | -0,01626551 | Phosphothreonine | A0A0D9SEV0 | Bifunctional polynucleotide phosphatase/kinase;Polynucleotide 3'-phosphatase;Polynucleotide 5'-hydroxyl-kinase | 118 | 636 |  |
| E9PQN2 | -0,01532401 | Phosphoserine | E9PQN2 | Bcl-2-associated transcription factor 1 | 295 | 637 |  |
| Q9Y6D5 | -0,01530747 | Phosphoserine | Q9Y6D5 | Brefeldin A-inhibited guanine nucleotide-exchange protein 2 | 218 | 638 |  |
| Q9Y6D5 | -0,01530747 | Phosphoserine | Q9Y6D5 | Brefeldin A-inhibited guanine nucleotide-exchange protein 2 | 227 | 639 |  |
| P52948 | -0,01443259 | Phosphoserine | P52948 | Nuclear pore complex protein Nup98-Nup96;Nuclear pore complex protein Nup98;Nuclear pore complex protein Nup96 | 623 | 640 |  |
| A9Z1X7 | -0,01413558 | Phosphoserine | A9Z1X7 | Serine/arginine repetitive matrix protein 1 | 625 | 641 |  |
| E9PMS6 | -0,01372319 | Phosphoserine | E9PMS6 | LIM domain only protein 7 | 553 | 642 |  |
| E9PQN2 | -0,01362421 | Phosphoserine | E9PQN2 | Bcl-2-associated transcription factor 1 | 283 | 643 |  |
| E9PQN2 | -0,01362421 | Phosphoserine | E9PQN2 | Bcl-2-associated transcription factor 1 | 288 | 644 |  |
| J3KMZ8 | -0,01296464 | Phosphoserine | J3KMZ8 | Zinc finger protein ubi-d4 | 142 | 645 |  |
| F5H7V1 | -0,01258549 | Phosphoserine | F5H7V1 | Nardilysin | 94 | 646 |  |
| J3KQ96 | -0,01223946 | Phosphoserine | J3KQ96 | Treacle protein | 1312 | 647 |  |
| O43896 | -0,01210766 | Phosphothreonine | O43896 | Kinesin-like protein KIF1C | 1083 | 648 |  |
| Q9BTC0 | -0,01077372 | Phosphoserine | Q9BTC0 | Death-inducer obliterator 1 | 1456 | 649 |  |
| Q9BZQ8 | -0,01023068 | Phosphoserine | Q9BZQ8 | Protein Niban | 602 | 650 |  |
| J3QRU8 | -0,01018135 | Phosphoserine | J3QRU8 | ARF GTPase-activating protein GIT1 | 592 | 651 |  |
| E7ERY9 | -0,01011555 | Phosphoserine | E7ERY9 | Calcium-transporting ATPase;Plasma membrane calcium-transporting ATPase 1 | 898 | 652 |  |
| H0Y564 | -0,00965489 | Phosphoserine | H0Y564 | Anaphase-promoting complex subunit 1 | 223 | 653 |  |
| Q09666 | -0,00930956 | Phosphoserine | Q09666 | Neuroblast differentiation-associated protein AHNAK | 5110 | 654 |  |
| A0A087WV66 | -0,00896432 | Phosphoserine | A0A087WV66 | Antigen KI-67 | 357 | 655 |  |
| H0YEV2 | -0,00894785 | Phosphoserine | H0YEV2 | Src substrate cortactin | 51 | 656 |  |
| G3V5E1 | -0,00853686 | Phosphoserine | G3V5E1 | Cyclin-K | 340 | 657 |  |
| Q09666 | -0,00755113 | Phosphothreonine | Q09666 | Neuroblast differentiation-associated protein AHNAK | 4766 | 658 |  |
| Q9Y4H2 | -0,00676303 | Phosphoserine | Q9Y4H2 | Insulin receptor substrate 2 | 365 | 659 |  |
| V9GYM8 | -0,00594249 | Phosphoserine | V9GYM8 | Rho guanine nucleotide exchange factor 2 | 741 | 660 |  |
| K7EN42 | -0,00569649 | Phosphoserine | K7EN42 | Glycylpeptide N-tetradecanoyltransferase 1 | 46 | 661 |  |
| B0QY95 | -0,00558162 | Phosphoserine | B0QY95 | Mitochondrial dynamics protein MID51 | 55 | 662 |  |
| Q9UDY2 | -0,00525367 | Phosphoserine | Q9UDY2 | Tight junction protein ZO-2 | 130 | 663 |  |
| Q15149 | -0,00497496 | Phosphoserine | Q15149 | Plectin | 4386 | 664 |  |
| Q15149 | -0,00497496 | Phosphoserine | Q15149 | Plectin | 4389 | 665 |  |
| B7ZKW8 | -0,00433573 | Phosphoserine | B7ZKW8 | CapZ-interacting protein | 238 | 666 |  |
| Q13425 | -0,0042702 | Phosphoserine | Q13425 | Beta-2-syntrophin | 110 | 667 |  |
| Q8IVF2 | -0,00336918 | Phosphoserine | Q8IVF2 | Protein AHNAK2 | 1112 | 668 |  |
| H0YFD2 | -0,00273067 | Phosphoserine | H0YFD2 | Protein KRI1 homolog | 115 | 669 |  |
| E9PQA1 | -0,00245245 | Phosphoserine | E9PQA1 | Small acidic protein | 17 | 670 |  |
| Q9UQ35 | -0,00089858 | Phosphoserine | Q9UQ35 | Serine/arginine repetitive matrix protein 2 | 1014 | 671 |  |
| Q9UQ35 | -0,00089858 | Phosphothreonine | Q9UQ35 | Serine/arginine repetitive matrix protein 2 | 1003 | 672 |  |
| P46821 | 0,000898585 | Phosphoserine | P46821 | Microtubule-associated protein 1B;MAP1B heavy chain;MAP1 light chain LC1 | 561 | 673 |  |
| Q9NVD7 | 0,000947639 | Phosphoserine | Q9NVD7 | Alpha-parvin | 8 | 674 |  |
| P46821 | 0,002383709 | Phosphoserine | P46821 | Microtubule-associated protein 1B;MAP1B heavy chain;MAP1 light chain LC1 | 541 | 675 |  |
| Q96JM3 | 0,003035903 | Phosphoserine | Q96JM3 | Chromosome alignment-maintaining phosphoprotein 1 | 476 | 676 |  |
| Q9C0C2 | 0,003410861 | Phosphoserine | Q9C0C2 | 182 kDa tankyrase-1-binding protein | 601 | 677 |  |
| Q9UQ35 | 0,006471857 | Phosphoserine | Q9UQ35 | Serine/arginine repetitive matrix protein 2 | 2702 | 678 |  |
| Q9UQ35 | 0,006471857 | Phosphoserine | Q9UQ35 | Serine/arginine repetitive matrix protein 2 | 2706 | 679 |  |
| P49792 | 0,006536901 | Phosphoserine | P49792 | E3 SUMO-protein ligase RanBP2 | 2900 | 680 |  |
| Q9UQ35 | 0,006602049 | Phosphoserine | Q9UQ35 | Serine/arginine repetitive matrix protein 2 | 377 | 681 |  |
| A0A087WY61 | 0,00687848 | Phosphoserine | A0A087WY61 | Nuclear mitotic apparatus protein 1 | 1741 | 682 |  |
| F5H836 | 0,007268623 | Phosphotyrosine\| by PTK6 | F5H836 | Paxillin | 119 | 683 |  |
| Q12888 | 0,008259878 | Phosphoserine | Q12888 | Tumor suppressor p53-binding protein 1 | 500 | 684 |  |
| Q92619 | 0,008909523 | Phosphoserine | Q92619 | Minor histocompatibility protein HA-1;Minor histocompatibility antigen HA-1 | 23 | 685 |  |
| Q86W56 | 0,00968869 | Phosphoserine | Q86W56 | Poly(ADP-ribose) glycohydrolase | 137 | 686 |  |
| Q14160 | 0,01325455 | Phosphoserine | Q14160 | Protein scribble homolog | 1378 | 687 |  |
| A0A087WVW7 | 0,01412849 | Phosphoserine | A0A087WVW7 | Methyl-CpG-binding protein 2 | 76 | 688 |  |
| C9IZZ0 | 0,01519589 | Phosphoserine | C9IZZ0 | Ras-related protein Rab-7a | 72 | 689 |  |
| Q99549 | 0,01530902 | Phosphoserine | Q99549 | M-phase phosphoprotein 8 | 51 | 690 |  |
| A0A087WWP4 | 0,01658551 | Phosphoserine | A0A087WWP4 | Putative RNA-binding protein 15 | 630 | 691 |  |
| A9Z1X7 | 0,01807064 | Phosphoserine | A9Z1X7 | Serine/arginine repetitive matrix protein 1 | 260 | 692 |  |
| Q96EN8 | 0,01942532 | Phosphoserine | Q96EN8 | Molybdenum cofactor sulfurase | 530 | 693 |  |
| O60264 | 0,02034383 | Phosphoserine | O60264 | SWI/SNF-related matrix-associated actin-dependent regulator of chromatin subfamily A member 5 | 116 | 694 |  |
| H7C446 | 0,02063377 | Phosphoserine | H7C446 | Suppressor of SWI4 1 homolog | 119 | 695 |  |
| F5GX09 | 0,02180898 | Phosphoserine | F5GX09 | Protein FAM76B | 193 | 696 |  |
| E9PG46 | 0,02237211 | Phosphothreonine | E9PG46 | AP2-associated protein kinase 1 | 620 | 697 |  |
| Q9H0D6 | 0,02402811 | Phosphoserine | Q9H0D6 | 5'-3' exoribonuclease 2 | 448 | 698 |  |
| Q6UN15 | 0,02510428 | Phosphothreonine | Q6UN15 | Pre-mRNA 3'-end-processing factor FIP1 | 494 | 699 |  |
| Q12872 | 0,02603516 | Phosphoserine | Q12872 | Splicing factor, suppressor of white-apricot homolog | 909 | 700 |  |
| F8VVL1 | 0,02606729 | Phosphoserine | F8VVL1 | Density-regulated protein | 73 | 701 |  |
| Q9UQ35 | 0,0268212 | Phosphoserine | Q9UQ35 | Serine/arginine repetitive matrix protein 2 | 398 | 702 |  |
| P46821 | 0,02728625 | Phosphoserine | P46821 | Microtubule-associated protein 1B;MAP1B heavy chain;MAP1 light chain LC1 | 1256 | 703 |  |
| Q5JXL7 | 0,02839199 | Phosphothreonine | Q5JXL7 | Septin-7 | 77 | 704 |  |
| Q86TB9 | 0,02885644 | Phosphoserine | Q86TB9 | Protein PAT1 homolog 1 | 179 | 705 |  |
| H0YEV2 | 0,02906461 | Phosphothreonine | H0YEV2 | Src substrate cortactin | 33 | 706 |  |
| A9Z1X7 | 0,02989706 | Phosphoserine | A9Z1X7 | Serine/arginine repetitive matrix protein 1 | 397 | 707 |  |
| A9Z1X7 | 0,02989706 | Phosphothreonine | A9Z1X7 | Serine/arginine repetitive matrix protein 1 | 401 | 708 |  |
| Q9UQ35 | 0,03013703 | Phosphothreonine | Q9UQ35 | Serine/arginine repetitive matrix protein 2 | 1208 | 709 |  |
| V9GYM8 | 0,03058501 | Phosphoserine | V9GYM8 | Rho guanine nucleotide exchange factor 2 | 998 | 710 |  |
| V9GYM8 | 0,03058501 | Phosphoserine | V9GYM8 | Rho guanine nucleotide exchange factor 2 | 1001 | 711 |  |
| P46821 | 0,03136857 | Phosphoserine | P46821 | Microtubule-associated protein 1B;MAP1B heavy chain;MAP1 light chain LC1 | 1265 | 712 |  |
| Q9C0C2 | 0,03417934 | Phosphoserine | Q9C0C2 | 182 kDa tankyrase-1-binding protein | 1621 | 713 |  |
| F8WBF9 | 0,03583781 | Phosphoserine | F8WBF9 | Protein NDRG3 | 236 | 714 |  |
| Q9C0C2 | 0,03585377 | Phosphoserine | Q9C0C2 | 182 kDa tankyrase-1-binding protein | 1138 | 715 |  |
| Q8NE71 | 0,03668223 | Phosphoserine | Q8NE71 | ATP-binding cassette sub-family F member 1 | 228 | 716 |  |
| O00203 | 0,03949861 | Phosphoserine | O00203 | AP-3 complex subunit beta-1 | 276 | 717 |  |
| H7BZ93 | 0,03967352 | Phosphoserine | H7BZ93 | Histone-lysine N-methyltransferase SETD2 | 342 | 718 |  |
| Q9C0C2 | 0,03970525 | Phosphoserine | Q9C0C2 | 182 kDa tankyrase-1-binding protein | 1385 | 719 |  |
| A0A087WWP4 | 0,04015025 | Phosphoserine | A0A087WWP4 | Putative RNA-binding protein 15 | 626 | 720 |  |
| E7EQT4 | 0,04148437 | Phosphoserine | E7EQT4 | Apoptotic chromatin condensation inducer in the nucleus | 450 | 721 |  |
| E9PNR6 | 0,04223026 | Phosphoserine | E9PNR6 | Rho GTPase-activating protein 1 | 51 | 722 |  |
| Q9UPU5 | 0,04354657 | Phosphoserine | Q9UPU5 | Ubiquitin carboxyl-terminal hydrolase 24 | 2047 | 723 |  |
| Q09666 | 0,04532096 | Phosphothreonine | Q09666 | Neuroblast differentiation-associated protein AHNAK | 101 | 724 |  |
| H0Y7N6 | 0,04582745 | Phosphoserine | H0Y7N6 | Actin-binding LIM protein 1 | 96 | 725 |  |
| O94804 | 0,04695074 | Phosphoserine | O94804 | Serine/threonine-protein kinase 10 | 438 | 726 |  |
| Q9C0C2 | 0,04704566 | Phosphoserine | Q9C0C2 | 182 kDa tankyrase-1-binding protein | 691 | 727 |  |
| E9PDE4 | 0,04971565 | Phosphoserine | E9PDE4 | Calpastatin | 171 | 728 |  |
| H0Y911 | 0,05055198 | Phosphoserine | H0Y911 | Transforming acidic coiled-coil-containing protein 2 | 247 | 729 |  |
| E5RIU6 | 0,05146669 | Phosphotyrosine\| by PKMYT1\| WEE1\| WEE2 and PKC/PRKCD;Phosphotyrosine\| by WEE1 | E5RIU6 | Cyclin-dependent kinase 1;Cyclin-dependent kinase 2;Cyclin-dependent kinase 3 | 15 | 730 |  |
| Q9UDY2 | 0,05190808 | Phosphoserine | Q9UDY2 | Tight junction protein ZO-2 | 1159 | 731 |  |
| H0YEV2 | 0,05348325 | Phosphoserine | H0YEV2 | Src substrate cortactin | 52 | 732 |  |
| Q9UQ35 | 0,05436458 | Phosphothreonine | Q9UQ35 | Serine/arginine repetitive matrix protein 2 | 983 | 733 |  |
| J3KQ96 | 0,05603138 | Phosphoserine | J3KQ96 | Treacle protein | 906 | 734 |  |
| K7EL60 | 0,05730382 | Phosphothreonine | K7EL60 | Eukaryotic translation initiation factor 3 subunit G | 41 | 735 |  |
| P80723 | 0,05940626 | Phosphothreonine | P80723 | Brain acid soluble protein 1 | 196 | 736 |  |
| Q8N573 | 0,05945326 | Phosphoserine | Q8N573 | Oxidation resistance protein 1 | 202 | 737 |  |
| H0YL55 | 0,0609263 | Phosphoserine | H0YL55 | SAFB-like transcription modulator | 213 | 738 |  |
| K7EL60 | 0,06111421 | Phosphoserine | K7EL60 | Eukaryotic translation initiation factor 3 subunit G | 42 | 739 |  |
| Q8N3X1 | 0,06116126 | Phosphoserine | Q8N3X1 | Formin-binding protein 4 | 116 | 740 |  |
| Q8NDI1 | 0,06136481 | Phosphoserine | Q8NDI1 | EH domain-binding protein 1 | 1058 | 741 |  |
| Q5T200 | 0,06271069 | Phosphoserine | Q5T200 | Zinc finger CCCH domain-containing protein 13 | 242 | 742 |  |
| E9PQN2 | 0,06438354 | Phosphotyrosine | E9PQN2 | Bcl-2-associated transcription factor 1 | 282 | 743 |  |
| P54725 | 0,06452417 | Phosphoserine | P54725 | UV excision repair protein RAD23 homolog A | 123 | 744 |  |
| F8W6I7 | 0,06621045 | Phosphoserine | F8W6I7 | Heterogeneous nuclear ribonucleoprotein A1;Heterogeneous nuclear ribonucleoprotein A1, N-terminally processed | 4 | 745 |  |
| F8VQE1 | 0,06814419 | Phosphoserine | F8VQE1 | LIM domain and actin-binding protein 1 | 448 | 746 |  |
| M0QZR4 | 0,06840909 | Phosphoserine | M0QZR4 | Rho guanine nucleotide exchange factor 1 | 919 | 747 |  |
| Q8TEA8 | 0,07065102 | Phosphoserine | Q8TEA8 | D-tyrosyl-tRNA(Tyr) deacylase 1 | 197 | 748 |  |
| E7EQT4 | 0,07321561 | Phosphoserine | E7EQT4 | Apoptotic chromatin condensation inducer in the nucleus | 176 | 749 |  |
| E9PMS6 | 0,07676747 | Phosphoserine | E9PMS6 | LIM domain only protein 7 | 1102 | 750 |  |
| F8W6I7 | 0,07768121 | Phosphoserine | F8W6I7 | Heterogeneous nuclear ribonucleoprotein A1;Heterogeneous nuclear ribonucleoprotein A1, N-terminally processed | 6 | 751 |  |
| A0A0A0MR52 | 0,07791336 | Phosphoserine | A0A0A0MR52 | Eukaryotic translation initiation factor 4 gamma 1 | 1010 | 752 |  |
| E9PS47 | 0,07901195 | Phosphoserine | E9PS47 | Phosphatidylserine synthase 2 | 16 | 753 |  |
| E7EVA0 | 0,08069687 | Phosphoserine | E7EVA0 | Microtubule-associated protein;Microtubule-associated protein 4 | 297 | 754 |  |
| F8WD88 | 0,08109849 | Phosphoserine | F8WD88 | Histone H3-like centromeric protein A | 17 | 755 |  |
| F8WD88 | 0,08109849 | Phosphoserine | F8WD88 | Histone H3-like centromeric protein A | 19 | 756 |  |
| H3BTK3 | 0,08150008 | Phosphoserine | H3BTK3 | Calcium-regulated heat stable protein 1 | 30 | 757 |  |
| H3BTK3 | 0,08150008 | Phosphoserine | H3BTK3 | Calcium-regulated heat stable protein 1 | 32 | 758 |  |
| H3BTK3 | 0,08150008 | Phosphoserine | H3BTK3 | Calcium-regulated heat stable protein 1 | 41 | 759 |  |
| J3QRU8 | 0,08156183 | Phosphoserine | J3QRU8 | ARF GTPase-activating protein GIT1 | 362 | 760 |  |
| V9GYM8 | 0,08259586 | Phosphoserine | V9GYM8 | Rho guanine nucleotide exchange factor 2 | 1005 | 761 |  |
| H3BUF6 | 0,08418418 | Phosphoserine | H3BUF6 | Ataxin-2-like protein | 111 | 762 |  |
| E9PP50 | 0,08553979 | Phosphoserine\| by NRK | E9PP50 | Cofilin-1 | 3 | 763 |  |
| Q6UN15 | 0,08557054 | Phosphoserine | Q6UN15 | Pre-mRNA 3'-end-processing factor FIP1 | 492 | 764 |  |
| C9J0I9 | 0,08723251 | Phosphoserine | C9J0I9 | Nuclear-interacting partner of ALK | 301 | 765 |  |
| Q5HY54 | 0,08893853 | Phosphoserine | Q5HY54 | Filamin-A | 2112 | 766 |  |
| Q9Y463 | 0,09018227 | Phosphotyrosine\| by autocatalysis | Q9Y463 | Dual specificity tyrosine-phosphorylation-regulated kinase 1B;Dual specificity tyrosine-phosphorylation-regulated kinase 1A | 273 | 767 |  |
| Q09666 | 0,0911489 | Phosphothreonine | Q09666 | Neuroblast differentiation-associated protein AHNAK | 5824 | 768 |  |
| Q9BU76 | 0,09213012 | Phosphoserine | Q9BU76 | Multiple myeloma tumor-associated protein 2 | 220 | 769 |  |
| Q8N7R7 | 0,0944427 | Phosphoserine | Q8N7R7 | Cyclin-Y-like protein 1 | 344 | 770 |  |
| Q9C0B5 | 0,09724033 | Phosphoserine | Q9C0B5 | Palmitoyltransferase ZDHHC5;Palmitoyltransferase | 380 | 771 |  |
| Q15149 | 0,0989041 | Phosphoserine | Q15149 | Plectin | 4613 | 772 |  |
| Q09666 | 0,09913296 | Phosphoserine | Q09666 | Neuroblast differentiation-associated protein AHNAK | 5752 | 773 |  |
| Q96GM8 | 0,09931596 | Phosphoserine | Q96GM8 | Target of EGR1 protein 1 | 5 | 774 |  |
| C9JY26 | 0,1003069 | Phosphoserine | C9JY26 | Ras-related protein Rab-34 | 244 | 775 |  |
| Q9P0V3 | 0,1004898 | Phosphoserine | Q9P0V3 | SH3 domain-binding protein 4 | 246 | 776 |  |
| H0YNE5 | 0,1006117 | Phosphoserine | H0YNE5 | Regulator of microtubule dynamics protein 3 | 46 | 777 |  |
| Q92615 | 0,1059501 | Phosphoserine | Q92615 | La-related protein 4B | 498 | 778 |  |
| A0A087WWC8 | 0,1060564 | Phosphoserine | A0A087WWC8 | Myelin expression factor 2 | 17 | 779 |  |
| P11831 | 0,1081951 | Phosphoserine | P11831 | Serum response factor | 224 | 780 |  |
| E9PJ24 | 0,1083164 | Phosphothreonine | E9PJ24 | PHD and RING finger domain-containing protein 1 | 913 | 781 |  |
| Q6IQ22 | 0,1085892 | Phosphoserine | Q6IQ22 | Ras-related protein Rab-12 | 21 | 782 |  |
| E7EVA0 | 0,1109663 | Phosphothreonine | E7EVA0 | Microtubule-associated protein;Microtubule-associated protein 4 | 538 | 783 |  |
| Q15424 | 0,1113595 | Phosphoserine | Q15424 | Scaffold attachment factor B1 | 604 | 784 |  |
| Q96S94 | 0,1120248 | Phosphoserine | Q96S94 | Cyclin-L2 | 369 | 785 |  |
| P51116 | 0,1124632 | Phosphoserine | P51116 | Fragile X mental retardation syndrome-related protein 2 | 601 | 786 |  |
| Q9Y613 | 0,1135509 | Phosphoserine | Q9Y613 | FH1/FH2 domain-containing protein 1 | 523 | 787 |  |
| P51812 | 0,1165831 | Phosphoserine | P51812 | Ribosomal protein S6 kinase alpha-3 | 415 | 788 |  |
| Q9NQC3 | 0,1168693 | Phosphoserine | Q9NQC3 | Reticulon-4 | 15 | 789 |  |
| E7ERS3 | 0,1186311 | Phosphoserine | E7ERS3 | Zinc finger CCCH domain-containing protein 18 | 558 | 790 |  |
| A0A087WZC7 | 0,1190373 | Phosphoserine | A0A087WZC7 | KRR1 small subunit processome component homolog | 3 | 791 |  |
| M0R0F9 | 0,1192028 | Phosphoserine | M0R0F9 | Cdc42-interacting protein 4 | 191 | 792 |  |
| V9GY71 | 0,122103 | Phosphoserine | V9GY71 | 1-phosphatidylinositol 4,5-bisphosphate phosphodiesterase gamma-1 | 66 | 793 |  |
| P28482 | 0,1225983 | Phosphothreonine\| by MAP2K1 and MAP2K2 | P28482 | Mitogen-activated protein kinase 1 | 185 | 794 |  |
| P28482 | 0,1225983 | Phosphotyrosine\| by MAP2K1 and MAP2K2 | P28482 | Mitogen-activated protein kinase 1 | 187 | 795 |  |
| E9PG46 | 0,1231834 | Phosphoserine | E9PG46 | AP2-associated protein kinase 1 | 637 | 796 |  |
| H0YEV2 | 0,1242931 | Phosphothreonine | H0YEV2 | Src substrate cortactin | 35 | 797 |  |
| Q9UDY2 | 0,1248325 | Phosphoserine | Q9UDY2 | Tight junction protein ZO-2 | 978 | 798 |  |
| Q9UDY2 | 0,1248325 | Phosphoserine | Q9UDY2 | Tight junction protein ZO-2 | 986 | 799 |  |
| J3QRS3 | 0,1250124 | Phosphoserine\| by CDC42BP\| CIT\| MLCK\| PAK1\| ROCK1\| ROCK2\| DAPK1\| DAPK2 and ZIPK/DAPK3;Phosphoserine\| by MLCK;Phosphoserine\| by MLCK and ZIPK/DAPK3 | J3QRS3 | Myosin regulatory light chain 12A;Myosin regulatory light chain 12B;Myosin regulatory light polypeptide 9 | 25 | 800 |  |
| J3QRS3 | 0,1250124 | Phosphothreonine\| by MLCK;Phosphothreonine\| by MLCK and ZIPK/DAPK3;Phosphothreonine\| by MLCK\| CIT and ROCK2 | J3QRS3 | Myosin regulatory light chain 12A;Myosin regulatory light chain 12B;Myosin regulatory light polypeptide 9 | 24 | 801 |  |
| Q9C0C2 | 0,1251172 | Phosphoserine | Q9C0C2 | 182 kDa tankyrase-1-binding protein | 1666 | 802 |  |
| P36871 | 0,1255067 | Phosphoserine | P36871 | Phosphoglucomutase-1 | 117 | 803 |  |
| Q9UQ35 | 0,1255217 | Phosphoserine | Q9UQ35 | Serine/arginine repetitive matrix protein 2 | 2394 | 804 |  |
| Q15149 | 0,1269437 | Phosphoserine | Q15149 | Plectin | 4386 | 805 |  |
| M0R0F9 | 0,1277215 | Phosphoserine | M0R0F9 | Cdc42-interacting protein 4 | 188 | 806 |  |
| P48634 | 0,1280056 | Phosphoserine | P48634 | Protein PRRC2A | 342 | 807 |  |
| P48634 | 0,1280056 | Phosphoserine | P48634 | Protein PRRC2A | 350 | 808 |  |
| F8W7U8 | 0,130739 | Phosphoserine | F8W7U8 | Double-strand break repair protein MRE11A | 648 | 809 |  |
| Q7Z434 | 0,1317235 | Phosphoserine | Q7Z434 | Mitochondrial antiviral-signaling protein | 222 | 810 |  |
| Q14573 | 0,1318726 | Phosphoserine | Q14573 | Inositol 1,4,5-trisphosphate receptor type 3 | 2670 | 811 |  |
| H3BQZ7 | 0,1325435 | Phosphoserine | H3BQZ7 | Heterogeneous nuclear ribonucleoprotein U-like protein 2 | 161 | 812 |  |
| Q9UKX7 | 0,1326776 | Phosphoserine | Q9UKX7 | Nuclear pore complex protein Nup50 | 221 | 813 |  |
| Q9NTI5 | 0,1344052 | Phosphoserine | Q9NTI5 | Sister chromatid cohesion protein PDS5 homolog B | 1383 | 814 |  |
| Q09666 | 0,1349261 | Phosphoserine | Q09666 | Neuroblast differentiation-associated protein AHNAK | 177 | 815 |  |
| Q9UQ35 | 0,1355509 | Phosphoserine | Q9UQ35 | Serine/arginine repetitive matrix protein 2 | 1124 | 816 |  |
| Q09666 | 0,1363389 | Phosphoserine | Q09666 | Neuroblast differentiation-associated protein AHNAK | 41 | 817 |  |
| A9Z1X7 | 0,1363538 | Phosphoserine | A9Z1X7 | Serine/arginine repetitive matrix protein 1 | 722 | 818 |  |
| F8VQE1 | 0,1382997 | Phosphoserine | F8VQE1 | LIM domain and actin-binding protein 1 | 214 | 819 |  |
| Q9NQQ7 | 0,1394126 | Phosphoserine | Q9NQQ7 | Solute carrier family 35 member C2 | 335 | 820 |  |
| A9Z1X7 | 0,1397241 | Phosphoserine | A9Z1X7 | Serine/arginine repetitive matrix protein 1 | 445 | 821 |  |
| A9Z1X7 | 0,1397241 | Phosphoserine | A9Z1X7 | Serine/arginine repetitive matrix protein 1 | 447 | 822 |  |
| P60981 | 0,1403616 | Phosphoserine | P60981 | Destrin | 3 | 823 |  |
| P02545 | 0,1420653 | Phosphoserine | P02545 | Prelamin-A/C;Lamin-A/C | 12 | 824 |  |
| Q9UQ35 | 0,1431457 | Phosphoserine | Q9UQ35 | Serine/arginine repetitive matrix protein 2 | 994 | 825 |  |
| Q9UQ35 | 0,1432788 | Phosphoserine | Q9UQ35 | Serine/arginine repetitive matrix protein 2 | 876 | 826 |  |
| Q5T200 | 0,1440035 | Phosphoserine | Q5T200 | Zinc finger CCCH domain-containing protein 13 | 877 | 827 |  |
| Q9UQ35 | 0,1440035 | Phosphothreonine | Q9UQ35 | Serine/arginine repetitive matrix protein 2 | 1003 | 828 |  |
| Q9UQ35 | 0,1448313 | Phosphoserine | Q9UQ35 | Serine/arginine repetitive matrix protein 2 | 2581 | 829 |  |
| E9PJF4 | 0,1457324 | Phosphoserine | E9PJF4 | Methylosome subunit pICln | 102 | 830 |  |
| F8WCJ4 | 0,1474445 | Phosphoserine | F8WCJ4 | Acetyl-coenzyme A synthetase, cytoplasmic | 30 | 831 |  |
| O60832 | 0,1488157 | Phosphoserine | O60832 | H/ACA ribonucleoprotein complex subunit 4 | 513 | 832 |  |
| Q5TBM3 | 0,1503916 | Phosphoserine | Q5TBM3 | Heat shock protein 105 kDa | 73 | 833 |  |
| H3BTK3 | 0,1518188 | Phosphoserine | H3BTK3 | Calcium-regulated heat stable protein 1 | 30 | 834 |  |
| O60343 | 0,1518335 | Phosphoserine | O60343 | TBC1 domain family member 4 | 341 | 835 |  |
| Q9H1B7 | 0,1521864 | Phosphoserine | Q9H1B7 | Interferon regulatory factor 2-binding protein-like | 657 | 836 |  |
| Q9H1B7 | 0,1521864 | Phosphoserine | Q9H1B7 | Interferon regulatory factor 2-binding protein-like | 659 | 837 |  |
| Q3B726 | 0,1532886 | Phosphoserine | Q3B726 | DNA-directed RNA polymerase I subunit RPA43 | 316 | 838 |  |
| H3BNA1 | 0,1538174 | Phosphoserine | H3BNA1 | Ubiquitin carboxyl-terminal hydrolase 10 | 88 | 839 |  |
| A9Z1X7 | 0,1576892 | Phosphoserine | A9Z1X7 | Serine/arginine repetitive matrix protein 1 | 614 | 840 |  |
| A9Z1X7 | 0,1576892 | Phosphoserine | A9Z1X7 | Serine/arginine repetitive matrix protein 1 | 616 | 841 |  |
| Q9UQ35 | 0,1579381 | Phosphoserine | Q9UQ35 | Serine/arginine repetitive matrix protein 2 | 1320 | 842 |  |
| Q9UQ35 | 0,1579381 | Phosphoserine | Q9UQ35 | Serine/arginine repetitive matrix protein 2 | 1329 | 843 |  |
| Q8NE71 | 0,1591823 | Phosphoserine\| by CK2 | Q8NE71 | ATP-binding cassette sub-family F member 1 | 109 | 844 |  |
| E3W994 | 0,1592701 | Phosphoserine | E3W994 | CLIP-associating protein 2 | 370 | 845 |  |
| Q9UQ35 | 0,1602793 | Phosphothreonine | Q9UQ35 | Serine/arginine repetitive matrix protein 2 | 2409 | 846 |  |
| Q7Z6E9 | 0,1603377 | Phosphoserine | Q7Z6E9 | E3 ubiquitin-protein ligase RBBP6 | 1328 | 847 |  |
| Q09666 | 0,1606447 | Phosphothreonine | Q09666 | Neuroblast differentiation-associated protein AHNAK | 4100 | 848 |  |
| H3BUF6 | 0,1627333 | Phosphoserine | H3BUF6 | Ataxin-2-like protein | 690 | 849 |  |
| H0Y2V6 | 0,1629084 | Phosphoserine | H0Y2V6 | Centrosomal protein of 170 kDa | 1076 | 850 |  |
| Q9H1E3 | 0,1630106 | Phosphoserine | Q9H1E3 | Nuclear ubiquitous casein and cyclin-dependent kinase substrate 1 | 181 | 851 |  |
| Q13247 | 0,1633899 | Phosphoserine\| by DYRK1A | Q13247 | Serine/arginine-rich splicing factor 6 | 303 | 852 |  |
| O75396 | 0,1640609 | Phosphoserine | O75396 | Vesicle-trafficking protein SEC22b | 137 | 853 |  |
| H0YA82 | 0,1645857 | Phosphoserine | H0YA82 | La-related protein 7 | 42 | 854 |  |
| A0A087WUT6 | 0,1648335 | Phosphoserine | A0A087WUT6 | Eukaryotic translation initiation factor 5B | 214 | 855 |  |
| M0QXD6 | 0,1672798 | Phosphoserine | M0QXD6 | General transcription factor IIF subunit 1 | 301 | 856 |  |
| Q7Z422 | 0,1678762 | Phosphoserine | Q7Z422 | SUZ domain-containing protein 1 | 107 | 857 |  |
| K7ENL0 | 0,1690682 | Phosphoserine | K7ENL0 | Septin-9 | 70 | 858 |  |
| Q5T8C6 | 0,1694315 | Phosphoserine | Q5T8C6 | Cell division cycle protein 16 homolog | 415 | 859 |  |
| P19338 | 0,1695912 | Phosphothreonine | P19338 | Nucleolin | 69 | 860 |  |
| Q9H0B6 | 0,1726095 | Phosphoserine | Q9H0B6 | Kinesin light chain 2 | 582 | 861 |  |
| F8W6G1 | 0,1727109 | Phosphothreonine | F8W6G1 | Nuclear receptor-binding protein | 441 | 862 |  |
| A0A0C4DG89 | 0,1730877 | Phosphoserine | A0A0C4DG89 | Probable ATP-dependent RNA helicase DDX46 | 804 | 863 |  |
| P08670 | 0,1733776 | Phosphoserine | P08670 | Vimentin | 73 | 864 |  |
| H3BTK3 | 0,1743768 | Phosphoserine | H3BTK3 | Calcium-regulated heat stable protein 1 | 41 | 865 |  |
| O00567 | 0,1749269 | Phosphoserine | O00567 | Nucleolar protein 56 | 570 | 866 |  |
| Q9P2D3 | 0,1757372 | Phosphoserine | Q9P2D3 | HEAT repeat-containing protein 5B | 1737 | 867 |  |
| Q12873 | 0,1760119 | Phosphoserine | Q12873 | Chromodomain-helicase-DNA-binding protein 3 | 713 | 868 |  |
| P49792 | 0,1769227 | Phosphoserine | P49792 | E3 SUMO-protein ligase RanBP2;RanBP2-like and GRIP domain-containing protein 4;RanBP2-like and GRIP domain-containing protein 3;RANBP2-like and GRIP domain-containing protein 5/6;RANBP2-like and GRIP domain-containing protein 8;RANBP2-like and GRIP domain-containing protein 1;RANBP2-like and GRIP domain-containing protein 2 | 788 | 869 |  |
| F8VX12 | 0,1773707 | Phosphoserine | F8VX12 | Sodium-coupled neutral amino acid transporter 1 | 52 | 870 |  |
| Q9C0C2 | 0,1782805 | Phosphoserine | Q9C0C2 | 182 kDa tankyrase-1-binding protein | 1620 | 871 |  |
| Q9C0C2 | 0,1782805 | Phosphoserine | Q9C0C2 | 182 kDa tankyrase-1-binding protein | 1621 | 872 |  |
| H3BTK3 | 0,1785404 | Phosphoserine | H3BTK3 | Calcium-regulated heat stable protein 1 | 30 | 873 |  |
| H3BTK3 | 0,1785404 | Phosphoserine | H3BTK3 | Calcium-regulated heat stable protein 1 | 32 | 874 |  |
| Q05209 | 0,1788147 | Phosphoserine | Q05209 | Tyrosine-protein phosphatase non-receptor type 12 | 673 | 875 |  |
| J3KNP4 | 0,1803149 | Phosphoserine | J3KNP4 | Semaphorin-4B | 830 | 876 |  |
| H3BPE1 | 0,1811798 | Phosphoserine | H3BPE1 | Microtubule-actin cross-linking factor 1, isoforms 1/2/3/5 | 4516 | 877 |  |
| E9PHI6 | 0,182764 | Phosphoserine | E9PHI6 | Cytoplasmic dynein 1 light intermediate chain 1 | 91 | 878 |  |
| E7EVA0 | 0,1839151 | Phosphothreonine | E7EVA0 | Microtubule-associated protein;Microtubule-associated protein 4 | 299 | 879 |  |
| Q9Y3Q8 | 0,1840589 | Phosphoserine | Q9Y3Q8 | TSC22 domain family protein 4 | 279 | 880 |  |
| Q5T757 | 0,1852088 | Phosphoserine | Q5T757 | Serine/arginine-rich splicing factor 11 | 389 | 881 |  |
| A0A087WY71 | 0,1854962 | Phosphothreonine | A0A087WY71 | AP-2 complex subunit mu | 155 | 882 |  |
| Q13469 | 0,1859272 | Phosphoserine | Q13469 | Nuclear factor of activated T-cells, cytoplasmic 2 | 148 | 883 |  |
| Q9H3Z4 | 0,1862143 | Phosphoserine | Q9H3Z4 | DnaJ homolog subfamily C member 5 | 10 | 884 |  |
| E9PHI6 | 0,1867887 | Phosphoserine | E9PHI6 | Cytoplasmic dynein 1 light intermediate chain 1 | 394 | 885 |  |
| Q8TB61 | 0,1870758 | Phosphoserine | Q8TB61 | Adenosine 3'-phospho 5'-phosphosulfate transporter 1 | 427 | 886 |  |
| Q66K74 | 0,1877931 | Phosphoserine | Q66K74 | Microtubule-associated protein 1S;MAP1S heavy chain;MAP1S light chain | 657 | 887 |  |
| Q12888 | 0,1883667 | Phosphoserine | Q12888 | Tumor suppressor p53-binding protein 1 | 1362 | 888 |  |
| P40818 | 0,18851 | Phosphoserine | P40818 | Ubiquitin carboxyl-terminal hydrolase 8 | 718 | 889 |  |
| Q14160 | 0,1890835 | Phosphoserine | Q14160 | Protein scribble homolog | 1220 | 890 |  |
| Q09666 | 0,190659 | Phosphoserine | Q09666 | Neuroblast differentiation-associated protein AHNAK | 93 | 891 |  |
| O00443 | 0,1915176 | Phosphoserine | O00443 | Phosphatidylinositol 4-phosphate 3-kinase C2 domain-containing subunit alpha | 259 | 892 |  |
| E9PI52 | 0,1926617 | Phosphoserine | E9PI52 | Arginine/serine-rich coiled-coil protein 2 | 30 | 893 |  |
| E9PI52 | 0,1926617 | Phosphoserine | E9PI52 | Arginine/serine-rich coiled-coil protein 2 | 32 | 894 |  |
| P06748 | 0,1935191 | Phosphoserine\| by CDK2 | P06748 | Nucleophosmin | 125 | 895 |  |
| Q9NXG2 | 0,1938049 | Phosphoserine | Q9NXG2 | THUMP domain-containing protein 1 | 86 | 896 |  |
| Q9NXG2 | 0,1938049 | Phosphoserine | Q9NXG2 | THUMP domain-containing protein 1 | 88 | 897 |  |
| E9PS34 | 0,199365 | Phosphoserine | E9PS34 | Nucleosome assembly protein 1-like 4 | 12 | 898 |  |
| Q13263 | 0,1995074 | Phosphoserine | Q13263 | Transcription intermediary factor 1-beta | 473 | 899 |  |
| O43379 | 0,2010715 | Phosphoserine | O43379 | WD repeat-containing protein 62 | 1228 | 900 |  |
| Q9H2G2 | 0,2017819 | Phosphoserine | Q9H2G2 | STE20-like serine/threonine-protein kinase | 571 | 901 |  |
| P48634 | 0,2043367 | Phosphoserine | P48634 | Protein PRRC2A | 761 | 902 |  |
| G5E972 | 0,2043367 | Phosphothreonine | G5E972 | Lamina-associated polypeptide 2, isoforms beta/gamma;Thymopoietin;Thymopentin;Lamina-associated polypeptide 2, isoform alpha;Thymopoietin;Thymopentin | 74 | 903 |  |
| Q13523 | 0,2049037 | Phosphoserine | Q13523 | Serine/threonine-protein kinase PRP4 homolog | 277 | 904 |  |
| Q6NYC8 | 0,2051872 | Phosphoserine | Q6NYC8 | Phostensin | 368 | 905 |  |
| P46821 | 0,2056123 | Phosphoserine | P46821 | Microtubule-associated protein 1B;MAP1B heavy chain;MAP1 light chain LC1 | 1427 | 906 |  |
| E9PL71 | 0,2061789 | Phosphoserine\| by CK2 | E9PL71 | Elongation factor 1-delta | 138 | 907 |  |
| P46821 | 0,2070283 | Phosphothreonine | P46821 | Microtubule-associated protein 1B;MAP1B heavy chain;MAP1 light chain LC1 | 1788 | 908 |  |
| P46821 | 0,2084431 | Phosphoserine | P46821 | Microtubule-associated protein 1B;MAP1B heavy chain;MAP1 light chain LC1 | 1312 | 909 |  |
| Q8NC51 | 0,2087257 | Phosphoserine | Q8NC51 | Plasminogen activator inhibitor 1 RNA-binding protein | 234 | 910 |  |
| Q92574 | 0,2087257 | Phosphoserine | Q92574 | Hamartin | 505 | 911 |  |
| A9Z1X7 | 0,2108448 | Phosphoserine | A9Z1X7 | Serine/arginine repetitive matrix protein 1 | 705 | 912 |  |
| Q9UDY2 | 0,2115505 | Phosphoserine | Q9UDY2 | Tight junction protein ZO-2 | 266 | 913 |  |
| A0A0A0MR52 | 0,2119736 | Phosphothreonine | A0A0A0MR52 | Eukaryotic translation initiation factor 4 gamma 1 | 1012 | 914 |  |
| Q01105 | 0,2138061 | Phosphoserine | Q01105 | Protein SET | 7 | 915 |  |
| C9J0I9 | 0,2149327 | Phosphoserine | C9J0I9 | Nuclear-interacting partner of ALK | 278 | 916 |  |
| P08621 | 0,2152143 | Phosphoserine | P08621 | U1 small nuclear ribonucleoprotein 70 kDa | 226 | 917 |  |
| Q13242 | 0,2164804 | Phosphoserine | Q13242 | Serine/arginine-rich splicing factor 9 | 211 | 918 |  |
| P46821 | 0,2184476 | Phosphoserine | P46821 | Microtubule-associated protein 1B;MAP1B heavy chain;MAP1 light chain LC1 | 1252 | 919 |  |
| O75592 | 0,2191495 | Phosphoserine | O75592 | E3 ubiquitin-protein ligase MYCBP2 | 3467 | 920 |  |
| F8WBS8 | 0,2192899 | Phosphoserine | F8WBS8 | 26S proteasome non-ATPase regulatory subunit 2 | 16 | 921 |  |
| I3L436 | 0,2205524 | Phosphoserine\| by MAPK8 and MTOR | I3L436 | Regulatory-associated protein of mTOR | 23 | 922 |  |
| I3L436 | 0,2205524 | Phosphoserine\| by MTOR | I3L436 | Regulatory-associated protein of mTOR | 19 | 923 |  |
| C9JWF0 | 0,2206927 | Phosphoserine | C9JWF0 | Structural maintenance of chromosomes protein 4 | 41 | 924 |  |
| F8W020 | 0,2208327 | Phosphothreonine | F8W020 | Nucleosome assembly protein 1-like 1 | 62 | 925 |  |
| E9PL71 | 0,2212533 | Phosphoserine | E9PL71 | Elongation factor 1-delta | 109 | 926 |  |
| P48681 | 0,2216738 | Phosphoserine | P48681 | Nestin | 680 | 927 |  |
| Q96PK6 | 0,2226542 | Phosphothreonine | Q96PK6 | RNA-binding protein 14 | 206 | 928 |  |
| Q9Y2W1 | 0,2260105 | Phosphoserine | Q9Y2W1 | Thyroid hormone receptor-associated protein 3 | 672 | 929 |  |
| Q09666 | 0,2261502 | Phosphoserine | Q09666 | Neuroblast differentiation-associated protein AHNAK | 5077 | 930 |  |
| Q8WX93 | 0,2265692 | Phosphoserine | Q8WX93 | Palladin | 893 | 931 |  |
| P46821 | 0,2267089 | Phosphoserine | P46821 | Microtubule-associated protein 1B;MAP1B heavy chain;MAP1 light chain LC1 | 1915 | 932 |  |
| Q05655 | 0,2274067 | Phosphoserine\| by autocatalysis | Q05655 | Protein kinase C delta type;Protein kinase C delta type regulatory subunit;Protein kinase C delta type catalytic subunit | 304 | 933 |  |
| O95466 | 0,2289409 | Phosphoserine | O95466 | Formin-like protein 1 | 184 | 934 |  |
| Q99961 | 0,2290803 | Phosphoserine | Q99961 | Endophilin-A2 | 288 | 935 |  |
| Q00839 | 0,2297772 | Phosphoserine | Q00839 | Heterogeneous nuclear ribonucleoprotein U | 4 | 936 |  |
| A0A087WUZ3 | 0,2297772 | Phosphoserine | A0A087WUZ3 | Spectrin beta chain, non-erythrocytic 1 | 2166 | 937 |  |
| P08670 | 0,2300557 | Phosphoserine | P08670 | Vimentin | 55 | 938 |  |
| P10412 | 0,2304736 | Phosphoserine | P10412 | Histone H1.4;Histone H1.3;Histone H1.2 | 36 | 939 |  |
| Q9UQ35 | 0,232978 | Phosphoserine | Q9UQ35 | Serine/arginine repetitive matrix protein 2 | 1320 | 940 |  |
| P35580 | 0,233256 | Phosphoserine | P35580 | Myosin-10 | 1956 | 941 |  |
| G3V2R1 | 0,233395 | Phosphoserine | G3V2R1 | Protein Smaug homolog 1 | 11 | 942 |  |
| Q01804 | 0,2342286 | Phosphoserine | Q01804 | OTU domain-containing protein 4 | 1006 | 943 |  |
| Q16643 | 0,2342286 | Phosphoserine | Q16643 | Drebrin | 339 | 944 |  |
| Q29RF7 | 0,2368651 | Phosphoserine | Q29RF7 | Sister chromatid cohesion protein PDS5 homolog A | 1305 | 945 |  |
| E9PCT5 | 0,2375583 | Phosphoserine | E9PCT5 | Caveolin;Caveolin-1 | 26 | 946 |  |
| Q00839 | 0,239497 | Phosphoserine | Q00839 | Heterogeneous nuclear ribonucleoprotein U | 271 | 947 |  |
| Q9UQ35 | 0,2397738 | Phosphoserine | Q9UQ35 | Serine/arginine repetitive matrix protein 2 | 992 | 948 |  |
| O14617 | 0,239912 | Phosphoserine | O14617 | AP-3 complex subunit delta-1 | 658 | 949 |  |
| A0A087WWP4 | 0,2403271 | Phosphothreonine | A0A087WWP4 | Putative RNA-binding protein 15 | 524 | 950 |  |
| Q03164 | 0,2417096 | Phosphoserine | Q03164 | Histone-lysine N-methyltransferase 2A;MLL cleavage product N320;MLL cleavage product C180 | 2098 | 951 |  |
| A0A087WWP4 | 0,2419858 | Phosphoserine | A0A087WWP4 | Putative RNA-binding protein 15 | 656 | 952 |  |
| Q9UQ35 | 0,2419858 | Phosphoserine | Q9UQ35 | Serine/arginine repetitive matrix protein 2 | 2688 | 953 |  |
| Q9UJX2 | 0,2435047 | Phosphoserine | Q9UJX2 | Cell division cycle protein 23 homolog | 588 | 954 |  |
| A9Z1X7 | 0,2451598 | Phosphothreonine | A9Z1X7 | Serine/arginine repetitive matrix protein 1 | 220 | 955 |  |
| E9PJ38 | 0,2459866 | Phosphoserine | E9PJ38 | Receptor-binding cancer antigen expressed on SiSo cells | 36 | 956 |  |
| Q9H6S3 | 0,2473637 | Phosphoserine | Q9H6S3 | Epidermal growth factor receptor kinase substrate 8-like protein 2 | 570 | 957 |  |
| Q9UQ35 | 0,247914 | Phosphoserine | Q9UQ35 | Serine/arginine repetitive matrix protein 2 | 2398 | 958 |  |
| E9PQN2 | 0,2514868 | Phosphoserine | E9PQN2 | Bcl-2-associated transcription factor 1 | 266 | 959 |  |
| E7EVA0 | 0,2520356 | Phosphoserine | E7EVA0 | Microtubule-associated protein;Microtubule-associated protein 4 | 14 | 960 |  |
| P19338 | 0,2528585 | Phosphoserine | P19338 | Nucleolin | 67 | 961 |  |
| O15439 | 0,2558718 | Phosphothreonine | O15439 | Multidrug resistance-associated protein 4 | 646 | 962 |  |
| Q16643 | 0,2562823 | Phosphoserine | Q16643 | Drebrin | 337 | 963 |  |
| Q14573 | 0,2564191 | Phosphoserine | Q14573 | Inositol 1,4,5-trisphosphate receptor type 3 | 1832 | 964 |  |
| E9PHI6 | 0,2571028 | Phosphoserine | E9PHI6 | Cytoplasmic dynein 1 light intermediate chain 1 | 400 | 965 |  |
| H7C1L9 | 0,2577861 | Phosphoserine | H7C1L9 | E3 ubiquitin-protein ligase TRIP12 | 153 | 966 |  |
| H3BPZ1 | 0,2583327 | Phosphoserine | H3BPZ1 | Very-long-chain (3R)-3-hydroxyacyl-CoA dehydratase 3 | 89 | 967 |  |
| O00499 | 0,2586057 | Phosphoserine | O00499 | Myc box-dependent-interacting protein 1 | 298 | 968 |  |
| F5H6E2 | 0,2602437 | Phosphoserine | F5H6E2 | Unconventional myosin-Ic | 384 | 969 |  |
| Q9UQ35 | 0,2606528 | Phosphoserine | Q9UQ35 | Serine/arginine repetitive matrix protein 2 | 952 | 970 |  |
| Q9UQ35 | 0,2606528 | Phosphoserine | Q9UQ35 | Serine/arginine repetitive matrix protein 2 | 954 | 971 |  |
| F8W930 | 0,2646022 | Phosphoserine | F8W930 | Insulin-like growth factor 2 mRNA-binding protein 2 | 168 | 972 |  |
| M0R0F9 | 0,2662334 | Phosphoserine | M0R0F9 | Cdc42-interacting protein 4 | 188 | 973 |  |
| Q99613 | 0,2677269 | Phosphoserine | Q99613 | Eukaryotic translation initiation factor 3 subunit C;Eukaryotic translation initiation factor 3 subunit C-like protein | 39 | 974 |  |
| Q6UN15 | 0,2682697 | Phosphoserine | Q6UN15 | Pre-mRNA 3'-end-processing factor FIP1 | 492 | 975 |  |
| P46821 | 0,2697611 | Phosphoserine | P46821 | Microtubule-associated protein 1B;MAP1B heavy chain;MAP1 light chain LC1 | 1797 | 976 |  |
| Q02040 | 0,2727394 | Phosphoserine | Q02040 | A-kinase anchor protein 17A | 537 | 977 |  |
| Q09666 | 0,2730098 | Phosphoserine | Q09666 | Neuroblast differentiation-associated protein AHNAK | 5790 | 978 |  |
| H0YBM4 | 0,2742261 | Phosphoserine | H0YBM4 | Arf-GAP with SH3 domain, ANK repeat and PH domain-containing protein 1 | 131 | 979 |  |
| P07814 | 0,2746314 | Phosphoserine\| by CDK5 | P07814 | Bifunctional glutamate/proline--tRNA ligase;Glutamate--tRNA ligase;Proline--tRNA ligase | 886 | 980 |  |
| B7ZKW8 | 0,2747665 | Phosphoserine\| by MAPK8\| in vitro | B7ZKW8 | CapZ-interacting protein | 186 | 981 |  |
| Q5JSH3 | 0,2750366 | Phosphoserine | Q5JSH3 | WD repeat-containing protein 44 | 50 | 982 |  |
| Q15424 | 0,2804272 | Phosphoserine | Q15424 | Scaffold attachment factor B1 | 604 | 983 |  |
| P27824 | 0,2837864 | Phosphoserine\| by MAPK3 | P27824 | Calnexin | 564 | 984 |  |
| Q9H1E3 | 0,2861999 | Phosphoserine | Q9H1E3 | Nuclear ubiquitous casein and cyclin-dependent kinase substrate 1 | 19 | 985 |  |
| Q9Y2W1 | 0,2863339 | Phosphoserine | Q9Y2W1 | Thyroid hormone receptor-associated protein 3 | 248 | 986 |  |
| Q9Y2W1 | 0,2863339 | Phosphoserine | Q9Y2W1 | Thyroid hormone receptor-associated protein 3 | 253 | 987 |  |
| E7EVA0 | 0,2874053 | Phosphoserine | E7EVA0 | Microtubule-associated protein;Microtubule-associated protein 4 | 1841 | 988 |  |
| A0A087WZH7 | 0,287673 | Phosphoserine | A0A087WZH7 | Myristoylated alanine-rich C-kinase substrate | 46 | 989 |  |
| E7EQT4 | 0,2899465 | Phosphoserine | E7EQT4 | Apoptotic chromatin condensation inducer in the nucleus | 689 | 990 |  |
| Q5VSL9 | 0,2900801 | Phosphoserine | Q5VSL9 | Striatin-interacting protein 1 | 335 | 991 |  |
| H7C2Y0 | 0,2903474 | Phosphoserine | H7C2Y0 | Septin-2 | 73 | 992 |  |
| H7BXI1 | 0,290748 | Phosphoserine | H7BXI1 | Extended synaptotagmin-2 | 724 | 993 |  |
| P08238 | 0,2908816 | Phosphoserine | P08238 | Heat shock protein HSP 90-beta;Putative heat shock protein HSP 90-beta 2 | 255 | 994 |  |
| J3KQ96 | 0,2911487 | Phosphoserine | J3KQ96 | Treacle protein | 381 | 995 |  |
| Q5JSH3 | 0,2915493 | Phosphoserine | Q5JSH3 | WD repeat-containing protein 44 | 403 | 996 |  |
| G5E972 | 0,2918161 | Phosphoserine | G5E972 | Lamina-associated polypeptide 2, isoforms beta/gamma;Thymopoietin;Thymopentin | 184 | 997 |  |
| H3BQN4 | 0,2922165 | Phosphoserine | H3BQN4 | Fructose-bisphosphate aldolase A | 46 | 998 |  |
| Q8NE71 | 0,2924834 | Phosphothreonine | Q8NE71 | ATP-binding cassette sub-family F member 1 | 108 | 999 |  |
| Q9HAZ1 | 0,2928834 | Phosphoserine | Q9HAZ1 | Dual specificity protein kinase CLK4 | 138 | 1000 |  |
| E9PQN2 | 0,2936833 | Phosphoserine | E9PQN2 | Bcl-2-associated transcription factor 1 | 510 | 1001 |  |
| H3BP35 | 0,295415 | Phosphoserine | H3BP35 | Diphosphomevalonate decarboxylase | 96 | 1002 |  |
| H0YBX5 | 0,2976764 | Phosphoserine | H0YBX5 | Cyclin-dependent kinase 16;Cyclin-dependent kinase 17 | 79 | 1003 |  |
| Q9BUH6 | 0,2978094 | Phosphoserine | Q9BUH6 | Uncharacterized protein C9orf142 | 152 | 1004 |  |
| P12270 | 0,2988722 | Phosphoserine | P12270 | Nucleoprotein TPR | 2155 | 1005 |  |
| Q8ND76 | 0,2999343 | Phosphoserine | Q8ND76 | Cyclin-Y | 326 | 1006 |  |
| C9JA08 | 0,3017911 | Phosphothreonine | C9JA08 | 60S ribosomal export protein NMD3 | 470 | 1007 |  |
| C9J2I0 | 0,3037778 | Phosphoserine | C9J2I0 | Arf-GAP domain and FG repeat-containing protein 1 | 103 | 1008 |  |
| Q13523 | 0,3037778 | Phosphotyrosine | Q13523 | Serine/threonine-protein kinase PRP4 homolog | 849 | 1009 |  |
| P08670 | 0,3051007 | Phosphoserine\| by CDK5 and CDK1 | P08670 | Vimentin | 56 | 1010 |  |
| Q6UN15 | 0,3068188 | Phosphoserine | Q6UN15 | Pre-mRNA 3'-end-processing factor FIP1 | 304 | 1011 |  |
| E9PS34 | 0,307479 | Phosphoserine | E9PS34 | Nucleosome assembly protein 1-like 4 | 125 | 1012 |  |
| M0R300 | 0,3099852 | Phosphoserine | M0R300 | Unconventional myosin-IXb | 1290 | 1013 |  |
| Q06210 | 0,3101169 | Phosphoserine | Q06210 | Glutamine--fructose-6-phosphate aminotransferase [isomerizing] 1 | 261 | 1014 |  |
| Q9UQ35 | 0,3107756 | Phosphoserine | Q9UQ35 | Serine/arginine repetitive matrix protein 2 | 1083 | 1015 |  |
| Q14204 | 0,3139334 | Phosphoserine | Q14204 | Cytoplasmic dynein 1 heavy chain 1 | 4368 | 1016 |  |
| J3QQX0 | 0,3147218 | Phosphoserine | J3QQX0 | COP9 signalosome complex subunit 1 | 62 | 1017 |  |
| O14974 | 0,3152472 | Phosphoserine\| by NUAK1 | O14974 | Protein phosphatase 1 regulatory subunit 12A | 910 | 1018 |  |
| H3BPL0 | 0,3153784 | Phosphoserine | H3BPL0 | Battenin | 12 | 1019 |  |
| Q8IXT5 | 0,3164284 | Phosphoserine | Q8IXT5 | RNA-binding protein 12B | 638 | 1020 |  |
| J3QR07 | 0,3172155 | Phosphoserine | J3QR07 | YTH domain-containing protein 1 | 308 | 1021 |  |
| O75822 | 0,3183951 | Phosphoserine | O75822 | Eukaryotic translation initiation factor 3 subunit J | 13 | 1022 |  |
| Q09666 | 0,3187882 | Phosphoserine | Q09666 | Neuroblast differentiation-associated protein AHNAK | 5793 | 1023 |  |
| H0YE40 | 0,3190501 | Phosphoserine | H0YE40 | CD44 antigen | 46 | 1024 |  |
| P51858 | 0,3193119 | Phosphoserine | P51858 | Hepatoma-derived growth factor | 165 | 1025 |  |
| M0R3F1 | 0,3207516 | Phosphoserine | M0R3F1 | Heterogeneous nuclear ribonucleoprotein U-like protein 1 | 43 | 1026 |  |
| O14974 | 0,3208825 | Phosphothreonine | O14974 | Protein phosphatase 1 regulatory subunit 12A | 443 | 1027 |  |
| E9PQN2 | 0,3219285 | Phosphoserine | E9PQN2 | Bcl-2-associated transcription factor 1 | 175 | 1028 |  |
| Q13425 | 0,323496 | Phosphoserine | Q13425 | Beta-2-syntrophin | 393 | 1029 |  |
| Q9UQ35 | 0,3236266 | Phosphoserine | Q9UQ35 | Serine/arginine repetitive matrix protein 2 | 1326 | 1030 |  |
| E7EQT4 | 0,3300093 | Phosphoserine | E7EQT4 | Apoptotic chromatin condensation inducer in the nucleus | 964 | 1031 |  |
| Q9Y281 | 0,3301393 | Phosphoserine | Q9Y281 | Cofilin-2 | 3 | 1032 |  |
| H0YCR2 | 0,3319576 | Phosphoserine | H0YCR2 | RalBP1-associated Eps domain-containing protein 1 | 88 | 1033 |  |
| E9PQN2 | 0,3336439 | Phosphoserine | E9PQN2 | Bcl-2-associated transcription factor 1 | 529 | 1034 |  |
| P35579 | 0,3349399 | Phosphoserine | P35579 | Myosin-9 | 1943 | 1035 |  |
| Q09666 | 0,3351988 | Phosphoserine | Q09666 | Neuroblast differentiation-associated protein AHNAK | 5731 | 1036 |  |
| Q9UQ35 | 0,3353283 | Phosphoserine | Q9UQ35 | Serine/arginine repetitive matrix protein 2 | 2449 | 1037 |  |
| E9PQN2 | 0,3366227 | Phosphoserine | E9PQN2 | Bcl-2-associated transcription factor 1 | 656 | 1038 |  |
| E5RIU6 | 0,3388204 | Phosphothreonine\| by CAK | E5RIU6 | Cyclin-dependent kinase 1 | 161 | 1039 |  |
| Q14160 | 0,3392079 | Phosphoserine | Q14160 | Protein scribble homolog | 1486 | 1040 |  |
| P46821 | 0,3425618 | Phosphoserine | P46821 | Microtubule-associated protein 1B;MAP1B heavy chain;MAP1 light chain LC1 | 1256 | 1041 |  |
| P46821 | 0,3425618 | Phosphoserine | P46821 | Microtubule-associated protein 1B;MAP1B heavy chain;MAP1 light chain LC1 | 1265 | 1042 |  |
| A9Z1X7 | 0,3460366 | Phosphoserine | A9Z1X7 | Serine/arginine repetitive matrix protein 1 | 645 | 1043 |  |
| A9Z1X7 | 0,3460366 | Phosphoserine | A9Z1X7 | Serine/arginine repetitive matrix protein 1 | 647 | 1044 |  |
| Q9Y478 | 0,3474497 | Phosphoserine | Q9Y478 | 5'-AMP-activated protein kinase subunit beta-1 | 108 | 1045 |  |
| Q9Y2W1 | 0,347835 | Phosphoserine | Q9Y2W1 | Thyroid hormone receptor-associated protein 3 | 682 | 1046 |  |
| E9PMS6 | 0,3488616 | Phosphoserine | E9PMS6 | LIM domain only protein 7 | 1178 | 1047 |  |
| J3QRS3 | 0,3500156 | Phosphoserine\| by CDC42BP\| CIT\| MLCK\| PAK1\| ROCK1\| ROCK2\| DAPK1\| DAPK2 and ZIPK/DAPK3;Phosphoserine\| by MLCK;Phosphoserine\| by MLCK and ZIPK/DAPK3 | J3QRS3 | Myosin regulatory light chain 12A;Myosin regulatory light chain 12B;Myosin regulatory light polypeptide 9 | 25 | 1048 |  |
| A8MQ02 | 0,351681 | Phosphoserine | A8MQ02 | Afadin | 1756 | 1049 |  |
| P46821 | 0,3518091 | Phosphoserine | P46821 | Microtubule-associated protein 1B;MAP1B heavy chain;MAP1 light chain LC1 | 1389 | 1050 |  |
| Q09666 | 0,3521931 | Phosphoserine | Q09666 | Neuroblast differentiation-associated protein AHNAK | 511 | 1051 |  |
| Q8N8A6 | 0,3533446 | Phosphoserine | Q8N8A6 | ATP-dependent RNA helicase DDX51 | 83 | 1052 |  |
| J3QS11 | 0,3533446 | Phosphothreonine | J3QS11 | Integrin beta-4 | 191 | 1053 |  |
| H0YG16 | 0,3550062 | Phosphoserine\| by AMPK | H0YG16 | Kinesin light chain 1 | 155 | 1054 |  |
| P46821 | 0,3561554 | Phosphoserine | P46821 | Microtubule-associated protein 1B;MAP1B heavy chain;MAP1 light chain LC1 | 1501 | 1055 |  |
| P48960 | 0,3565382 | Phosphoserine | P48960 | CD97 antigen;CD97 antigen subunit alpha;CD97 antigen subunit beta | 831 | 1056 |  |
| P55081 | 0,356921 | Phosphoserine | P55081 | Microfibrillar-associated protein 1 | 116 | 1057 |  |
| P55081 | 0,356921 | Phosphoserine | P55081 | Microfibrillar-associated protein 1 | 118 | 1058 |  |
| A0A087X2D8 | 0,3571762 | Phosphothreonine | A0A087X2D8 | C-Jun-amino-terminal kinase-interacting protein 4 | 217 | 1059 |  |
| A6NKB8 | 0,358451 | Phosphoserine | A6NKB8 | Aminopeptidase B | 7 | 1060 |  |
| Q9Y2W1 | 0,3589607 | Phosphoserine | Q9Y2W1 | Thyroid hormone receptor-associated protein 3 | 243 | 1061 |  |
| H0Y5T1 | 0,3627773 | Phosphoserine | H0Y5T1 | CLIP-associating protein 1 | 381 | 1062 |  |
| O60832 | 0,3627773 | Phosphoserine | O60832 | H/ACA ribonucleoprotein complex subunit 4 | 494 | 1063 |  |
| A9Z1X7 | 0,3640473 | Phosphoserine | A9Z1X7 | Serine/arginine repetitive matrix protein 1 | 606 | 1064 |  |
| A0A087WUT6 | 0,3710122 | Phosphoserine | A0A087WUT6 | Eukaryotic translation initiation factor 5B | 113 | 1065 |  |
| Q16513 | 0,3715175 | Phosphoserine | Q16513 | Serine/threonine-protein kinase N2 | 583 | 1066 |  |
| A0A0D9SEZ9 | 0,3724013 | Phosphoserine | A0A0D9SEZ9 | Eukaryotic translation initiation factor 3 subunit F | 197 | 1067 |  |
| O14974 | 0,3735367 | Phosphoserine\| by NUAK1 | O14974 | Protein phosphatase 1 regulatory subunit 12A | 445 | 1068 |  |
| P00558 | 0,3736627 | Phosphoserine | P00558 | Phosphoglycerate kinase 1 | 203 | 1069 |  |
| P52948 | 0,3765601 | Phosphoserine | P52948 | Nuclear pore complex protein Nup98-Nup96;Nuclear pore complex protein Nup98;Nuclear pore complex protein Nup96 | 888 | 1070 |  |
| H0Y5T1 | 0,376686 | Phosphoserine | H0Y5T1 | CLIP-associating protein 1 | 867 | 1071 |  |
| Q9UQ35 | 0,3771893 | Phosphoserine | Q9UQ35 | Serine/arginine repetitive matrix protein 2 | 1329 | 1072 |  |
| Q96QR8 | 0,3779438 | Phosphoserine | Q96QR8 | Transcriptional activator protein Pur-beta | 101 | 1073 |  |
| A0A087WWP4 | 0,3795773 | Phosphoserine | A0A087WWP4 | Putative RNA-binding protein 15 | 250 | 1074 |  |
| O14974 | 0,3798285 | Phosphoserine | O14974 | Protein phosphatase 1 regulatory subunit 12A | 507 | 1075 |  |
| C9JSU1 | 0,3804561 | Phosphoserine | C9JSU1 | Leucine-rich repeat flightless-interacting protein 2 | 96 | 1076 |  |
| O60343 | 0,3807071 | Phosphoserine\| by PKB/AKT1 | O60343 | TBC1 domain family member 4 | 588 | 1077 |  |
| P05412 | 0,3814598 | Phosphoserine\| by MAPK8 and PLK3 | P05412 | Transcription factor AP-1 | 63 | 1078 |  |
| Q13045 | 0,3833399 | Phosphoserine | Q13045 | Protein flightless-1 homolog | 856 | 1079 |  |
| P05387 | 0,3844666 | Phosphoserine | P05387 | 60S acidic ribosomal protein P2 | 17 | 1080 |  |
| Q5T200 | 0,3853425 | Phosphoserine | Q5T200 | Zinc finger CCCH domain-containing protein 13 | 993 | 1081 |  |
| Q7L8J4 | 0,3853425 | Phosphoserine | Q7L8J4 | SH3 domain-binding protein 5-like | 362 | 1082 |  |
| Q8N6N3 | 0,3855926 | Phosphoserine | Q8N6N3 | UPF0690 protein C1orf52 | 158 | 1083 |  |
| P52948 | 0,3875923 | Phosphoserine | P52948 | Nuclear pore complex protein Nup98-Nup96;Nuclear pore complex protein Nup98;Nuclear pore complex protein Nup96 | 612 | 1084 |  |
| K7ENL0 | 0,3894644 | Phosphoserine | K7ENL0 | Septin-9 | 67 | 1085 |  |
| Q99567 | 0,3900878 | Phosphoserine | Q99567 | Nuclear pore complex protein Nup88 | 517 | 1086 |  |
| B1AUU8 | 0,3925791 | Phosphoserine | B1AUU8 | Epidermal growth factor receptor substrate 15 | 662 | 1087 |  |
| X6RJP6 | 0,3933257 | Phosphoserine | X6RJP6 | Transgelin-2 | 163 | 1088 |  |
| P53621 | 0,3946933 | Phosphoserine | P53621 | Coatomer subunit alpha;Xenin;Proxenin | 173 | 1089 |  |
| Q99575 | 0,3949419 | Phosphoserine | Q99575 | Ribonucleases P/MRP protein subunit POP1 | 730 | 1090 |  |
| J3QRS3 | 0,3953145 | Phosphothreonine\| by MLCK;Phosphothreonine\| by MLCK and ZIPK/DAPK3;Phosphothreonine\| by MLCK\| CIT and ROCK2 | J3QRS3 | Myosin regulatory light chain 12A;Myosin regulatory light chain 12B;Myosin regulatory light polypeptide 9 | 24 | 1091 |  |
| A0A0C4DGV5 | 0,3960596 | Phosphoserine | A0A0C4DGV5 | Zinc finger Ran-binding domain-containing protein 2 | 120 | 1092 |  |
| P80723 | 0,3965562 | Phosphothreonine | P80723 | Brain acid soluble protein 1 | 31 | 1093 |  |
| E9PHV5 | 0,4038604 | Phosphoserine | E9PHV5 | Sperm-specific antigen 2 | 92 | 1094 |  |
| H0YG16 | 0,4042308 | Phosphoserine\| by AMPK | H0YG16 | Kinesin light chain 1 | 152 | 1095 |  |
| Q7L2J0 | 0,4052181 | Phosphoserine | Q7L2J0 | 7SK snRNA methylphosphate capping enzyme | 152 | 1096 |  |
| E7EVA0 | 0,4071907 | Phosphoserine | E7EVA0 | Microtubule-associated protein;Microtubule-associated protein 4 | 2073 | 1097 |  |
| Q9C0C2 | 0,4079297 | Phosphoserine | Q9C0C2 | 182 kDa tankyrase-1-binding protein | 429 | 1098 |  |
| Q9UDY2 | 0,4090376 | Phosphoserine | Q9UDY2 | Tight junction protein ZO-2 | 244 | 1099 |  |
| E5RJR5 | 0,4097756 | Phosphothreonine | E5RJR5 | S-phase kinase-associated protein 1 | 131 | 1100 |  |
| Q5JRI1 | 0,4172581 | Phosphoserine | Q5JRI1 | Serine/arginine-rich splicing factor 10 | 133 | 1101 |  |
| Q5JSH3 | 0,4183587 | Phosphoserine | Q5JSH3 | WD repeat-containing protein 44 | 96 | 1102 |  |
| A9Z1X7 | 0,4194586 | Phosphoserine | A9Z1X7 | Serine/arginine repetitive matrix protein 1 | 447 | 1103 |  |
| E7EN95 | 0,4209237 | Phosphoserine | E7EN95 | Filamin-B | 1336 | 1104 |  |
| E9PQN2 | 0,4217777 | Phosphoserine | E9PQN2 | Bcl-2-associated transcription factor 1 | 383 | 1105 |  |
| Q9Y2W1 | 0,4226311 | Phosphoserine | Q9Y2W1 | Thyroid hormone receptor-associated protein 3 | 379 | 1106 |  |
| Q9UK58 | 0,4253103 | Phosphoserine | Q9UK58 | Cyclin-L1 | 445 | 1107 |  |
| O43815 | 0,4264049 | Phosphoserine | O43815 | Striatin | 245 | 1108 |  |
| O95684 | 0,4271341 | Phosphoserine | O95684 | FGFR1 oncogene partner | 156 | 1109 |  |
| O95684 | 0,4271341 | Phosphoserine | O95684 | FGFR1 oncogene partner | 160 | 1110 |  |
| B7ZKW8 | 0,4281059 | Phosphoserine | B7ZKW8 | CapZ-interacting protein | 147 | 1111 |  |
| V9GYZ0 | 0,4293197 | Phosphoserine | V9GYZ0 | Pantothenate kinase 2, mitochondrial | 66 | 1112 |  |
| A0A0A0MR52 | 0,4318653 | Phosphoserine | A0A0A0MR52 | Eukaryotic translation initiation factor 4 gamma 1 | 988 | 1113 |  |
| Q14C86 | 0,4325918 | Phosphoserine | Q14C86 | GTPase-activating protein and VPS9 domain-containing protein 1 | 902 | 1114 |  |
| H0Y5T1 | 0,4329549 | Phosphoserine | H0Y5T1 | CLIP-associating protein 1 | 933 | 1115 |  |
| Q13242 | 0,4336809 | Phosphoserine | Q13242 | Serine/arginine-rich splicing factor 9 | 189 | 1116 |  |
| P42677 | 0,4338018 | Phosphoserine | P42677 | 40S ribosomal protein S27 | 11 | 1117 |  |
| J3KSR8 | 0,435615 | Phosphoserine | J3KSR8 | Serine/arginine-rich splicing factor 1 | 94 | 1118 |  |
| J3KSR8 | 0,435615 | Phosphoserine | J3KSR8 | Serine/arginine-rich splicing factor 1 | 96 | 1119 |  |
| Q9Y2X3 | 0,4412813 | Phosphoserine | Q9Y2X3 | Nucleolar protein 58 | 502 | 1120 |  |
| P07900 | 0,4414016 | Phosphoserine | P07900 | Heat shock protein HSP 90-alpha | 263 | 1121 |  |
| Q5TCC6 | 0,4453665 | Phosphoserine | Q5TCC6 | Isoleucine--tRNA ligase, cytoplasmic | 100 | 1122 |  |
| Q9BUH6 | 0,4454865 | Phosphoserine | Q9BUH6 | Uncharacterized protein C9orf142 | 148 | 1123 |  |
| Q9UQ35 | 0,4523089 | Phosphoserine | Q9UQ35 | Serine/arginine repetitive matrix protein 2 | 1101 | 1124 |  |
| Q01804 | 0,4524283 | Phosphoserine | Q01804 | OTU domain-containing protein 4 | 1024 | 1125 |  |
| Q09666 | 0,454576 | Phosphoserine | Q09666 | Neuroblast differentiation-associated protein AHNAK | 210 | 1126 |  |
| Q09666 | 0,454576 | Phosphoserine | Q09666 | Neuroblast differentiation-associated protein AHNAK | 216 | 1127 |  |
| E9PQN2 | 0,4560058 | Phosphoserine | E9PQN2 | Bcl-2-associated transcription factor 1 | 658 | 1128 |  |
| Q9BXP5 | 0,456125 | Phosphoserine | Q9BXP5 | Serrate RNA effector molecule homolog | 74 | 1129 |  |
| Q5JPT2 | 0,4573153 | Phosphoserine | Q5JPT2 | SH3 domain-containing kinase-binding protein 1 | 567 | 1130 |  |
| E9PQN2 | 0,4601684 | Phosphoserine | E9PQN2 | Bcl-2-associated transcription factor 1 | 179 | 1131 |  |
| Q8WWQ0 | 0,4626602 | Phosphoserine | Q8WWQ0 | PH-interacting protein | 1315 | 1132 |  |
| Q9NQC3 | 0,4669219 | Phosphoserine | Q9NQC3 | Reticulon-4 | 181 | 1133 |  |
| E9PG73 | 0,4686939 | Phosphoserine | E9PG73 | Peptidyl-prolyl cis-trans isomerase G | 672 | 1134 |  |
| P08670 | 0,4697561 | Phosphoserine | P08670 | Vimentin | 430 | 1135 |  |
| Q08211 | 0,4702278 | Phosphoserine | Q08211 | ATP-dependent RNA helicase A | 321 | 1136 |  |
| E7EQT4 | 0,4723492 | Phosphoserine | E7EQT4 | Apoptotic chromatin condensation inducer in the nucleus | 670 | 1137 |  |
| C9JIG9 | 0,4743496 | Phosphoserine | C9JIG9 | Serine/threonine-protein kinase OSR1 | 339 | 1138 |  |
| P14618 | 0,4749376 | Phosphoserine | P14618 | Pyruvate kinase PKM;Pyruvate kinase | 37 | 1139 |  |
| Q16543 | 0,4776386 | Phosphoserine | Q16543 | Hsp90 co-chaperone Cdc37;Hsp90 co-chaperone Cdc37, N-terminally processed | 13 | 1140 |  |
| D6REM6 | 0,4779906 | Phosphoserine | D6REM6 | Matrin-3 | 188 | 1141 |  |
| Q08AD1 | 0,478108 | Phosphoserine | Q08AD1 | Calmodulin-regulated spectrin-associated protein 2 | 464 | 1142 |  |
| Q9Y5K6 | 0,4792803 | Phosphoserine | Q9Y5K6 | CD2-associated protein | 510 | 1143 |  |
| E5RJY1 | 0,4809201 | Phosphoserine\| by SGK1 | E5RJY1 | Protein NDRG1 | 39 | 1144 |  |
| A9Z1X7 | 0,4812714 | Phosphoserine | A9Z1X7 | Serine/arginine repetitive matrix protein 1 | 883 | 1145 |  |
| Q9UQ35 | 0,4813884 | Phosphoserine | Q9UQ35 | Serine/arginine repetitive matrix protein 2 | 1103 | 1146 |  |
| Q9UQ35 | 0,4819734 | Phosphoserine | Q9UQ35 | Serine/arginine repetitive matrix protein 2 | 1014 | 1147 |  |
| M0R300 | 0,4824413 | Phosphothreonine | M0R300 | Unconventional myosin-IXb | 1346 | 1148 |  |
| A9Z1X7 | 0,4836101 | Phosphoserine | A9Z1X7 | Serine/arginine repetitive matrix protein 1 | 458 | 1149 |  |
| A9Z1X7 | 0,4836101 | Phosphoserine | A9Z1X7 | Serine/arginine repetitive matrix protein 1 | 460 | 1150 |  |
| H7BXI1 | 0,485245 | Phosphoserine | H7BXI1 | Extended synaptotagmin-2 | 718 | 1151 |  |
| D6REM6 | 0,4865284 | Phosphoserine | D6REM6 | Matrin-3 | 195 | 1152 |  |
| Q9NTI5 | 0,4880435 | Phosphoserine | Q9NTI5 | Sister chromatid cohesion protein PDS5 homolog B | 1358 | 1153 |  |
| Q09666 | 0,489208 | Phosphoserine | Q09666 | Neuroblast differentiation-associated protein AHNAK | 5780 | 1154 |  |
| E9PG73 | 0,4904876 | Phosphoserine | E9PG73 | Peptidyl-prolyl cis-trans isomerase G | 382 | 1155 |  |
| Q9Y2W1 | 0,494784 | Phosphoserine | Q9Y2W1 | Thyroid hormone receptor-associated protein 3 | 939 | 1156 |  |
| Q05209 | 0,4953636 | Phosphoserine | Q05209 | Tyrosine-protein phosphatase non-receptor type 12 | 435 | 1157 |  |
| Q5VU10 | 0,4973325 | Phosphoserine | Q5VU10 | Ribonuclease P protein subunit p30 | 195 | 1158 |  |
| A0A0C4DGV5 | 0,4974483 | Phosphoserine | A0A0C4DGV5 | Zinc finger Ran-binding domain-containing protein 2 | 188 | 1159 |  |
| H3BTK3 | 0,5004541 | Phosphoserine | H3BTK3 | Calcium-regulated heat stable protein 1 | 52 | 1160 |  |
| P62995 | 0,5029926 | Phosphoserine | P62995 | Transformer-2 protein homolog beta | 14 | 1161 |  |
| E9PQN2 | 0,5074819 | Phosphoserine | E9PQN2 | Bcl-2-associated transcription factor 1 | 494 | 1162 |  |
| P46821 | 0,5077116 | Phosphoserine | P46821 | Microtubule-associated protein 1B;MAP1B heavy chain;MAP1 light chain LC1 | 1785 | 1163 |  |
| Q9UDY2 | 0,508401 | Phosphoserine | Q9UDY2 | Tight junction protein ZO-2 | 174 | 1164 |  |
| Q9BX95 | 0,5090899 | Phosphoserine | Q9BX95 | Sphingosine-1-phosphate phosphatase 1 | 112 | 1165 |  |
| M0R300 | 0,5106963 | Phosphoserine | M0R300 | Unconventional myosin-IXb | 1354 | 1166 |  |
| H0YEN2 | 0,5141324 | Phosphoserine | H0YEN2 | Serine/threonine-protein phosphatase 6 regulatory subunit 3 | 286 | 1167 |  |
| Q08AD1 | 0,5142466 | Phosphoserine | Q08AD1 | Calmodulin-regulated spectrin-associated protein 2 | 1319 | 1168 |  |
| O14974 | 0,5153902 | Phosphoserine | O14974 | Protein phosphatase 1 regulatory subunit 12A | 871 | 1169 |  |
| Q9UQ35 | 0,5250731 | Phosphoserine | Q9UQ35 | Serine/arginine repetitive matrix protein 2 | 1103 | 1170 |  |
| B7ZKW8 | 0,5282485 | Phosphoserine\| by MAPKAPK2 and MAPKAPK3 | B7ZKW8 | CapZ-interacting protein | 149 | 1171 |  |
| S4R3V8 | 0,5299468 | Phosphoserine | S4R3V8 | Lipolysis-stimulated lipoprotein receptor | 445 | 1172 |  |
| Q9NTI5 | 0,5311911 | Phosphoserine | Q9NTI5 | Sister chromatid cohesion protein PDS5 homolog B | 1257 | 1173 |  |
| B8ZZQ6 | 0,5329987 | Phosphoserine | B8ZZQ6 | Prothymosin alpha;Prothymosin alpha, N-terminally processed;Thymosin alpha-1 | 10 | 1174 |  |
| J3KSH8 | 0,5333375 | Phosphoserine | J3KSH8 | Hematological and neurological expressed 1 protein;Hematological and neurological expressed 1 protein, N-terminally processed | 41 | 1175 |  |
| Q9BSJ8 | 0,5404319 | Phosphoserine | Q9BSJ8 | Extended synaptotagmin-1 | 1034 | 1176 |  |
| Q9UPU5 | 0,5451421 | Phosphoserine | Q9UPU5 | Ubiquitin carboxyl-terminal hydrolase 24 | 2561 | 1177 |  |
| O75822 | 0,5464852 | Phosphoserine | O75822 | Eukaryotic translation initiation factor 3 subunit J | 11 | 1178 |  |
| Q15149 | 0,5477151 | Phosphoserine | Q15149 | Plectin | 720 | 1179 |  |
| A0A0A0MT60 | 0,5499487 | Phosphoserine | A0A0A0MT60 | FK506-binding protein 15 | 981 | 1180 |  |
| A9Z1X7 | 0,5512873 | Phosphoserine | A9Z1X7 | Serine/arginine repetitive matrix protein 1 | 747 | 1181 |  |
| A9Z1X7 | 0,5512873 | Phosphoserine | A9Z1X7 | Serine/arginine repetitive matrix protein 1 | 749 | 1182 |  |
| P13861 | 0,5588487 | Phosphoserine | P13861 | cAMP-dependent protein kinase type II-alpha regulatory subunit | 78 | 1183 |  |
| P13861 | 0,5588487 | Phosphoserine | P13861 | cAMP-dependent protein kinase type II-alpha regulatory subunit | 80 | 1184 |  |
| Q9BX95 | 0,5608438 | Phosphothreonine | Q9BX95 | Sphingosine-1-phosphate phosphatase 1 | 114 | 1185 |  |
| Q7L8J4 | 0,5618403 | Phosphoserine | Q7L8J4 | SH3 domain-binding protein 5-like | 358 | 1186 |  |
| Q99567 | 0,5669224 | Phosphoserine | Q99567 | Nuclear pore complex protein Nup88 | 35 | 1187 |  |
| Q9Y2W1 | 0,5690163 | Phosphothreonine | Q9Y2W1 | Thyroid hormone receptor-associated protein 3 | 874 | 1188 |  |
| E9PG73 | 0,571767 | Phosphoserine | E9PG73 | Peptidyl-prolyl cis-trans isomerase G | 675 | 1189 |  |
| P51858 | 0,5753897 | Phosphoserine | P51858 | Hepatoma-derived growth factor | 132 | 1190 |  |
| P51858 | 0,5753897 | Phosphoserine | P51858 | Hepatoma-derived growth factor | 133 | 1191 |  |
| P68363 | 0,5779094 | Phosphoserine | P68363 | Tubulin alpha-1C chain;Tubulin alpha-3C/D chain;Tubulin alpha-3E chain;Tubulin alpha-1B chain;Tubulin alpha-1A chain | 48 | 1192 |  |
| C9JA08 | 0,5783472 | Phosphoserine | C9JA08 | 60S ribosomal export protein NMD3 | 468 | 1193 |  |
| Q9UK58 | 0,5783472 | Phosphoserine | Q9UK58 | Cyclin-L1 | 352 | 1194 |  |
| E7EX17 | 0,5788942 | Phosphoserine | E7EX17 | Eukaryotic translation initiation factor 4B | 406 | 1195 |  |
| Q9NWB6 | 0,586204 | Phosphoserine | Q9NWB6 | Arginine and glutamate-rich protein 1 | 77 | 1196 |  |
| P04792 | 0,5901164 | Phosphoserine\| by MAPKAPK2\| MAPKAPK3 and MAPKAPK5 | P04792 | Heat shock protein beta-1 | 82 | 1197 |  |
| Q09666 | 0,5914181 | Phosphoserine | Q09666 | Neuroblast differentiation-associated protein AHNAK | 135 | 1198 |  |
| B7ZKW8 | 0,5929354 | Phosphoserine\| by MAPK8\| in vitro | B7ZKW8 | CapZ-interacting protein | 53 | 1199 |  |
| Q09666 | 0,5953164 | Phosphoserine | Q09666 | Neuroblast differentiation-associated protein AHNAK | 3426 | 1200 |  |
| A0A087WUT6 | 0,5961813 | Phosphoserine | A0A087WUT6 | Eukaryotic translation initiation factor 5B | 107 | 1201 |  |
| E7EN95 | 0,6039418 | Phosphoserine | E7EN95 | Filamin-B | 2285 | 1202 |  |
| F2Z2T1 | 0,605768 | Phosphoserine\| by RPS6KB1 | F2Z2T1 | Nuclear cap-binding protein subunit 1 | 22 | 1203 |  |
| E9PMS6 | 0,6086639 | Phosphoserine | E9PMS6 | LIM domain only protein 7 | 155 | 1204 |  |
| H0Y2V6 | 0,609949 | Phosphoserine | H0Y2V6 | Centrosomal protein of 170 kDa;Perilipin-5 | 1124 | 1205 |  |
| Q9UQ35 | 0,6129432 | Phosphoserine | Q9UQ35 | Serine/arginine repetitive matrix protein 2 | 1102 | 1206 |  |
| A9Z1X7 | 0,6139044 | Phosphoserine | A9Z1X7 | Serine/arginine repetitive matrix protein 1 | 625 | 1207 |  |
| P30050 | 0,6184876 | Phosphoserine | P30050 | 60S ribosomal protein L12 | 38 | 1208 |  |
| P18206 | 0,621995 | Phosphoserine | P18206 | Vinculin | 290 | 1209 |  |
| Q15785 | 0,6240107 | Phosphoserine | Q15785 | Mitochondrial import receptor subunit TOM34 | 186 | 1210 |  |
| M0R1Q8 | 0,6245406 | Phosphoserine | M0R1Q8 | Regulator of chromosome condensation | 113 | 1211 |  |
| P53396 | 0,6252823 | Phosphoserine\| by PKA and PKB/AKT1 or PKB/AKT2 | P53396 | ATP-citrate synthase | 455 | 1212 |  |
| E9PPU1 | 0,6306739 | Phosphothreonine | E9PPU1 | 40S ribosomal protein S3 | 136 | 1213 |  |
| B1AKZ5 | 0,630885 | Phosphoserine | B1AKZ5 | Astrocytic phosphoprotein PEA-15 | 94 | 1214 |  |
| A9Z1X7 | 0,6310959 | Phosphothreonine | A9Z1X7 | Serine/arginine repetitive matrix protein 1 | 881 | 1215 |  |
| Q53EU6 | 0,6316234 | Phosphoserine | Q53EU6 | Glycerol-3-phosphate acyltransferase 3 | 68 | 1216 |  |
| Q0JRZ9 | 0,6353095 | Phosphoserine | Q0JRZ9 | FCH domain only protein 2 | 403 | 1217 |  |
| V9GYM8 | 0,6409782 | Phosphoserine | V9GYM8 | Rho guanine nucleotide exchange factor 2 | 690 | 1218 |  |
| P50914 | 0,6474594 | Phosphoserine | P50914 | 60S ribosomal protein L14 | 139 | 1219 |  |
| P49006 | 0,6476679 | Phosphothreonine | P49006 | MARCKS-related protein | 148 | 1220 |  |
| Q9C0C2 | 0,6562979 | Phosphoserine | Q9C0C2 | 182 kDa tankyrase-1-binding protein | 1620 | 1221 |  |
| P26038 | 0,657127 | Phosphothreonine\| by ROCK2;Phosphothreonine\| by ROCK2 and PKC/PRKCI;Phosphothreonine\| by ROCK2 and STK10 | P26038 | Ezrin;Moesin;Radixin | 558 | 1222 |  |
| O00264 | 0,6627108 | Phosphoserine | O00264 | Membrane-associated progesterone receptor component 1 | 57 | 1223 |  |
| F8W6I7 | 0,6662154 | Phosphoserine | F8W6I7 | Heterogeneous nuclear ribonucleoprotein A1;Heterogeneous nuclear ribonucleoprotein A1, N-terminally processed | 6 | 1224 |  |
| P30086 | 0,6663184 | Phosphoserine | P30086 | Phosphatidylethanolamine-binding protein 1;Hippocampal cholinergic neurostimulating peptide | 52 | 1225 |  |
| Q9H1E3 | 0,6665243 | Phosphoserine | Q9H1E3 | Nuclear ubiquitous casein and cyclin-dependent kinase substrate 1 | 79 | 1226 |  |
| G5E9Q4 | 0,6670388 | Phosphoserine | G5E9Q4 | Interferon-inducible double-stranded RNA-dependent protein kinase activator A | 18 | 1227 |  |
| J3KSH8 | 0,6713539 | Phosphoserine | J3KSH8 | Hematological and neurological expressed 1 protein;Hematological and neurological expressed 1 protein, N-terminally processed | 42 | 1228 |  |
| M0R2U2 | 0,6713539 | Phosphoserine | M0R2U2 | rRNA 2'-O-methyltransferase fibrillarin | 111 | 1229 |  |
| P13861 | 0,675554 | Phosphoserine\| by PKA | P13861 | cAMP-dependent protein kinase type II-alpha regulatory subunit | 99 | 1230 |  |
| E7EX17 | 0,6774962 | Phosphoserine | E7EX17 | Eukaryotic translation initiation factor 4B | 93 | 1231 |  |
| H0Y449 | 0,6826966 | Phosphoserine;Phosphoserine\| by PKB/AKT1 | H0Y449 | Nuclease-sensitive element-binding protein 1;Y-box-binding protein 2;Y-box-binding protein 3 | 152 | 1232 |  |
| E9PNJ4 | 0,6829002 | Phosphoserine | E9PNJ4 | Stromal interaction molecule 1 | 84 | 1233 |  |
| Q09666 | 0,6867622 | Phosphoserine | Q09666 | Neuroblast differentiation-associated protein AHNAK | 5749 | 1234 |  |
| A0A087X055 | 0,6880811 | Phosphoserine | A0A087X055 | Ras-responsive element-binding protein 1 | 236 | 1235 |  |
| P11387 | 0,6910189 | Phosphoserine | P11387 | DNA topoisomerase 1 | 10 | 1236 |  |
| P52948 | 0,6931425 | Phosphoserine | P52948 | Nuclear pore complex protein Nup98-Nup96;Nuclear pore complex protein Nup98;Nuclear pore complex protein Nup96 | 1028 | 1237 |  |
| Q9H1E3 | 0,6939507 | Phosphothreonine | Q9H1E3 | Nuclear ubiquitous casein and cyclin-dependent kinase substrate 1 | 179 | 1238 |  |
| P18206 | 0,7016059 | Phosphoserine | P18206 | Vinculin | 721 | 1239 |  |
| Q9UQ35 | 0,7078208 | Phosphoserine | Q9UQ35 | Serine/arginine repetitive matrix protein 2 | 2694 | 1240 |  |
| D6RD83 | 0,7079209 | Phosphoserine | D6RD83 | Heterogeneous nuclear ribonucleoprotein D0 | 37 | 1241 |  |
| F2Z2T1 | 0,7091208 | Phosphothreonine\| by RPS6KB1 | F2Z2T1 | Nuclear cap-binding protein subunit 1 | 21 | 1242 |  |
| A0A087WZH7 | 0,7112179 | Phosphoserine\| by PKC | A0A087WZH7 | Myristoylated alanine-rich C-kinase substrate | 170 | 1243 |  |
| P18031 | 0,7150049 | Phosphoserine\| by PKB/AKT1\| CLK1 and CLK2 | P18031 | Tyrosine-protein phosphatase non-receptor type 1 | 50 | 1244 |  |
| Q9NS69 | 0,7271956 | Phosphoserine | Q9NS69 | Mitochondrial import receptor subunit TOM22 homolog | 15 | 1245 |  |
| E7EN38 | 0,7282808 | Phosphothreonine | E7EN38 | DNA repair protein RAD50 | 629 | 1246 |  |
| A9Z1X7 | 0,7305472 | Phosphoserine | A9Z1X7 | Serine/arginine repetitive matrix protein 1 | 747 | 1247 |  |
| D6RD83 | 0,7312363 | Phosphoserine | D6RD83 | Heterogeneous nuclear ribonucleoprotein D0 | 36 | 1248 |  |
| Q09666 | 0,7333998 | Phosphoserine | Q09666 | Neuroblast differentiation-associated protein AHNAK | 210 | 1249 |  |
| Q7Z417 | 0,7348731 | Phosphoserine | Q7Z417 | Nuclear fragile X mental retardation-interacting protein 2 | 652 | 1250 |  |
| P46821 | 0,7369332 | Phosphoserine | P46821 | Microtubule-associated protein 1B;MAP1B heavy chain;MAP1 light chain LC1 | 1779 | 1251 |  |
| H0Y2V6 | 0,7377172 | Phosphoserine | H0Y2V6 | Centrosomal protein of 170 kDa;Perilipin-5 | 1124 | 1252 |  |
| K7ES59 | 0,7423148 | Phosphoserine | K7ES59 | Polypyrimidine tract-binding protein 1 | 111 | 1253 |  |
| P08670 | 0,7435839 | Phosphothreonine | P08670 | Vimentin | 426 | 1254 |  |
| P06748 | 0,7543746 | Phosphoserine | P06748 | Nucleophosmin | 243 | 1255 |  |
| P17252 | 0,7557299 | Phosphoserine | P17252 | Protein kinase C alpha type | 226 | 1256 |  |
| Q9NZM1 | 0,7583401 | Phosphoserine | Q9NZM1 | Myoferlin | 174 | 1257 |  |
| Q5TCC6 | 0,7745713 | Phosphoserine | Q5TCC6 | Isoleucine--tRNA ligase, cytoplasmic | 102 | 1258 |  |
| Q5T757 | 0,7822865 | Phosphoserine | Q5T757 | Serine/arginine-rich splicing factor 11 | 147 | 1259 |  |
| F8W6I7 | 0,7904331 | Phosphoserine | F8W6I7 | Heterogeneous nuclear ribonucleoprotein A1;Heterogeneous nuclear ribonucleoprotein A1, N-terminally processed | 4 | 1260 |  |
| Q8TCJ2 | 0,79656 | Phosphoserine | Q8TCJ2 | Dolichyl-diphosphooligosaccharide--protein glycosyltransferase subunit STT3B | 498 | 1261 |  |
| Q9C0C2 | 0,7999422 | Phosphoserine | Q9C0C2 | 182 kDa tankyrase-1-binding protein | 836 | 1262 |  |
| M0QYL4 | 0,8039717 | Phosphoserine | M0QYL4 | A-kinase anchor protein 8 | 142 | 1263 |  |
| Q14151 | 0,805842 | Phosphoserine | Q14151 | Scaffold attachment factor B2 | 513 | 1264 |  |
| H3BPE1 | 0,8064026 | Phosphoserine | H3BPE1 | Microtubule-actin cross-linking factor 1, isoforms 1/2/3/5 | 275 | 1265 |  |
| Q96CV9 | 0,8152504 | Phosphoserine\| by TBK1 | Q96CV9 | Optineurin | 177 | 1266 |  |
| E9PD43 | 0,8154361 | Phosphoserine | E9PD43 | Negative elongation factor E | 115 | 1267 |  |
| M0QY76 | 0,8269944 | Phosphoserine\| by MAPKAPK2 | M0QY76 | Tristetraprolin | 192 | 1268 |  |
| O95239 | 0,8306736 | Phosphoserine | O95239 | Chromosome-associated kinesin KIF4A | 801 | 1269 |  |
| A0A087WTP3 | 0,8340685 | Phosphoserine | A0A087WTP3 | Far upstream element-binding protein 2 | 181 | 1270 |  |
| Q5HY54 | 0,8404695 | Phosphoserine | Q5HY54 | Filamin-A | 1459 | 1271 |  |
| A9Z1X7 | 0,8452972 | Phosphoserine | A9Z1X7 | Serine/arginine repetitive matrix protein 1 | 458 | 1272 |  |
| E7ETM8 | 0,846388 | Phosphoserine | E7ETM8 | Nexilin | 301 | 1273 |  |
| O00264 | 0,8508336 | Phosphoserine | O00264 | Membrane-associated progesterone receptor component 1 | 54 | 1274 |  |
| P48651 | 0,860854 | Phosphoserine | P48651 | Phosphatidylserine synthase 1 | 417 | 1275 |  |
| C9JZW3 | 0,8646273 | Phosphoserine | C9JZW3 | Elongation factor 1-beta | 106 | 1276 |  |
| A0A087WW66 | 0,8765117 | Phosphoserine | A0A087WW66 | 26S proteasome non-ATPase regulatory subunit 1 | 315 | 1277 |  |
| A0A087WW66 | 0,8765117 | Phosphothreonine | A0A087WW66 | 26S proteasome non-ATPase regulatory subunit 1 | 311 | 1278 |  |
| C9JXZ5 | 0,8805107 | Phosphoserine | C9JXZ5 | Vesicle-associated membrane protein 8 | 29 | 1279 |  |
| J3KSR8 | 0,8905039 | Phosphoserine | J3KSR8 | Serine/arginine-rich splicing factor 1 | 94 | 1280 |  |
| P51858 | 0,8927056 | Phosphoserine | P51858 | Hepatoma-derived growth factor | 132 | 1281 |  |
| F5H2U8 | 0,8934094 | Phosphoserine | F5H2U8 | High mobility group protein HMGI-C | 44 | 1282 |  |
| P17812 | 0,8979756 | Phosphoserine | P17812 | CTP synthase 1 | 575 | 1283 |  |
| H0YE82 | 0,9081103 | Phosphoserine | H0YE82 | N-acetyl-D-glucosamine kinase | 25 | 1284 |  |
| M0R0F9 | 0,9229215 | Phosphoserine | M0R0F9 | Cdc42-interacting protein 4 | 318 | 1285 |  |
| H0Y3K4 | 0,9254183 | Phosphoserine | H0Y3K4 | RNA-binding protein 33 | 537 | 1286 |  |
| Q8TCJ2 | 0,9291984 | Phosphoserine | Q8TCJ2 | Dolichyl-diphosphooligosaccharide--protein glycosyltransferase subunit STT3B | 498 | 1287 |  |
| Q8TCJ2 | 0,9291984 | Phosphoserine | Q8TCJ2 | Dolichyl-diphosphooligosaccharide--protein glycosyltransferase subunit STT3B | 499 | 1288 |  |
| E5RJY1 | 0,9301419 | Phosphothreonine\| by SGK1 | E5RJY1 | Protein NDRG1 | 75 | 1289 |  |
| O00567 | 0,9460872 | Phosphoserine | O00567 | Nucleolar protein 56 | 563 | 1290 |  |
| Q9NXG2 | 0,9491367 | Phosphoserine | Q9NXG2 | THUMP domain-containing protein 1 | 86 | 1291 |  |
| E9PDE4 | 0,9629482 | Phosphoserine | E9PDE4 | Calpastatin | 292 | 1292 |  |
| O43399 | 0,9653767 | Phosphoserine | O43399 | Tumor protein D54 | 166 | 1293 |  |
| P51858 | 0,9942871 | Phosphoserine | P51858 | Hepatoma-derived growth factor | 133 | 1294 |  |
| H0Y6S1 | 1,006212 | Phosphoserine | H0Y6S1 | Sex comb on midleg-like protein 2 | 15 | 1295 |  |
| H0Y6S1 | 1,006212 | Phosphoserine | H0Y6S1 | Sex comb on midleg-like protein 2 | 27 | 1296 |  |
| P05387 | 1,007919 | Phosphoserine | P05387 | 60S acidic ribosomal protein P2;60S acidic ribosomal protein P1 | 105 | 1297 |  |
| H3BM89 | 1,013514 | Phosphoserine | H3BM89 | 60S ribosomal protein L4 | 201 | 1298 |  |
| Q9NXV6 | 1,047346 | Phosphoserine | Q9NXV6 | CDKN2A-interacting protein | 131 | 1299 |  |
| P28482 | 1,077 | Phosphotyrosine\| by MAP2K1 and MAP2K2 | P28482 | Mitogen-activated protein kinase 1 | 187 | 1300 |  |
| F2Z2K0 | 1,081407 | Phosphoserine | F2Z2K0 | NSFL1 cofactor p47 | 114 | 1301 |  |
| H7C5W9 | 1,091486 | Phosphoserine | H7C5W9 | Sarcoplasmic/endoplasmic reticulum calcium ATPase 2 | 554 | 1302 |  |
| F8W726 | 1,130366 | Phosphoserine | F8W726 | Ubiquitin-associated protein 2-like | 620 | 1303 |  |
| F5H0B0 | 1,141962 | Phosphothreonine | F5H0B0 | Tumor protein D52 | 173 | 1304 |  |
| Q8ND56 | 1,16196 | Phosphoserine | Q8ND56 | Protein LSM14 homolog A | 216 | 1305 |  |
| P07355 | 1,180531 | Phosphotyrosine\| by SRC | P07355 | Annexin A2;Annexin;Putative annexin A2-like protein | 24 | 1306 |  |
| Q14315 | 1,228732 | Phosphoserine | Q14315 | Filamin-C | 2233 | 1307 |  |
| P02545 | 1,23631 | Phosphoserine | P02545 | Prelamin-A/C;Lamin-A/C | 392 | 1308 |  |
| J3QL05 | 1,281973 | Phosphoserine | J3QL05 | Serine/arginine-rich splicing factor 2 | 26 | 1309 |  |
| F8W930 | 1,297676 | Phosphoserine | F8W930 | Insulin-like growth factor 2 mRNA-binding protein 2 | 170 | 1310 |  |
| Q9H307 | 1,350272 | Phosphoserine | Q9H307 | Pinin | 66 | 1311 |  |
| Q96PK6 | 1,372843 | Phosphoserine | Q96PK6 | RNA-binding protein 14 | 618 | 1312 |  |
| M0R300 | 1,380958 | Phosphoserine | M0R300 | Unconventional myosin-IXb | 1405 | 1313 |  |
| H0YN26 | 1,423484 | Phosphoserine | H0YN26 | Acidic leucine-rich nuclear phosphoprotein 32 family member A | 17 | 1314 |  |
| Q15084 | 1,452727 | Phosphoserine | Q15084 | Protein disulfide-isomerase A6 | 428 | 1315 |  |
| F8WF69 | 1,465148 | Phosphoserine | F8WF69 | Clathrin light chain A | 105 | 1316 |  |
| H0YB56 | 1,489441 | Phosphoserine | H0YB56 | Protein LYRIC | 181 | 1317 |  |
| Q09666 | 1,531756 | Phosphoserine | Q09666 | Neuroblast differentiation-associated protein AHNAK | 5830 | 1318 |  |
| Q9UQ35 | 1,563057 | Phosphoserine | Q9UQ35 | Serine/arginine repetitive matrix protein 2 | 2694 | 1319 |  |
| P34910 | 1,76268 | Phosphoserine | P34910 | Protein EVI2B | 294 | 1320 |  |
| Q92615 | 2,030049 | Phosphoserine | Q92615 | La-related protein 4B | 244 | 1321 |  |
| E7EPK0 | 2,333385 | Phosphoserine | E7EPK0 | LIM and calponin homology domains-containing protein 1 | 218 | 1322 |  |
| E7ETY4 | 2,373064 | Phosphothreonine\| by PKC/PRKCZ | E7ETY4 | Serine/threonine-protein kinase MARK2 | 516 | 1323 |  |
