## Supplementray Table2 for "The serine/threonine kinase MINK1 directly regulates the function of promigratory proteins"

**Supplementary Table 2:** List of breast cancer data sets included in the study

| **Reference** | **Source of data** | **N° of samples** | **Technological platform** | **N° of probe sets** | **N° of samples used in the present study** |
| --- | --- | --- | --- | --- | --- |
| Expression Project for Oncology (expO), 2005 | https://expo.intgen.org/geo GEO: GSE2109 | 348 | Affymetrix U133 Plus 2.0 | 54K | 348 |
| Ivshina et al.,  Cancer Res 2006 | GEO: GSE4922, GSE1456 | 448 | Affymetrix  U133 A+B | 2x22K | 448 |
| Bonnefoi et al.,  Lancet Oncol 2007 | GEO: GSE6861, GSE4779 | 161 | Affymetrix X3P | 61K | 125 |
| Bos et al., Nature 2009 | GEO: GSE12276 | 204 | Affymetrix U133 Plus 2.0 | 54K | 204 |
| Hoeflich et al.,  Clin Cancer Res 2009 | GEO: GSE12763 | 30 | Affymetrix U133 Plus 2.0 | 54K | 30 |
| Marty et al.,  Breast Cancer Res 2008 | GEO: GSE13787 | 23 | Affymetrix U133 Plus 2.0 | 54K | 23 |
| Barry et al.,  J Clin Oncol 2010 | GEO: GSE23593 | 50 | Affymetrix U133 Plus 2.0 | 54K | 50 |
| Korde et al.,  Breast Cancer Res Treat 2010 | GEO: GSE18728 | 61 | Affymetrix U133 Plus 2.0 | 54K | 61 |
| Silver et al.,  J Clin Oncol 2010 | GEO: GSE18864 | 84 | Affymetrix U133 Plus 2.0 | 54K | 84 |
| Jonsson et al.,  BCR 2010 | GEO: GSE22133 | 359 | Swegene H_v2.1.1 55K | 55K | 343 |
| Chen et al.,  Breast Cancer Res Treat 2010 | GEO: GSE10780 | 185 | Affymetrix U133 Plus 2.0 | 54K | 42 |
| Desmedt et al.,  J Clin Oncol 2011 | GEO: GSE16446 | 120 | Affymetrix U133 Plus 2.0 | 54K | 120 |
| Guedj et al.,  Oncogene 2011 | Array Express: E-MTAB-365 | 537 | Affymetrix U133 Plus 2.0 | 54K | 452 |
| TCGA,  Nature 2012 | TCGA Data Portal - BRCA - | 1215 | Illumina, RNAseq V2 | 20K | 1,092 |
| Ellis et al.,  Nature 2012 | GEO: GSE29442, GSE35186 | 201 | Agilent-014850 4x44K | 44K | 201 |
| Curtis et al.,  Nature 2012 | EGA: EGAS00000000083 | 2136 | Illumina HT 12 | 49K | 1974 |
| Sabatier R et al., (our IPC series) PLoS One 2011 | GEO: GSE31448 | 353 | Affymetrix U133 Plus 2.0 | 54K | 286 |
| TOTAL | | 6,515 |  |  | 5,883 |
