## Supplementray Table3 for "The serine/threonine kinase MINK1 directly regulates the function of promigratory proteins"

**Supplementary Table 3:** Clinicopathological characteristics of samples in the whole data set

| **Characteristics** | | **N (%)** |
| --- | --- | --- |
| Age at diagnosis (years) | |  |
|  | <=50 | 1363 (28%) |
|  | >50 | 3564 (72%) |
| Pathological type | |  |
|  | ductal | 3520 (79%) |
|  | lobular | 477 (11%) |
|  | other | 483 (11%) |
| Pathological tumor size (pT) | |  |
|  | pT1 | 1512 (36%) |
|  | pT2 | 2214 (53%) |
|  | pT3 | 445 (11%) |
| Pathological axillary lymph node status (pN) | | |
|  | negative | 2117 (49%) |
|  | positive | 2233 (51%) |
| Pathological grade | |  |
|  | 1-2 | 2213 (53%) |
|  | 3 | 1936 (47%) |
| Molecular subtype | |  |
|  | HR+/HER2- | 3941 (67%) |
|  | HER2+ | 728 (12%) |
|  | TN | 1197 (20%) |
| Metastatic relapse | |  |
|  | no | 16329 (79%) |
|  | yes | 424 (21%) |
| Follow-up, months | median (range) | 42 (1-232) |
| 5-year MFS | % [95%CI] | 71% [69-74] |
